# Phantom genetic nurture: assortative mating accounts for most of the apparent association between parental genotypes and childhood cognitive performance

**DOI:** 10.64898/2026.09.01.748516

**Authors:** Daniel S. Malawsky, Laura Hegemann, Olivia Wootton, Alexandra Karoline Havdahl, Hans Fredrik Sunde, Hilary C. Martin

## Abstract

Parental genotypes may influence offspring outcomes through the environments parents provide, a process known as genetic nurture, but estimating such effects from polygenic scores is complicated jointly by measurement error, unobserved genetic variation, and assortative mating. Here, we introduce RAVEL, a structural equation modeling framework that uses two independently constructed polygenic scores for the same trait to estimate latent genetic liability associations in parent-offspring trios and to distinguish direct and non-transmitted genetic associations. Through analytic derivations and simulations, we show that naive trio regressions can produce spurious non-zero parental coefficients, attenuate genuine genetic nurture effects, and obscure asymmetric parental effects, whereas RAVEL recovers unbiased estimates of the underlying coefficients when the assortative mating history and the extent of unobserved genetic variation are correctly specified. Applying RAVEL to national test scores in the Norwegian Mother, Father and Child Cohort Study using polygenic scores for educational attainment, we find that parental genetic liabilities explain less than 0.5% as much variance in childhood cognitive performance as the child’s own genetic liability (substantially lower than the 10% observed with naive trio regressions) once assortative mating is modeled, with only a small paternal association significantly surviving the correction. In contrast, we find a maternal-specific non-transmitted association with offspring premature birth. RAVEL provides a general framework for interpreting trio polygenic score analyses of direct and non-transmitted genetic effects, clarifying the contribution of parental genetic liabilities to offspring traits.

## Introduction

Whether parental genotypes causally affect offspring traits via influence on their rearing environments (‘genetic nurture’) is a central inferential problem in human genetics.^1,2^ Several methods leveraging pedigree and molecular data have been developed to estimate statistics related to this potential role of parental genetics. The classical twin variance decomposition (ACE model) and its more intricate descendants have shown that many complex traits have a significant contribution from the environment shared between twins and siblings.^3–9^ While not necessarily implying genetic nurture effects, shared environmental effects are suggestive of it. However, simplifying assumptions in earlier studies may have led to overestimation of the role of shared environment, as recent work with more relaxed assumptions suggests environmental influence may be smaller than first thought.^4,10^

A complementary suite of methods has been developed that leverages molecular genetic data to estimate the relative contribution of non-transmitted genotypes to offspring traits. Variance decomposition methods including full-sibling IBD regression, relatedness disequilibrium regression, and trio-GCTA leverage realized genetic relatedness (i.e., molecular genetic estimates of kinship between individuals) to estimate the relative contribution of offspring and parental genetics to the variance of a given trait (Table 1).^11–14^ A related and distinct approach has been to regress the phenotype on the individual’s polygenic score (PGS) for a certain trait, conditioning on the corresponding parental values (trio model).^1,2,15^ The resulting associations decompose the regression estimates from standard focal-individual PGS associations into the direct genetic effects of the offspring genotypes on the trait (DGEs) and the residual association tagged by the parental non-transmitted alleles (non-transmitted coefficients; NTCs). Most recently, JODIE extends this family of approaches by jointly modelling direct, maternal, paternal and parent-of-origin effects genome-wide in phased trios.^16^ The relatedness-based and PGS methods have been used widely in the field, particularly trio PGS regressions, and interpretations of NTCs as evidence of genetic nurture are pervasive.^17–25^

**Table 1.**
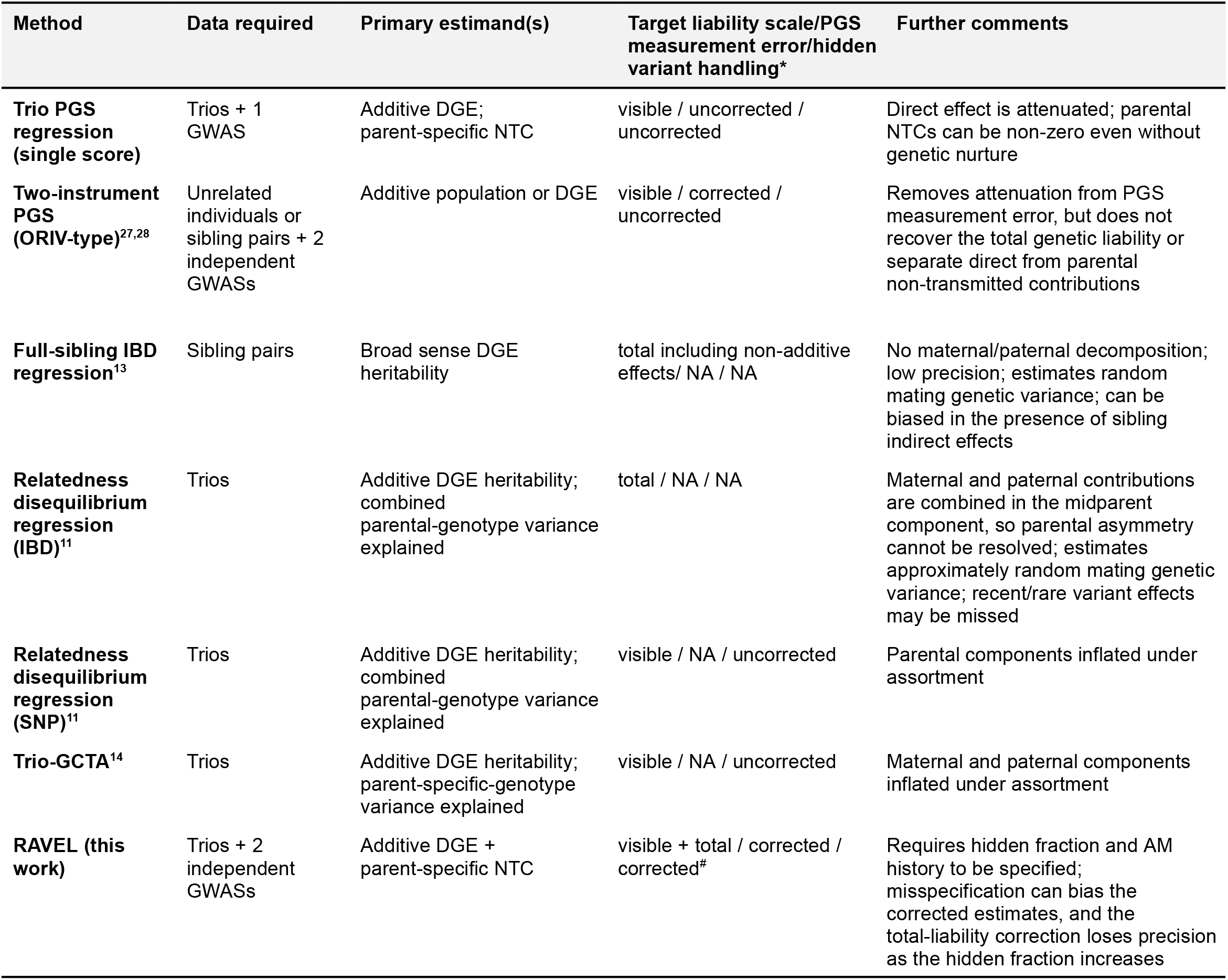
Molecular approaches to recovering DGEs and NTCs leveraging PGSs or genetic relatedness, and the biases each is subject to. ORIV, obviously related instrumental variables. * Hidden variants are those variants relevant to trait heritability that are not assayed. #The hidden fraction corrections are conditional on correctly specifying two structural parameters (Supplementary Note Sections 2 and 6; sensitivity analyses in Supplementary Table 5).

| Method | Data required | Primary estimand(s) | Target liability scale/PGS measurement error/hidden variant handling* | Further comments |
| --- | --- | --- | --- | --- |
| **Trio PGS regression (single score)** | Trios + 1 GWAS | Additive DGE; parent-specific NTC | visible / uncorrected / uncorrected | Direct effect is attenuated; parental NTCs can be non-zero even without genetic nurture |
| **Two-instrument PGS (ORIV-type)<sup>27,28</sup>** | Unrelated individuals or sibling pairs + 2 independent GWASs | Additive population or DGE | visible / corrected / uncorrected | Removes attenuation from PGS measurement error, but does not recover the total genetic liability or separate direct from parental non-transmitted contributions |
| **Full-sibling IBD regression<sup>13</sup>** | Sibling pairs | Broad sense DGE heritability | total including non-additive effects/ NA / NA | No maternal/paternal decomposition; low precision; estimates random mating genetic variance; can be biased in the presence of sibling indirect effects |
| **Relatedness disequilibrium regression (IBD)<sup>11</sup>** | Trios | Additive DGE heritability; combined parental-genotype variance explained | total / NA / NA | Maternal and paternal contributions are combined in the midparent component, so parental asymmetry cannot be resolved; estimates approximately random mating genetic variance; recent/rare variant effects may be missed |
| **Relatedness disequilibrium regression (SNP)<sup>11</sup>** | Trios | Additive DGE heritability; combined parental-genotype variance explained | visible / NA / uncorrected | Parental components inflated under assortment |
| **Trio-GCTA<sup>14</sup>** | Trios | Additive DGE heritability; parent-specific-genotype variance explained | visible / NA / uncorrected | Maternal and paternal components inflated under assortment |
| **RAVEL (this work)** | Trios + 2 independent GWASs | Additive DGE + parent-specific NTC | visible + total / corrected / corrected<sup>#</sup> | Requires hidden fraction and AM history to be specified; misspecification can bias the corrected estimates, and the total-liability correction loses precision as the hidden fraction increases |

However, trio PGS analyses currently face several limitations. First, PGSs are a noisy estimate of the effects of the variants they include, and they do not capture variance attributable to variants they do not include or tag. Both the reliability (i.e. 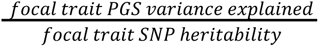) and the fraction of the trait’s heritability captured by the SNP heritability are not typically known. Under random mating, both features merely attenuate the direct effect estimate, but, as we discuss below, they interact with assortative mating (AM) to distort the non-transmitted coefficients in complex ways.^26^ The genetic relatedness-based variance decomposition methods sidestep the first problem entirely, so in principle they reflect the full additive genetic contribution; however, they forgo several advantages discussed below and, like the trio regressions, many remain confounded by AM (Table 1). Neither family of methods, as currently applied, recover interpretable estimates of parental genetic contributions to variation in offspring traits.

Despite their interpretive limitations, trio model PGS analyses have three attractive features relative to the genetic relatedness-based variance decomposition methods. First, PGSs for any trait, including those that may be reasonable candidates for indexing genetic nurture effects, can be studied in the context of any offspring outcome (for example, assessing the role of genetic liability for educational attainment (EA) on offspring cognitive development). Second, they can leverage estimates of genetic effects from large-scale GWASs such that PGS associations can be precisely estimated in relatively small cohorts. Third, each family member’s score indexes the same genetic liability, which need not be that of the outcome, so a non-transmitted coefficient reflects correlations with a specified genetic liability. Genetic relatedness-based methods partition variance regardless of whether the liability is shared or distinct between family members, and so, for example, cannot distinguish a maternal contribution acting through the same liability underlying the direct effect from one acting through an uncorrelated liability (e.g. the maternal contribution to her child’s cognitive performance may not act through the direct genetic effects on her own childhood cognition, but rather other traits). In principle, this feature of the trio PGS models gives one a place to start when looking for potential rearing environment moderators of the genetic nurture effect.

To address these limitations, we developed **RAVEL** (**R**ecovering **A**ssortment-adjusted **V**ariance **E**xplained by genetic **L**iabilities), a structural equation modeling framework that leverages PGSs constructed using two independent GWASs to estimate the full (disattenuated) variance explained by both the offspring genetic liability indexed by the PGSs and the variance explained by the parental genetic liabilities after correcting for AM. Through simulations, we show that RAVEL can accurately recover parameter estimates across all tested settings. We then show several empirical examples in which RAVEL leads to qualitatively different conclusions from naive trio analyses. Our work demonstrates that previously reported potential genetic nurture effects have likely been upward biased by AM, and provides a method to robustly correct for this source of confounding.

## Results

### Trio PGS regression yields biased estimates of genetic nurture under assortative mating

Trio PGS regression is a widely used approach for decomposing offspring genotype-phenotype associations into DGEs and NTCs,^17–25,29^ but two phenomena bias the NTC from estimating genetic nurture: polygenic scores are noisy estimators of the underlying genetic liability, and they include or tag only a subset of the variants contributing to a trait’s genetic liability. To quantify how each source contributes to the bias under AM, and to determine the extent to which removing PGS measurement error alone would suffice to recover unbiased coefficients (referred to here as the visible-scale liability; see Box 1), we analytically derived the expected coefficients for a noisy PGS and the visible-scale liability under three AM scenarios and two genetic architectures (Figure 1). We estimate these coefficients assuming the noisy PGS explains 50% of the variance that is explained by the visible-scale liability (i.e. PGS reliability = 0.50), and by varying the fraction of total heritability not explained by the visible-scale liability under random mating (the hidden fraction, π). (With a standard dense, genome-wide PGS, the visible-scale liability should recover what is commonly called the SNP heritability, h^2^ when regressed on the same phenotype as PGS trait).

**Figure 1.**
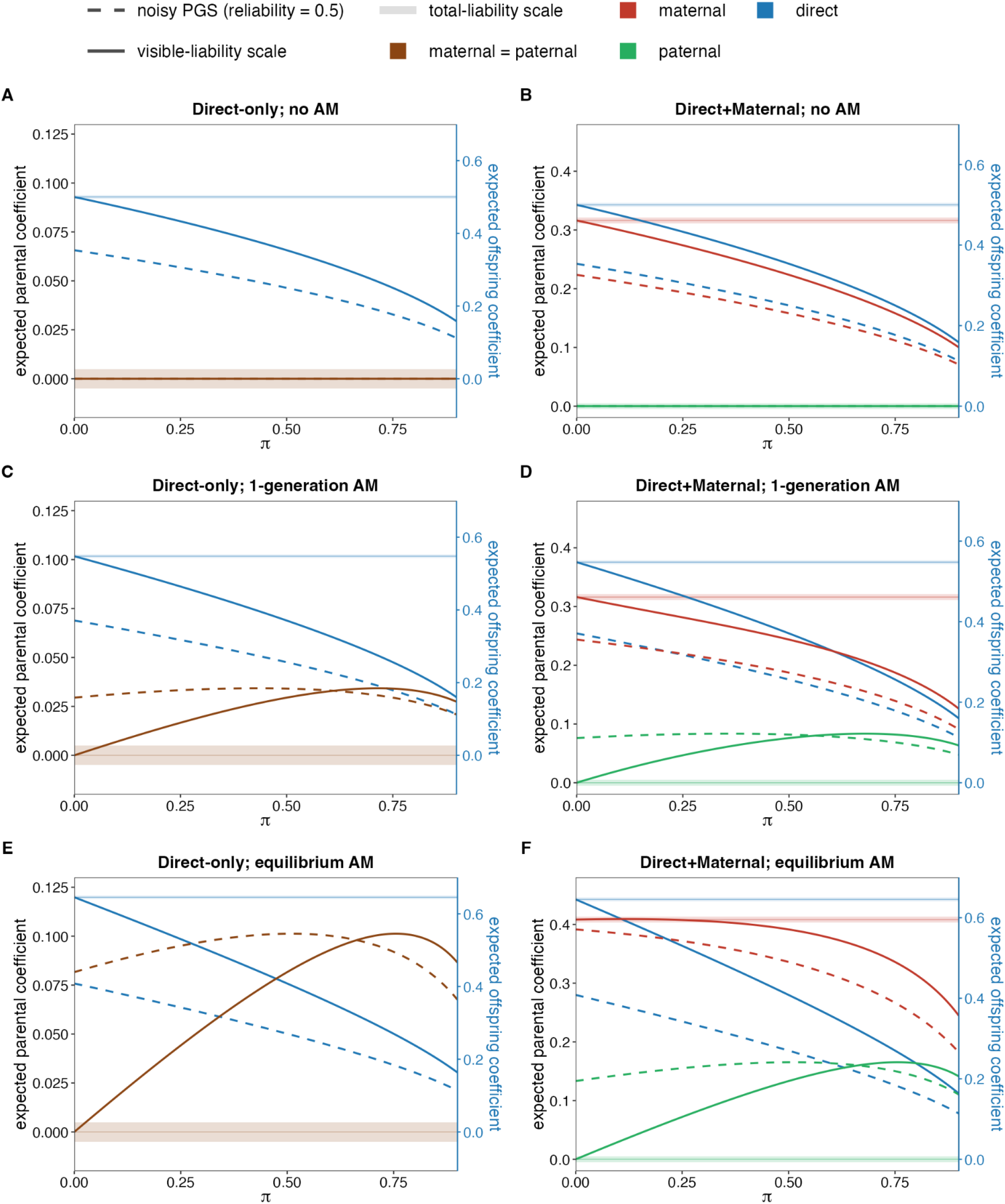
Expected coefficients in trio PGS analyses. Expected standardized DGEs and parental NTCs from trio PGS regression as a function of the hidden fraction, π, for two architectures and three AM regimes. Dashed lines indicate the coefficient for a noisy PGS (reliability = 0.50 under random mating), solid for a hypothetically noise-free PGS, i.e., the visible-scale liability (reliability = 1). Line colour indicates the coefficient plotted: brown for the parental NTC (expected to be equal if there are no NTCs absent AM) in the direct-only architecture (β_D_ = 0.5, β_M_ = β_P_ = 0); red for the maternal NTC in the direct+maternal architecture (β_D_ = 0.5, β_M_ = 0.32, β_P_ = 0); green for the paternal NTC in the direct+maternal architecture; blue for the offspring DGE expectations. Horizontal shaded bands give the true total-liability value of each DGE and NTC (the causal target), drawn in the same colour as the corresponding lines. Panels: (A,B) no AM; (C,D) one generation of AM; (E,F) equilibrium AM. Genotypic mate correlation ρ = 0.4 in the AM panels.

The two genetic architectures we considered were one with only a DGE (direct-only: [β_D_ = 0.5, β_M_ = β_P_ = 0]) and one with both a DGE and a maternal genetic nurture effect only (direct+maternal: [β_D_ = 0.5, β_M_ = 0.32, β_P_ = 0]) and three AM regimes (no AM, one generation of AM, equilibrium AM, the latter two with genotypic mate correlation ρ = 0.4). In the absence of AM, both the noisy PGS and visible-scale liability recover qualitatively correct DGEs and NTCs across different values of the hidden fraction i.e. the NTC is estimated at 0 under the direct-only architecture, whereas under the direct+maternal architecture, the maternal NTC estimate is positive (although attenuated below the true value on the visible-scale unless the hidden fraction is 0) and the paternal NTC is correctly estimated at 0 (Figure 1A,B). PGS reliability below 1 and non-zero hidden fraction alone therefore produce attenuations but do not introduce spurious non-zero NTCs; AM is the source of the qualitative biases that follow.

##### Box 1. Recurring terminology.

##### Terminology

###### Assortative mating (AM)

Non-random mating in which individuals are more likely to choose partners with (dis)similar values of a trait to themselves.

###### Equilibrium assortative mating (equilibrium AM)

The limiting state reached after assortative mating has operated for enough generations that the genetic liability (co)variance structure no longer changes appreciably from one generation to the next.

###### Genetic liability

An individual’s additive genetic propensity toward a trait represented as a latent quantitative variable.

###### Total-scale liability (T)

The genetic liability that indexes all additive genetic variation relevant to a given trait. In a regression of a given additive trait on its total-scale liability, the variance explained will be equivalent to the full heritability of the trait.

###### Visible-scale liability (G)

The component of the total-scale genetic liability indexed by the measured variants used to construct the PGSs (typically genome-wide common SNPs). It is the latent genetic factor identified by the two observed PGSs in RAVEL. In a regression of a given additive trait on its own visible-scale liability, the variance explained will be equivalent to the SNP heritability of the trait.

However, this is not the case when the phenotype underlying the genetic liability and the focal regression trait differ.

###### Hidden fraction (π)

The proportion of total-scale liability variance that is *not* captured by the visible-scale genetic liability under random mating. Equivalently, (1-π) is the fraction of total additive genetic variance captured by the visible component under random mating.

###### Direct genetic effect (DGE)

The association of the offspring’s genetic liability with their own phenotype, conditional on the corresponding maternal and paternal genetic liabilities.

###### Parental non-transmitted coefficient

The association between parental genetic liabilities and the offspring phenotype after conditioning on the offspring’s and the other parent’s genetic liabilities. After correction to the total-liability scale, it represents the association attributable to parental liabilities that are not transmitted to the child.

###### Genotypic mate correlation (ρ)

The correlation between the genetic liabilities of mating partners for the trait under study. This is distinct from the phenotypic correlation between mates.

###### PGS measurement error

Error arising because GWAS-estimated variant weights imperfectly measure the effects underlying the genetic liability.

In the direct-only architecture under AM, both estimands yield non-zero DGEs but also non-zero parental NTCs (Figure 1C,E), confirming that AM alone can produce a spurious NTC in the absence of any genetic nurture. When the visible-scale liability coincides with the total-scale genetic liability, it correctly recovers a parental NTC of zero, but only with perfect reliability: the noisy PGS still yields a substantial spurious NTC. Removing PGS measurement error alone therefore eliminates the spurious NTC only in the special case where the visible-scale liability captures the total-scale liability, which is unlikely in most settings. Equilibrium AM increases the spurious NTCs relative to one generation of AM, reflecting the multi-generational accumulation of within-person covariance between the variants captured by the PGS and those not included.

In the direct+maternal architecture under AM, both the PGS and visible-scale estimands attenuate the genuine maternal NTC and inflate the paternal NTC toward a spurious non-zero value (Figure 1D,F). The maternal NTC from the visible-scale liability is slightly upward biased at low hidden fractions under equilibrium AM (Figure 1F) before declining, while the maternal NTC from the noisy PGS decreases nearly monotonically across the range of π in both AM regimes. The paternal NTC follows a qualitatively similar trajectory to the parental NTC in the direct-only setting. At high hidden fractions (π > 0.8) the spurious paternal NTC approaches the attenuated maternal NTC in magnitude under both estimators, such that a naive trio PGS regression under these conditions would infer approximately equal maternal and paternal contributions when only the maternal effect is causal (Figure 1D,F).

Two implications motivate the method developed below. First, removing PGS measurement error alone is not sufficient to obtain NTC estimates unbiased by AM. Second, no fixed multiplicative scalar correction maps the visible-scale liability trio coefficients onto the desired total-liability coefficients. Under AM, such a regression still yields a residual bias unless all variants influencing the trait are included or fully tagged by the score.

### Recovering assortment-adjusted variance explained by genetic liabilities

We developed RAVEL as a structural equation model to estimate the association between latent total genetic liabilities of parents and offspring and an outcome. Suppose that each individual in the trio with a role, r (i.e. mother, father, or offspring), has a standardized additive genetic liability *G_r_* towards a given trait, as indexed by a set of observed genetic variants (Figure 2). Note that the GWAS used to construct the PGS need not be of the same trait as the outcome (i.e., one can study the effect of genetic liability for a given trait e.g., EA, on an arbitrary phenotype of interest, e.g. cognitive performance). Rather than observing *G_r_*, one observes two standardized polygenic scores *P*_r_ ^(1)^ and *P*_r_ ^(2)^, each constructed from independent discovery GWASs such that their estimation errors are independent. Each score covaries with *G_r_* with a correlation λ_1_ and λ_2_ and score-specific residual ε^(k)^:

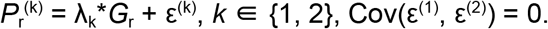

**Figure 2.**
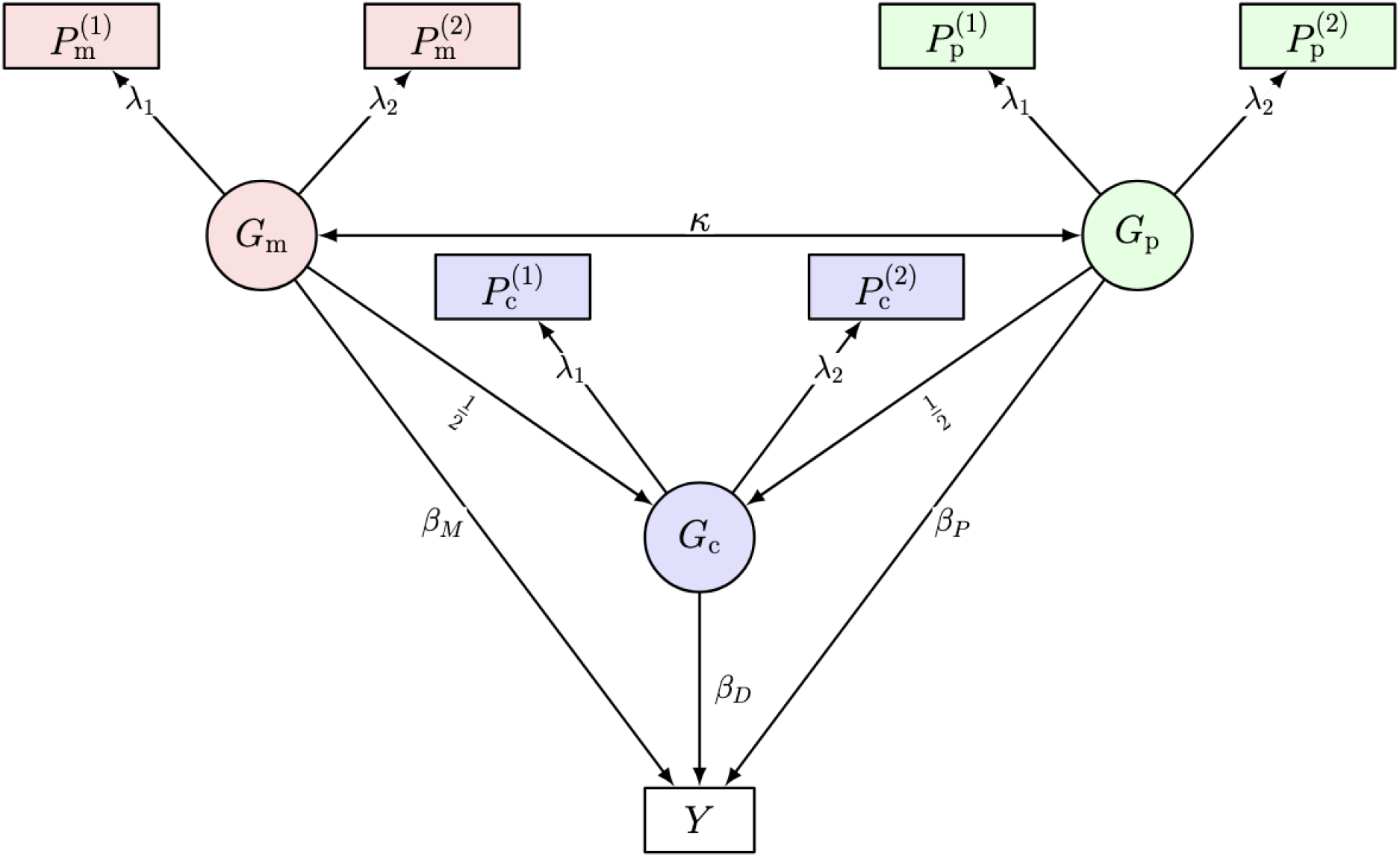
Conceptual path diagram for the RAVEL trio model. Each family member has an additive genetic liability *G*_r_, r ∈ {m (mother), p (father), c (child)} indexed by two independently constructed polygenic scores, P_r_^(1)^ and P ^(2)^. AM is represented by the covariance between *G* and *G* , denoted κ. The offspring liability *G*_c_ is generated from the parental liabilities, and the offspring phenotype Y is regressed on *G*_c_, *G*_m_, and *G*_p_ with coefficients *β_D_*, *β_M_*, and *β_P_*. Residual covariances between relatives’ polygenic scores are omitted to reduce clutter.

In words, the two scores are noisy, independent proxies of an underlying genetic liability tagged by the variants included in the PGSs. The covariance of the scores therefore identifies the product,

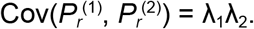

Thus, with two independently constructed scores per individual, the product of the loading of each score on the observed component of the genetic liability is identifiable directly. This two-instrument logic has been used in single-individual and sib-difference PGS regression settings previously.^27,28,30^

The above formulation now recovers the associations with the visible-scale liabilities of the trio members, but this is not typically the target regression of interest. The RAVEL SEM estimates only the visible-scale model fit. To recover the regression of the total-scale genetic liabilities on the outcome, one has to apply a post-estimation correction that further encodes the relationship between the visible-scale estimates and those of the total genetic liability towards the trait, *T*, which includes variants not included or tagged by the PGSs, that nonetheless contribute to the heritability of the trait (Figure S1, Supplementary Note Section 5). To do so, one must specify i) the fraction of total heritability not explained by the visible-scale liability under random mating, π and ii) the effective number of generations of constant-strength AM acting on the trait. While these are not identifiable from the RAVEL framework itself, they can be directly or indirectly inferred with minimal bias (Supplementary Note Section 2).^4,31^ Estimating the regression on total-scale liabilities then allows for an intuitive interpretation of the estimated NTCs, as they no longer reflect potential assortment-induced tagging.

### Simulations

To assess whether RAVEL recovers unbiased estimates of the DGEs and NTCs on both the visible (*G*) and total (*T*) liability scales, we evaluated its performance in two trio simulation frameworks. The first is a single generation simulator that draws causal variant effects, matches parental haplotypes to achieve a target AM strength, and constructs trio offspring by Mendelian transmission; polygenic scores are generated by adding per variant estimation noise to a fraction of the causal variants, with the remaining fraction left “hidden” and contributing a fixed share, π, of the total genetic variance under random mating (see Methods and Supplementary Note Section 10). The second is the multigenerational forward-time simulator GeneEvolve^32,33^, modified to expose the haplotype matrices necessary for trio polygenic score calculation, in which the population evolves for 15 generations under a specified AM and indirect effect scenario, which is approximately equilibrium AM.^31^ Simulations used 16,000 trios per replicate with 200 replicates per parameter combination across five and three genetic architectures in the one-generation and 15-generation settings (three DGE-only architectures spanning β_D_ of {0, 0.5, 0.71}; one asymmetric “direct+maternal” architecture with β_D_ = 0.5 and β_M_ = .32, β_P_ = 0.0; one symmetric “both parents” architecture with β_D_ = 0.45 and β_M_ = β_P_ = 0.22), three AM strengths (ρ ∈ {0, 0.2, 0.4}), and two hidden variant fractions (π ∈ {0.4, 0.8}) with the two PGS reliabilities calibrated to 0.3 and 0.5. Per-parameter combination estimates and ground truth values are reported in Supplementary Tables 1 and 2, demonstrating unbiased recovery of the true parameters across all tested scenarios. We discuss a few examples below.

As expected, when the data generating process has no parental indirect effects, visible-scale RAVEL recovers a DGE coefficient that is attenuated relative to the total-scale truth by an amount determined primarily by the hidden fraction (Figure 3A). Under the same DGE-only simulations, visible-scale RAVEL recovers non-zero parental coefficients whose magnitudes scale jointly with AM and the hidden fraction and are symmetric between the maternal and paternal sides up to sampling error (Figure 3B). Specifying the hidden fraction and the AM history within RAVEL (‘total-scale RAVEL’) recovers the total liability-scale truth across the four (ρ, π) combinations and returns β_M_ and β_P_ to 95% empirical intervals centered at zero (Figure 3A,B, Supplementary Table 1). The width of the total-liability 95% intervals was governed principally by the hidden fraction. At a hidden fraction of 0.4, the 95% empirical interval for each parental coefficient spanned approximately ±0.035 at 16,000 trios, versus ±0.024 on the visible scale; at a hidden fraction of 0.8, where the scores index only a fifth of the additive genetic variance, the same interval widened to approximately ±0.074. The correction therefore trades precision for interpretability in proportion to how little of the genetic variance the scores capture. Misspecifying the hidden fraction and/or the AM history as being too high led to overcorrected, biased estimates along the same gradients as those going from the visible-scale to total-scale correction, e.g., overcorrection in the direct-only scenario led to overestimation of DGEs and negative NTC estimates (Supplementary Table 1). We then considered RAVEL’s performance when correcting for 15 generations of assortment (Figure 3C). RAVEL was able to recover unbiased estimates of the visible and total liability coefficients provided the correct AM history.

**Figure 3.**
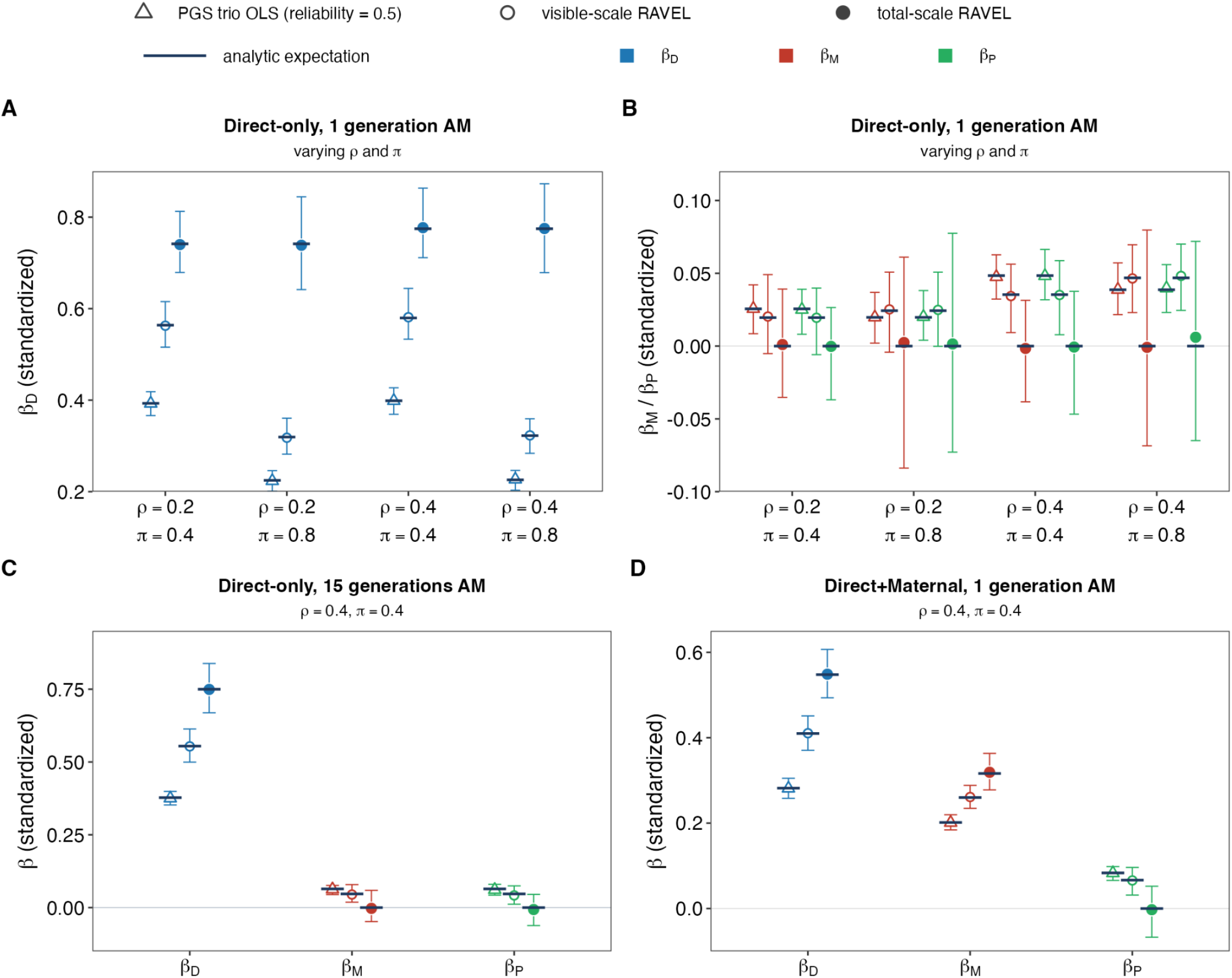
Simulation validation of RAVEL. Each panel reports point estimates with 95% empirical intervals across simulation replicates of N = 16,000 trios. Triangles are the estimate from a noisy PGS (reliability=0.5 under random mating), open circles are standardized visible-scale RAVEL, and filled circles are standardized total-scale RAVEL estimates, fit with π and the AM history specified to match the data-generating process. Two PGSs were constructed to have reliabilities of 0.5 and 0.3, respectively, in all simulations. Solid horizontal black bars indicate analytic expectation. (A) Recovery of the visible and total β_D_ across four (ρ, π) parameter combinations with one generation of AM with a DGE-only architecture (β_D_ = 0.71, β_M_ = 0, β_P_ = 0). (B) Visible and total scale parental coefficients from the same direct-only simulations as in (A). Visible-scale RAVEL recovers non-zero values for β_M_ (red) and β_P_ (green) whose magnitudes scale jointly with ρ and π; specifying π and the AM history returns unbiased estimates of β_M_ and β_P_. (C) Recovery of the visible and total effects under 15 generations of assortment at ρ = 0.4 and π = 0.4 with a DGE-only architecture (β_D_ = 0.71, β_M_ = 0, β_P_ = 0). (D) Recovery of the visible and total effects under one generation of asymmetric direct+maternal architecture (β_D_ = 0.50, β_M_ = 0.32, β_P_ = 0) at ρ = 0.4 and π = 0.4. Visible-scale RAVEL recovers an attenuated maternal coefficient and a non-zero paternal coefficient.

We also considered a more complex direct+maternal architecture, in which the data-generating process contains a real maternal effect but no paternal effect, in a population with one generation of strong assortment (ρ = 0.4) and substantial hidden heritability (hidden fraction of 0.4) (Figure 3D). RAVEL recovered the visible and total-scale coefficients unbiasedly (Figure 3D). Together, these results show that RAVEL recovers the visible and the total-liability coefficients without bias whenever the assumed hidden fraction and AM history are correctly specified.

We next assessed the precision of RAVEL estimates as a function of the number of observed complete trios. We repeated the direct+maternal simulation 200 times across a range of trio sample sizes, using two PGSs calibrated to have reliabilities of 0.30 and 0.50 as before (Figure S2). These values were chosen to represent realistic levels of PGS measurement error: For a trait with a heritability of 0.5 that is highly polygenic, reliabilities of these magnitudes can be achieved with discovery GWAS sample sizes on the order of 10^5^ individuals.^34^ As expected, the standard errors of the RAVEL coefficients declined approximately linearly with N^-1/2^ as the number of trios increased (Figure S2). The largest gains in precision occurred between 4,000 and 16,000 trios, with further but progressively smaller improvements at 32,000 and 64,000 trios. Across the sample size range, the direct, maternal, and paternal coefficients showed similar magnitude and scaling of uncertainty, indicating that the parental non-transmitted coefficients are not intrinsically less precisely estimated than the offspring direct effect once the latent genetic liability is identified by two independent PGSs. These simulations suggest that RAVEL can be informative in trio cohorts of the size currently available (e.g. FinnGen, Estonian Biobank, MoBa and deCODE all have >10,000 trios^35^).

### RAVEL reveals qualitatively different patterns of associations from trio regressions

We then applied RAVEL to study the association between polygenic scores for EA and childhood cognitive performance using data from the MoBa cohort. We operationalized cognitive performance using the students’ scores in the national tests that are taken in Norway in grades 5 and 8 (ages ∼10 and ∼13, respectively).^36^ These tests assess basic skills in reading (Norwegian), mathematics (numeracy), and a foreign language (English), sampling a broad range of cognitive processes. They are standardized and administered under similar conditions across schools and regions, and unlike school grades, do not have direct consequences for students’ further schooling and are therefore less affected by strategic grading practices or high-stakes pressures, and are not subject to individual teacher bias. We took the average of the tests to produce overall grade 5 and grade 8 scores. We constructed two EA PGSs using 23andMe Research Institute as one discovery cohort (n=2,713,033), and the EA4 GWAS meta-analysis excluding 23andMe (n=765,283) as the other cohort.^37^

For each outcome we report the offspring DGEs and the parental NTCs using three approaches: standard trio regression on the PGSs (one for each of the two scores), visible-scale RAVEL, and total-scale RAVEL assuming a hidden fraction 0.46 and three generations of AM as has been previously estimated.^31^ The assumed hidden fraction was chosen as it recovered the parental total-liability genotypic spousal correlation consistently (e.g. in the grade 8 test analysis ρ = 0.343 [95% CI 0.320 - 0.365]), matching contemporary Norwegian estimates of the EA genotypic liability spousal correlation based on a two-generation pedigree-based method (iAM-COTS).^4^

The continuous test scores display the bias structure that the simulations and analytical derivations predict for a trait predominantly influenced by DGEs (Figure 4A, Supplementary Table 3-4). For grade 8, single-PGS trio regressions yielded standardized β_D_ estimates of 0.276 (PGS1 based on Okbay et al without 23andMe^37^; 95% CI [0.261, 0.291]) and 0.252 (PGS2 based on 23andMe only; [0.237-0.267]); visible-scale RAVEL recovered a β_D_ of 0.361 (0.342-0.381), and the full RAVEL correction further increased β_D_ to 0.529 (0.500-0.558). The maternal coefficient followed the opposite trajectory: the two PGS regression estimates and the visible-scale RAVEL estimate were all positive and highly significant, but the fully corrected estimate decreased substantially and was no longer significant. The paternal coefficient attenuated but retained a positive total-liability estimate of 0.030 (95% CI [0.014 - 0.046]). Comparing the PGS coefficients of the maternal and paternal effects from the naive trio regressions (and disregarding the covariance between family members’ genetic liabilities), one would infer the maternal and paternal NTCs together account for over a tenth as much phenotypic variance as the direct genetic effect, whereas RAVEL implies that they explain less than 0.5% as much. We carried out sensitivity analyses by varying the assumed number of generations between 1 to 15 and the hidden fraction parameter from 0.4 to 0.5 (spanning an implied genotypic mate correlation of 0.317 to 0.361 assuming 3 generations of AM), which did not qualitatively change the results (Figure S3, Supplementary Table 5).

**Figure 4.**
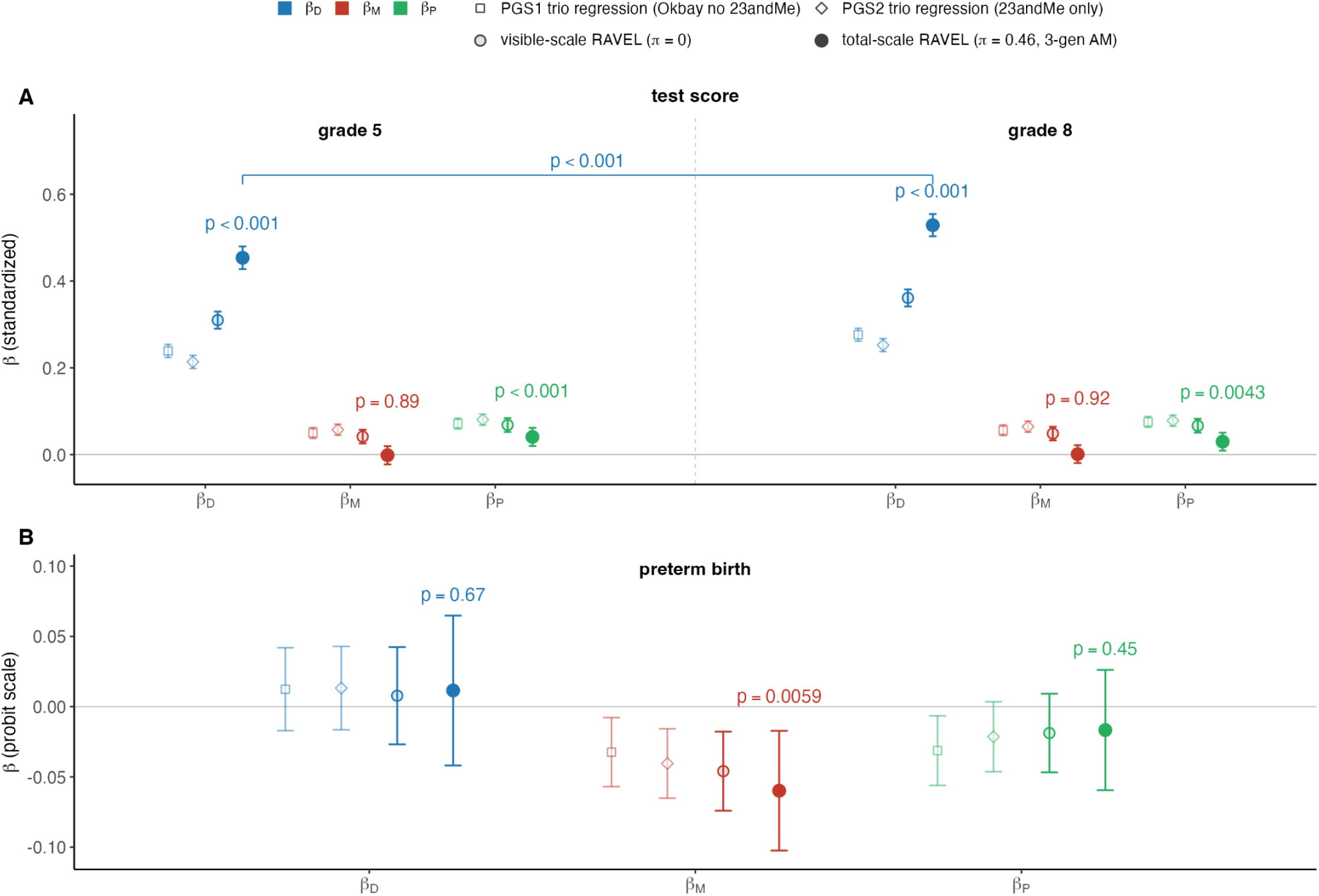
RAVEL applied to test scores and preterm birth in MoBa. Each panel reports point estimates with 95% confidence intervals from four estimators applied to complete genotyped MoBa trios. Open squares are PGS1 trio OLS estimates (PGS1 constructed from the Okbay et al. 2022 educational attainment GWAS with 23andMe excluded) and open diamonds are PGS2 trio OLS estimates (PGS2 constructed from the 23andMe-only EA GWAS). Open circles are visible-scale RAVEL estimates fit with no correction (π = 0), and filled circles are total-liability RAVEL estimates corrected with π = 0.46 and three generations of assortment. Coefficients are color-coded: blue for the offspring direct genetic effect β_D_, red for the maternal non-transmitted coefficient β_M_, and green for the paternal non-transmitted coefficient β_P_. Continuous score estimates are reported on the standardized phenotype scale; binary outcomes are reported on the probit scale. (A) Continuous average test scores at grade 5 (left) and grade 8 (right) (n trios=34,295). (B) Premature birth, defined as gestational age ≤ 37 completed weeks (n trios=38,052). P values displayed for the total-liability RAVEL estimates. Square brackets indicate tests for difference in effect sizes estimated via bootstrap (Methods).

The grade 5 results followed the same qualitative pattern as grade 8, but with slightly smaller magnitudes across estimates (Figure 4). We previously observed that the DGE of the EA PGS on cognitive performance increased across development in a British birth cohort, ALSPAC.^38^ To test for a similar pattern here, we fit RAVEL to the within-person difference in standardized test scores between grades 8 and 5. Consistent with our ALSPAC findings, the child’s total genetic liability was significantly associated with an increase in relative test performance with age (β_D_ = 0.0743, [0.0570 - 0.0916]), whereas neither maternal nor paternal total genetic liability showed evidence of association with change (β_M_ = 0.0024, [-0.0112 - 0.0160]; β_P_ = -0.0102, [-0.0239 - 0.0034]).

We previously observed that the association between cognition-related PGSs and IQ is strongest at the top of the IQ distribution, and weakest at the bottom of the distribution.^38^ We thus considered whether association estimates differed at the extremes of the phenotypic distribution. At both grades 5 and 8, we observe a larger DGE at the top 10th relative to the top 90th percentile (bootstrap p value for DGE at top 10% > top 90% = 0.0025 grade 5, 0.0017 grade 8), consistent with the prior observation (Figure S4). The estimates attained for binarized test scores at the top 10th percentile yielded similar results to the continuous phenotype. The total-liability RAVEL maternal NTCs were not significant. Only the corrected paternal coefficient was significant after multiple testing (0.05/4) at the top 10% at grade 8 and nominally significant at the top 10% at grade 5.

Premature birth is associated with poorer cognitive outcomes^38–42^, and lower maternal EA is associated with increased risk of premature delivery.^43^ Consistent with this, we previously observed that, in a cohort of children with severe neurodevelopmental conditions, probands with higher EA PGSs were less likely to have been born prematurely^39^, and that this was driven by significant maternal and paternal NTCs, with no significant direct effect.^44^ Thus, here we were interested in exploring the association between genetic liability towards EA and premature birth. The pattern of associations differed strikingly from that of test scores (Figure 4B, Supplementary Table 3-4). The offspring DGE was statistically indistinguishable from zero, consistent with offspring EA genetic liability not influencing their risk of premature birth.^44^ Both PGS regression estimates of the maternal NTC were negative and significant, and unlike the cognitive outcomes, the maternal NTC was amplified rather than attenuated by both RAVEL corrections. The paternal NTC was no longer significant after the full RAVEL correction.

### Triangulation of the direct genetic effect heritability of average test scores

Next, we wanted to assess concordance between the association estimates above and those obtained by other study designs for assessing the DGE heritability of the average test scores and similar phenotypes. The above analysis estimated β_D_, the standardized DGE coefficient of the total genetic liability towards EA on test scores, Y. We can use this quantity to estimate the DGE heritability of test scores by using the correlation between the genetic liabilities for EA and test scores (i.e. their r_g_) h ^2^ = (β / r (EA , Y))^2^.

We observed a genetic correlation between the average of the test score phenotypes and EA4 of 0.746 (95% CI 0.687-0.805), which implied the DGE heritabilities of the test scores at grades 5 and 8 of 0.369 (0.294-0.444; independent-estimate delta method) and 0.503 (0.406-0.600) (Figure 5A). However, note these heritabilities are likely underestimated since this r_g_ estimate was obtained using population GWASs, which have been previously shown to overestimate the genetic correlation between EA and cognitive ability obtained using within-family GWAS^45^. We then calculated the heritability for the test scores using a large pedigree-based study design including millions of Norwegian relative pairs, estimating grade 5 and 8 narrow-sense heritabilities in line with those estimated by RAVEL (Figure 5A, Methods).^46^ The increase in heritability between these timepoints was similarly significant (difference = 0.0764, bootstrap p value = 0.0008). A separate Norwegian study previously leveraged a children-of-twins-and-siblings pedigree design to estimate the narrow-sense heritability of these same grade 5 and 8 test scores, again showing similar heritability estimates to those estimated here.^47^ Furthermore, we estimate full-sibling IBD regression estimates of heritability of 0.536 (0.173-0.898) and 0.868 (0.456-1.280) for average grade 5 and 8 test scores (Supplementary Table 6), respectively, which are consistent with the full-sibling IBD regression estimate attained in a meta-analysis of cognitive performance measures across several cohorts^48^, though these estimates were imprecise (Figure 5A).

**Figure 5.**
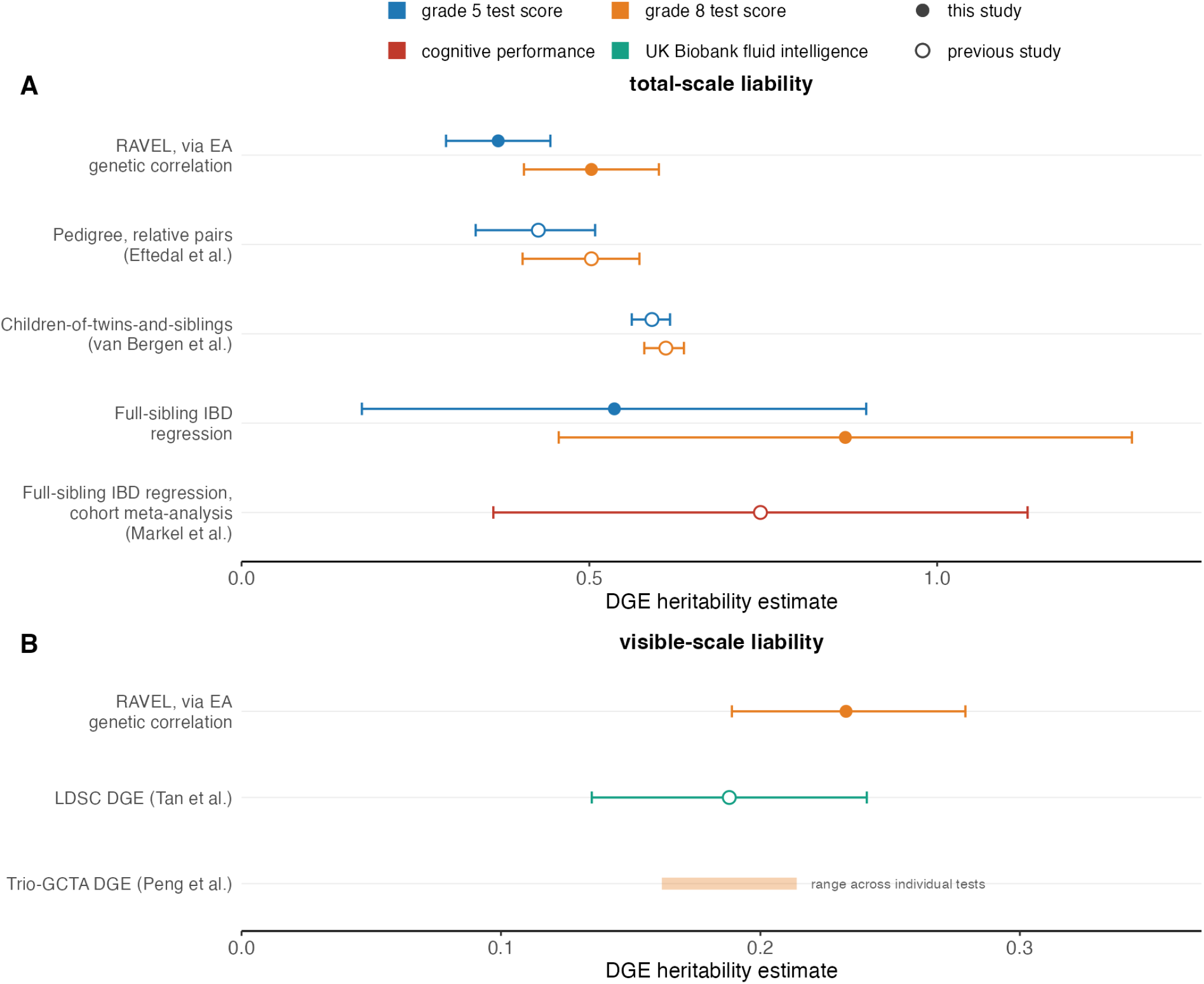
Triangulation of direct genetic effect (DGE) heritability estimates for cognitive performance-related traits. Each panel reports point estimates with 95% confidence intervals from complementary study designs, one estimator per row. Filled circles are estimates from the present study and open circles are estimates reported by previously published studies. Estimates are color-coded by phenotype: blue for the grade 5 test grade, orange for the grade 8 test grade, purple for the average of the test scores, red for cognitive performance, and teal for UK Biobank fluid intelligence. Because the designs shown target slightly different estimands, the horizontal axis is labelled generically as the DGE heritability estimate. (A) Total-scale liability. Rows are, from top, RAVEL rescaled by the EA genetic correlation; a pedigree design comprising millions of Norwegian relative pairs^46^; a children-of-twins-and-siblings design applied to the average grade 5 and 8 test scores^47^; full-sibling IBD regression in MoBa; and full-sibling IBD regression meta-analysed across cohorts of cognitive performance measures^48^. (B) Visible-scale liability. Rows are, from top, the same RAVEL rescaling applied to the transformed grade 8 visible-scale DGE variance explained; the LDSC DGE SNP heritability of fluid intelligence in UK Biobank^45^; and trio-GCTA DGE SNP heritabilities of the individual grade 8 tests^49^. The trio-GCTA entry is drawn as a shaded bar spanning the range of estimates across the individual tests rather than as a point estimate with a confidence interval.

Additional molecular genetic-derived estimates of the DGE SNP heritability have also been estimated for cognitive performance traits, offering an additional triangulation by considering the transformed visible-scale liability DGE variance explained. Applying the same transformation as above to the grade 8 test score yielded an estimate of 0.233 (0.189-0.279) for the DGE SNP heritability (Figure 5B). This was consistent with a previous cognitive ability LDSC DGE SNP heritability estimate for fluid intelligence in UK Biobank^45^, as well as trio-GCTA-estimated DGE SNP heritability of the individual test scores in grade 8 in MoBa (Figure 5B).^49^ Thus, the RAVEL estimates are consistent with several alternative means of estimating the DGE total and SNP heritabilities, pointing towards a convergence of heritability estimates.

## Discussion

We introduced RAVEL to improve the interpretability of trio PGS analyses in the context of AM, which has hitherto presented a challenge to genetic studies of social and behavioural traits. We demonstrated analytically and through simulations that naive trio regressions on PGSs can produce counterintuitive parental associations under AM and when the PGS incompletely tags all relevant genetic variation. We then showed that RAVEL recovers the relevant parameters without bias when the π and the AM history are (approximately) correctly specified. Applying RAVEL in MoBa showed that genetic liability for EA confers substantial DGEs on childhood test scores, whereas the apparent maternal NTC was entirely accounted for by the RAVEL correction and only a small paternal NTC remained, consistent with prior pedigree-based estimates from Norway^47^. In contrast, premature birth showed a distinct pattern, with a persistent maternal NTC and no evidence for an offspring direct effect.

Our analyses of the test scores are consistent with our prior observations in a British birth cohort that the association between EA and cognitive performance PGSs increased with age and that the associations are stronger at the top of the phenotypic distribution.^38^ We observed that the paternal total EA genetic liability was modestly associated with their child being in the top 10 percentile of the cognitive ability distribution, which cannot be explained by residual uncorrected AM and is likely reflective of either paternally mediated genetic nurture, paternal dynastic effects, residual population stratification, or ascertainment biases that are specific to the fathers in complete genotyped trios. Notably, a prior study in MoBa using within family Mendelian randomization similarly found significantly asymmetric, paternal-leaning NTCs on childhood test scores,^29,50^ and we previously observed that paternal, but not maternal, phenotypic EA is positively associated with IQ among children at the top of the IQ phenotypic distribution in a British cohort.^38^ This stands in contrast with interpretations of prior results that found roughly equal effect size estimates for the maternal and paternal EA PGSs in naive trio regressions using the same cohort.^51^ That study also inferred, by leveraging parental siblings, that the majority of the association with parental effects was not due to the alleles specific to the proband’s parent, hypothesizing potentially pervasive ‘dynastic effects’, whereas our analyses suggest this was likely largely a reflection of assortment. More broadly, these results do not imply that children’s environments are unimportant for cognitive performance. Apart from the imprecise full-sibling IBD regression estimates, all total-scale DGE heritability estimates were well below one (Figure 5), leaving substantial variance in test scores attributable to other sources. Some of that variance may still reflect parental influences; our estimates constrain only the component correlated with parental genetic liability for EA, which we find to be modest.

Our results also support the hypothesis that maternal EA may influence risk of premature birth. A prior study found that controlling for maternal BMI, drug use, and other factors attenuated the association between maternal EA and risk of premature birth.^43^ Importantly, after properly accounting for assortment, we did not find any evidence for an association between the paternal EA genetic liability and premature birth (which we had previously observed in naive trio regressions^44^), demonstrating that RAVEL can be used to better clarify both the relative role of direct and indirect genetic effects as well as the relative association of the parental NTCs.

The triangulation of the RAVEL estimate of DGEs with other study designs provided an important plausibility check, as RAVEL requires prespecified structural parameters. Because we estimated the standardized DGE between offspring EA genetic liability and test scores, combining this coefficient with the genetic correlation between EA and test scores gave an implied direct additive heritability for the test phenotype. These estimates were close to additive heritabilities estimated in a large Norwegian pedigree study,^46^ while the visible-scale grade 8 estimate was consistent with direct-effects SNP heritability estimates for cognitive ability and test scores from trio-GCTA and LDSC. The broad concordance was reassuring since the corrected RAVEL direct coefficients are substantially larger than the naive trio PGS coefficients; the triangulation suggests that this increase is consistent with recovery of the full genetic liability rather than an implausible overcorrection. Additionally, reassuringly, the π parameter that recovered the expected genotypic mate correlation (π∼0.46) was consistent with prior estimates comparing the contribution of common and rare variation to EA in UK Biobank (albeit not within-family)^52^. Specifically, that study found that common genetic variants (MAF >5%) explain 58% of the total variance explained by all variants down to a MAF of 0.01% and 49% of the variance relative to a pedigree-based estimate after highly conservative correction for population stratification, suggesting the implied π parameter value in our analysis is reasonable.

As we have emphasised here, polygenic scores are noisy measures of the visible-scale liability indexed by the variants they include, but they also systematically miss classes of trait relevant variation not well tagged by common SNPs (e.g., rare variants, structural variants, variants in low-complexity regions of the genome). Until PGSs are built using every class of genetic variation contributing to trait heritability, this “hidden fraction” will be non-negligible, and even measurement-error-corrected PGS analyses will be biased by AM if one seeks to interpret the NTCs. We note that the considerations presented here regarding unobserved genetic variation apply elsewhere, such as in RDR-SNP and trio-GCTA analyses, since the genetic relatedness matrices are constructed from a subset of typically common genetic variants that do not tag all the variants contributing to trait heritability.^52^ Thus, any variance attributed to parental genotypes should be interpreted with caution when AM is not taken into account. This bears directly on how the existing literature should be interpreted. Prior work using trio-based PGS analyses has either not controlled for AM at all and has occasionally specified it as a limitation,^17–25^ or opted to detect evidence of genetic nurture by considering asymmetric NTCs which is only qualitatively interpretable and would fail to detect genetic nurture effects when they are largely symmetric.^53^

RAVEL can both freely estimate the cross-person residual correlations of the score-specific error terms (i.e., the correlation of the noisy component of the PGSs across family members) as well as have those parameters fixed which, when correctly specified, can increase power especially at small sample sizes. We report the results both for the fixed and freely estimated analyses, which reassuringly showed approximately equivalent parameter estimates (supplementary Table 3). We observed the freely estimated correlations were not significantly different from their expectation, which provides a useful sanity check for model assumptions (i.e., 0.5 between parent and offspring and near 0 between unrelated parents, Supplementary Table 3). When analysing complete trios and additive, common variant PGSs in homogeneous samples, it is reasonable to fix this correlation to 0.5 as done here, though this is not the case when using Mendelian imputed parental PGSs^54^ in mixed cohorts of duos and trios (see Supplementary Note Section 8). Thus, for most common applications, fixing the parameters should increase precision of the other parameter estimates, though care should be taken when doing so to ensure assumptions are met given the score and sample used.

This work has several limitations. First, it requires structural parameters regarding the fraction of heritability missing due to variants not being included or tagged by those in the PGS, and an estimate of the number of generations of constant strength assortment. If there is no external anchoring estimate (e.g., the spousal total-liability genotypic correlation as we use here), these parameters may be varied across a reasonable sensitivity grid. Additionally, we did not allow for arbitrary AM histories to be specified and instead assumed that assortment began and remained constant for the last n-generations before the current generation. However, the *specific* history does not matter as it is purely the within-and between-individual correlations of the visible and hidden components that are needed for RAVEL, and most AM histories likely to be relevant in human populations can be stated as an equivalent ‘effective’ number of generations of constant-strength AM since random mating (Supplementary Note Section 6.3). It is possible to quantify the effective AM duration based on intergenerational PGS (co)variances, which was the feature exploited for the estimates used here.^31^ Second, the PGSs were built using summary statistics from non-Norwegian cohorts and then applied in a Norwegian cohort, MoBa; although the genetic correlation for EA between MoBa and the Okbay et al. GWAS is 0.93, it is significantly below 1.^55^ Thus, the genetic liability being indexed by the EA PGSs is not necessarily reflective of that in the Norwegian population per se. Similarly, the genetic correlation between the EA4 GWAS excluding 23andMe and the 23andMe GWAS is 0.94 and significantly below 1.^37^ If the distinct genetic signals are uncorrelated with the outcome trait, these are effectively uncorrelated noise and thus will not bias estimates obtained by RAVEL; however, if the two distinct genetic signals in the two PGSs are independently associated with the outcome trait, that would theoretically bias the effect size estimates, though likely only minimally when the genetic correlation is relatively high as is the case here (see Supplementary Note Section 9). Third, if the unobserved genetic variation contributing to trait heritability (e.g. *de novo* variation) has not been influenced by the same AM history as the observed genetic variation, then this would violate the assumption of proportional AM influence on the observed and unobserved genetic components. For example, in the case of a trait where *de novo* variation makes a substantial contribution to the trait heritability (e.g. intellectual disability), RAVEL will systematically underestimate DGEs. Fourth, RAVEL also requires two independent GWASs for the genetic liability of interest, which may not be available for all traits of interest. Fifth, because the RAVEL total-liability correction assumes the genetic correlation between the PGS liability trait and the analyzed phenotype is identical across the PGS variants and hidden variant fraction, and some emerging evidence suggests rare variants appear to have higher cross-trait genetic correlations than common variants,^56^ total-liability coefficient estimates may be deflated in cross-trait analyses. Lastly, our analyses were restricted to complete trios that constitute a genetically inferred homogenous subset of the MoBa cohort, which may have led to ascertainment bias.

Here, we have introduced RAVEL, a method to estimate the direct and non-transmitted influence of genetic liabilities on outcomes by leveraging two PGSs constructed from independent GWASs while controlling for AM and measurement error. Where assumptions are met, RAVEL produces unbiased estimates of DGEs and NTCs that are robust to a range of AM and hidden fraction specifications. Future work should incorporate the ability to test mediation of parental associations by parental phenotypes, incorporate polygenic scores from multiple phenotypes into a single model, and allow for additional family relationships to be fit in the SEM (e.g. indirect genetic effects of siblings). Our results suggest that polygenic score trio analyses and SNP-based methods that exploit genetic relatedness for exploring parental genetic effects need to be interpreted with caution, and provide a path forward for estimating interpretable genetic nurture effects.

## Methods

### RAVEL SEM Model

The full mathematical details of the RAVEL model are given in the Supplementary Note. In brief, RAVEL models two independently constructed polygenic scores for each member of a parent-offspring trio as noisy indicators of a latent visible genetic liability for the child, mother and father. The phenotype is then regressed on these three latent liabilities to estimate the offspring’s direct effect and the maternal and paternal indirect effects. The trio latent covariance block includes the spouse covariance and the offspring-parent transmission covariances, and same-score residual covariances are allowed across family members to capture family-correlated GWAS weight noise. When a hidden-heritability fraction π is specified, visible-scale estimates are mapped to a broader total-liability scale using an AM correction based on the implied cross-component covariance structure. Continuous outcomes were fit with robust maximum likelihood; binary outcomes were fit on a probit liability scale (WLSMV). A full derivation of model identification, the visible to total genetic liability mapping correction and the finite-generation versus equilibrium assortment corrections is provided in the Supplementary Note.

### Analytic expectations of DGEs and NTCs

We calculated the expected offspring, maternal and paternal coefficients as functions of the hidden-variant fraction π, genotypic mate correlation ρ, number of generations of constant-strength assortment *g*, and PGS reliability under random mating *r_0_* . We show the formulas below, but the derivations are provided in the Supplementary Note. Let β_T_ = (β*_D_* , β*_M_*, β*_P_*) denote the standardized total-scale liability effects for the offspring (c), mother (m), and father (p).

For a trait with an additive genetic architecture, the total genetic variance after *j* generations of assortment is

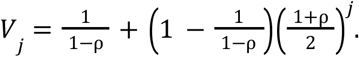

We set *V_m_*_/*p*_ = *V_g_*_−1_ and *V_c_* = *V_g_* . The corresponding visible-scale genetic variances are

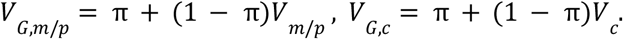

The expected visible-scale offspring coefficient is

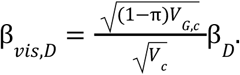

Define

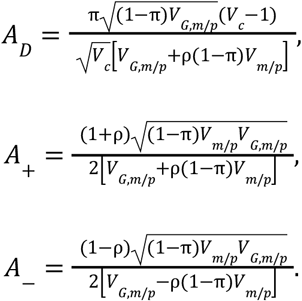

The expected maternal and paternal visible-scale coefficients are then

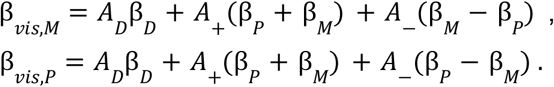

Here, *A_D_* β*_D_* is the portion of the offspring direct effect allocated to parental predictors because they tag hidden offspring liability under assortment, the *A*_+_ term represents the parental average effect, and the *A*_−_ represents the maternal-paternal contrast.

We can then treat the observed PGS expected coefficients as a special case of the visible genetic component. If the visible component captures fraction 1 − π of total genetic liability and the PGS has random mating reliability *r*0 with respect to that component, the PGS captures fraction *r*_0_ (1 − π) of total liability. We therefore defined an effective PGS hidden fraction π*_PGS_* = 1 − *r* _0_(1 − π). Expected PGS-scale trio coefficients were calculated by evaluating the visible-scale expectation functions at π*_PGS_*, while retaining the same AM correlation and number of generations. This reduction follows because score error generated by a shared set of SNP weights has additive parent-offspring covariance consistent with Mendelian transmission.

### One-generation AM forward simulation

The native validation framework was a custom trio simulator designed to match the assumptions of the one-generation AM correction. We simulated 16,000 trios per replicate using 1,000 visible causal variants. When hidden heritability was present, we simulated an equally sized hidden variant set and defined the total genetic liability as a weighted combination of visible and hidden components. Parental pairing was induced by Gaussian-copula sorting on the simulated total genetic liability, after which offspring genotypes were generated by Mendelian transmission. Phenotypes were generated as a linear function of the offspring, maternal and paternal total genetic liabilities plus residual noise. Two observed PGS were then created from the visible variants by adding Gaussian noise to the true SNP effects and applying the same noisy weight vector to all family members, yielding family-correlated measurement error; target reliabilities were 0.50 for PGS1 and 0.30 for PGS2. The main native grid of parameters contained five genetic architectures: no effects (β_D_^2^ = β_M_^2^ = β_P_^2^ = 0), DGE-only (β_D_^2^ = 0.25/0.5, β_M_^2^ = β_P_^2^ = 0), symmetric parental effects (β_D_^2^ = 0.2, β_M_^2^ = β_P_^2^ = 0.05), and direct+maternal (β_D_^2^ = 0.25, β_M_^2^ = 0.10, β_P_^2^ = 0).

We crossed these architectures with spouse genetic correlations of 0, 0.20 and 0.40 and hidden-heritability fractions of 0.40 and 0.80, with 200 replicates per combination of parameters. For every replicate, we fit RAVEL under three analysis conditions: no hidden correction (i.e., visible-scale), the matched hidden fraction with a one-generation AM correction, and the matched hidden fraction with the equilibrium correction. We fit the RAVEL model using lavaan v.0.6-21.^57^

### Multi-generation AM forward simulation

We used the GeneEvolve^32^ multigenerational forward simulator that we modified to expose the last-generation haplotypes, allowing exact reconstruction of trio genotypes. We simulated 15 generations with population size 16,000 and 500 causal variants per trait, with minor allele frequencies sampled uniformly from 0.1 to 0.5, and used the same grid as the one-generation simulation, omitting the simulations of no effects and no assortment. From the reconstructed trio genotypes, we generated two observed PGS using a shared noisy effect vector, as above, calibrated to target reliabilities of 0.50 and 0.30. For every replicate, we fit RAVEL under three analysis conditions: no hidden correction (i.e., visible-scale), the matched hidden fraction with a one-generation AM correction, and the matched hidden fraction with the equilibrium correction. As GeneEvolve does not allow for asymmetric parental effects, we did not include the direct+maternal parameter combination.

### The MoBa Cohort

We applied RAVEL to the Norwegian Mother, Father and Child Cohort Study (MoBa), a population-based pregnancy cohort conducted by the Norwegian Institute of Public Health^58,59^. Pregnant women were recruited between 1999 and 2008, and approximately 41% of those invited consented to participate; the study now includes information on 114,500 children, 95,200 mothers, and 75,200 fathers, with genotyping completed for substantial subsets of each. The present analysis was restricted to complete genotyped trios for whom at least one of the focal outcome measures described below was available. Quality control, genotype imputation, and ancestry and relationship determinations are described in ^60^. Sample sizes for each regression are provided in Supplementary Table 4.

### Outcome measures

We considered three outcomes, all derived from Norwegian national registries linked to the MoBa cohort: standardized national test (Nasjonale prøver) score averages at grade 5 and grade 8 (grade 9 was excluded as only two of the three tests are taken), and gestational age. National test score averages were computed within each grade level by averaging across the three subjects administered (reading comprehension in Norwegian, mathematics, and English as a foreign language). Score averages were treated as continuous measures and residualised on child birth year as a categorical variable, genotyping batch, sex, age at testing, age at testing squared, and the first twenty principal components computed from the genotyped sample. We additionally constructed binary indicator variables of the top 10th and the top 90th percentiles of the residualised distribution within each grade level, which were used in secondary analyses targeting the upper and lower tails of academic performance, respectively. Gestational age was measured in weeks, and preterm birth was defined as gestational age at birth less than or equal to 37 weeks and 0 days. The binary outcomes were modeled with the same fixed effect covariate set as above, fit on the probit scale which assumes an underlying normal liability distribution.

### Polygenic scores

We constructed two polygenic scores per individual using summary statistics from two independent EA GWASs with non-overlapping discovery samples used in Okbay et al.^37^ (commonly referred to as “EA4”). The first score was built from the EA GWAS with the 23andMe contribution excluded from the meta-analysis; the second score was built from the 23andMe-only educational-attainment GWAS. We limited variants to those that were well imputed in MoBa, present in both GWAS summary statistics, and that were in the HapMap3 SNP list, resulting in 1,305,822 autosomal SNPs.^61^ Scores were generated with in-sample LD matrices using LDpred-inf as implemented in *bigsnpr* v1.12.20.^62,63^ We applied each score to all genotyped trio members (mother, father, child) using the same set of weights per score. The two raw scores were each residualised on the same covariate set used for the outcomes (birth year, genotyping batch, sex, age, age^2^, and the first twenty genetic principal components), and were then centred and scaled for all family members using the child’s mean and standard deviation, preserving the parent-child mean and variance contrast on a common reference scale.

### RAVEL estimates in MoBa

For each outcome we fit RAVEL to the trio data using *lavaan* v.0.6-21.^57^ Two structural parameters are not identified within RAVEL and must be supplied: the hidden fraction π (see Box 1) and the number of generations of constant strength assortment. We fixed the assortment history at three generations, the value estimated for EA in Norway from intergenerational polygenic score (co)variances.^4,31^ We then set π by estimating for each candidate value a refit model and considered the implied spousal correlation of the parental total genetic liabilities, selecting the value at which this matched the independently estimated Norwegian EA genotypic mate correlation of 0.34.^4,31^ This yielded π = 0.46. Because π was chosen to reproduce ρ, the recovered ρ is not an independent validation of the model. We further considered every combination of π varying across (0.4, 0.42, 0.44, 0.46, 0.48, 0.5) and 1, 2, 3, 4, 5, and 15 generations of AM. Standard errors and 95% confidence intervals for the visible-scale and total-scale coefficients were derived using the *lavaan* asymptotic variance-covariance matrix.

### Full-sibling IBD regression

We identified full sibling probands in MoBa and estimated total IBD sharing between siblings using KING 2.3.1 using the *--ibdseg* command. We then fit the full-sibling IBD regression using previously published code.^64^

### Genome wide association studies and genetic correlations

GWASs were conducted using REGENIE v3.1, with genotype batch, sex, and 10 genetic PCs included as covariates.^65^ SNP heritabilities and genetic correlations were then estimated using LD score regression with the provided European reference LD panel on Hapmap3 SNPs^61^.^66,67^

### Pedigree-based additive heritability estimation from relative phenotype correlations

We estimated additive heritability and genotypic mate correlations from published correlations between relative within different kinship classes using Norwegian registry data using the “bio + relatives-in-law” model from the original analysis code.^46^ Adoptive relatives and unknown-zygosity categories were excluded. Models were fit by weighted nonlinear least squares on log relative correlations, weighting each row by the inverse squared standard error.

To test whether test score heritability increased from grade 5 to grade 8, we used a paired bootstrap over matched relative categories. Grade 5 and grade 8 correlations were matched by relative type, sex composition, maternal/paternal status, half-relative status, relatedness distance, and relatives-in-law status. In each of 5,000 bootstrap replicates, matched relative-category pairs were sampled with replacement, the heritability model was refit separately at each age, and the difference in the heritability estimates was estimated. Bootstrap standard errors, percentile confidence intervals, and two-sided p-values were calculated from the empirical distribution of these paired differences.

## Acknowledgments

We are grateful to Peter Visscher for suggesting the title and for critical discussion of the work, in particular the Supplementary Note and the framing of the results. We are grateful to Qin Qin Huang for comments on the manuscript.

We would like to thank the research participants and employees of 23andMe Research Institute for making this work possible.

The Norwegian Mother, Father, and Child Cohort Study is supported by the Norwegian Ministry of Health and Care Services and the Ministry of Education and Research. We are grateful to all the participating families in Norway who take part in this on-going cohort study. We also thank the NORMENT Center for providing genotype data, funded by the Research Council of Norway (#223273), South East Norway Health Authorities, and Stiftelsen Kristian Gerhard Jebsen, and in collaboration with deCODE Genetics. We further thank the Center for Diabetes Research, the University of Bergen for providing genotype data funded by the ERC AdG project SELECTionPREDISPOSED, Stiftelsen Kristian Gerhard Jebsen, Trond Mohn Foundation, the Research Council of Norway, the Novo Nordisk Foundation, the University of Bergen, and the Western Norway Health Authorities. The MoBa work was performed on the TSD (Tjeneste for Sensitive Data) facilities, owned by the University of Oslo, operated and developed by the TSD service group at the University of Oslo, IT Department (USIT).

This research was funded in part by Wellcome (grant no. 220540/Z/20/A, “Wellcome Sanger Institute Quinquennial Review 2021–2026”). This work was partly supported by the Research Council of Norway through its Centres of Excellence funding scheme (#262700). This work was supported by the Research Council of Norway (#334093, #336085) and South-Eastern Norway Regional Health Authority (#2020022, #2026069) and the Horizon 2020 Research and Innovation programme of the European Union (FAMILY, grant agreement no. 101057529; Marie Skłodowska-Curie grant ESSGN no. 101073237).) . This work was performed on the TSD (Tjenester for Sensitive Data) facilities, owned by the University of Oslo, operated and developed by the TSD service group at the University of Oslo, IT-Department (USIT). For the purpose of open access, the authors have applied a CC-BY public copyright license to any author accepted manuscript version arising from this submission.

## Contributions

DSM conceived the method, wrote the code developed in this work, conducted all of the analyses, and wrote the first draft of the manuscript. HFS contributed intellectually to the development of the technical Supplementary Note and provided input on the manuscript. LH and OW assisted in the data preparation in MoBa and provided input on the manuscript. AH and HCM supervised the analyses and provided input on the manuscript. All authors read and commented on the final manuscript.

## Code availability

RAVEL, simulation code, and analytic expected coefficient value calculator are available at https://github.com/malawsky/RAVEL.

## Data availability

The Okbay et al. GWAS summary statistics can be downloaded from http://www.thessgac.org/data. The full GWAS summary statistics from 23andMe are available through 23andMe to qualified researchers under an agreement with 23andMe that protects the privacy of the 23andMe participants. Please visit https://research.23andme.com/collaborate/#dataset-access/ for more information and to apply to access the data. Data from the Norwegian Mother, Father and Child Cohort Study (MoBa) used in this study are managed by the national health register holders in Norway (Norwegian Institute of Public Health) and can be made available to researchers, provided approval from the Regional Committees for Medical and Health Research Ethics (REC), compliance with the EU General Data Protection Regulation (GDPR) and approval from the data owners. The consent given by the participants does not open for storage of data on an individual level in repositories or journals. Researchers who want access to data sets for replication should apply through helsedata.no. Access to data sets requires approval from The Regional Committee for Medical and Health Research Ethics in Norway and an agreement with MoBa.

---

**Supplementary Figure 1.**
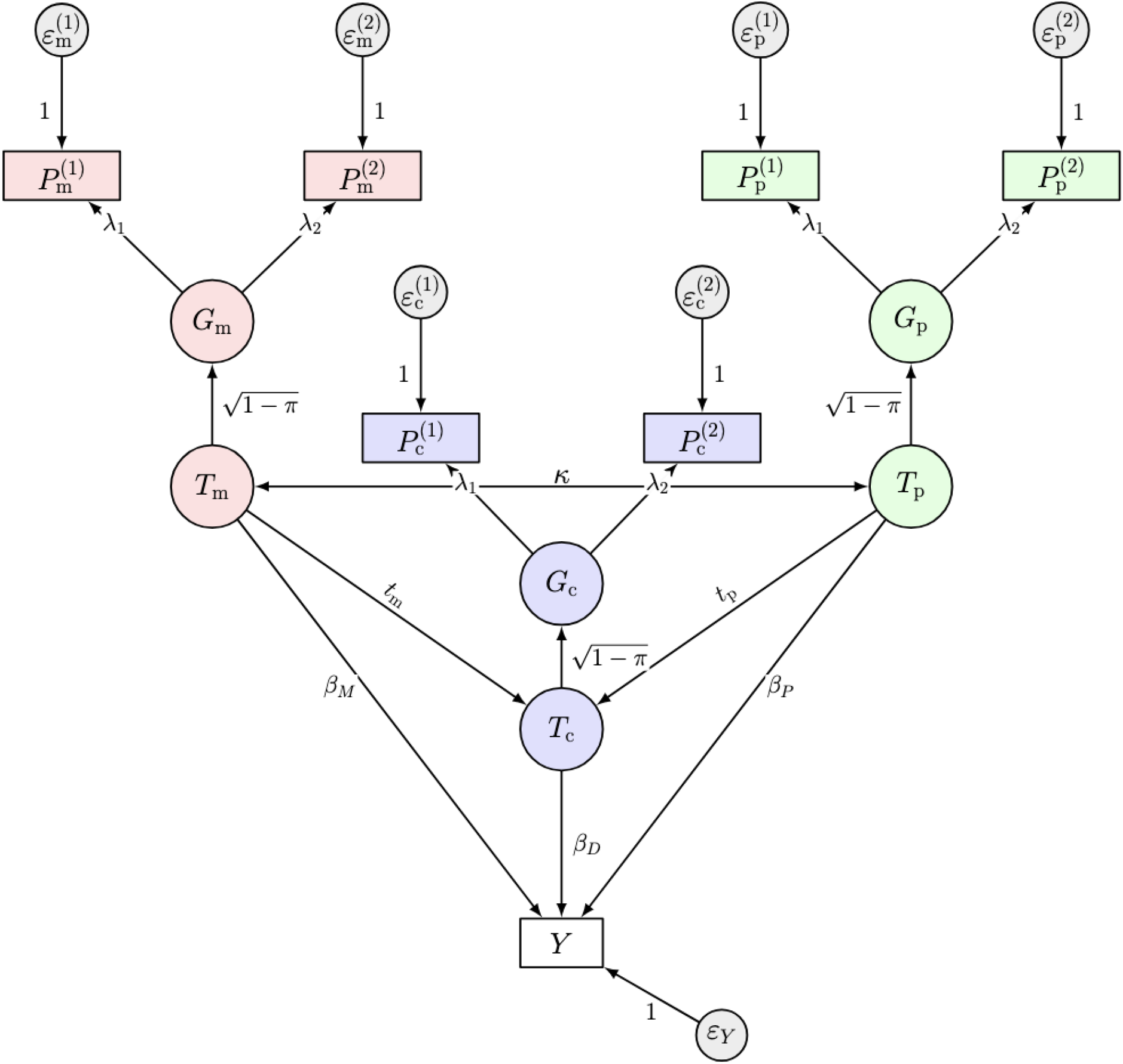
Conceptual path diagram for the RAVEL trio model with explicit total-liability nodes. Each family member has a total additive genetic liability *T*_r_, r ∈ {m (mother), p (father), c (child)}, and a visible genetic factor *G*_r_ indexed by two independently constructed polygenic scores, P_r_^(1)^ and P ^(2)^. The loading from *T*_r_ to *G*_r_ is (1−π)^1/2^, corresponding to the share of additive genetic variance captured by the observed scores when there is hypothetically no assortative-mating-induced tagging. AM is represented on the total-liability scale by the covariance between *T*_m_ and *T*_p_, denoted κ (and ρ when expressed as a correlation). The offspring total liability *T*_c_ is generated from the parental total liabilities with transmission coefficients *t*_m_ and *t*_p_, and the offspring phenotype Y is regressed on *T*_c_, *T*_m_, and *T*_p_ with coefficients *β_D_*, *β_M_*, and *β_P_*.

**Supplementary Figure 2.**
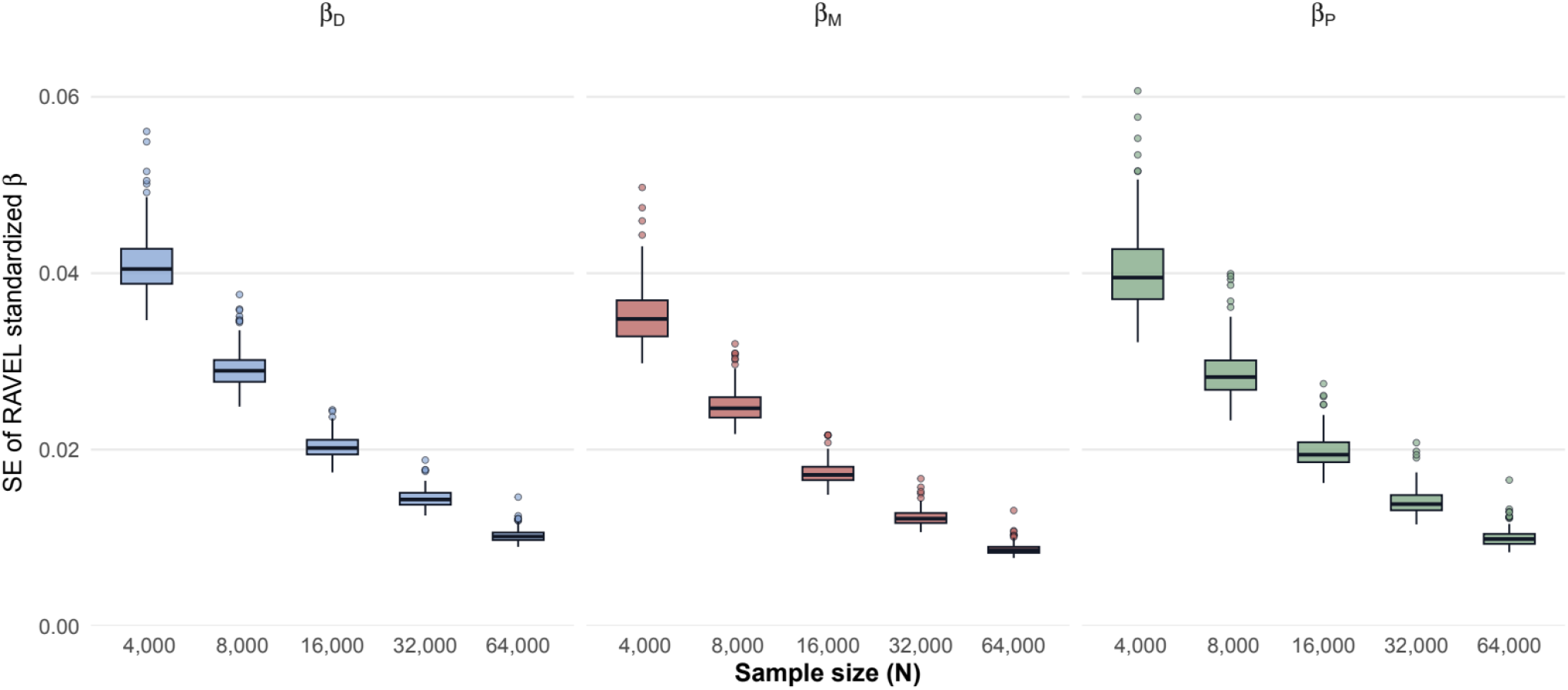
Distribution of standard errors across different numbers of full trios observed. Standard errors estimated by fitting RAVEL with the correct AM and π values assuming one generation of AM for a trait with an asymmetric direct+maternal architecture (β_D_ = 0.5, β_M_ = 0.32, β_P_ = 0) at ρ = 0.4 and π = 0.4 across 200 simulation replicates with 16,000 trios. The two PGSs in the simulation were calibrated to have reliabilities of 0.3 and 0.5 as done in the simulations described in the Results.

**Supplementary Figure 3.**
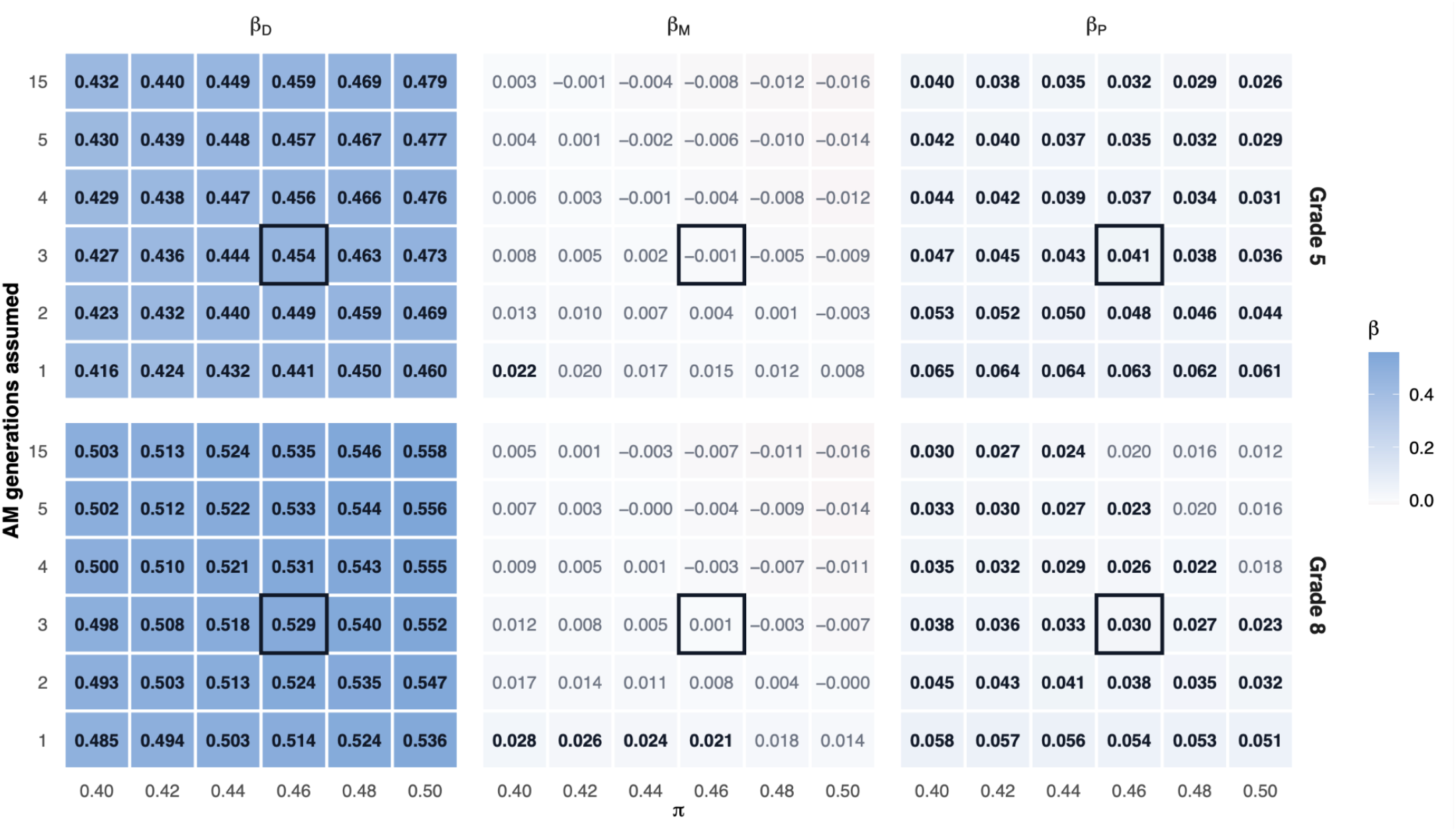
Sensitivity to varying assumed numbers of generations of AM and π. Estimated standardized RAVEL coefficients for the DGEs and NTCs for grades 5 and 8 test scores assuming different combinations of generations of AM and π. Black squares indicate the combination used in the main text analyses. Bold values indicate where the estimate is at least nominally significantly different from 0 (p<.05 two-sided Wald test). Full results from the sensitivity analyses are available in Supplementary Table 5.

**Supplementary Figure 4.**
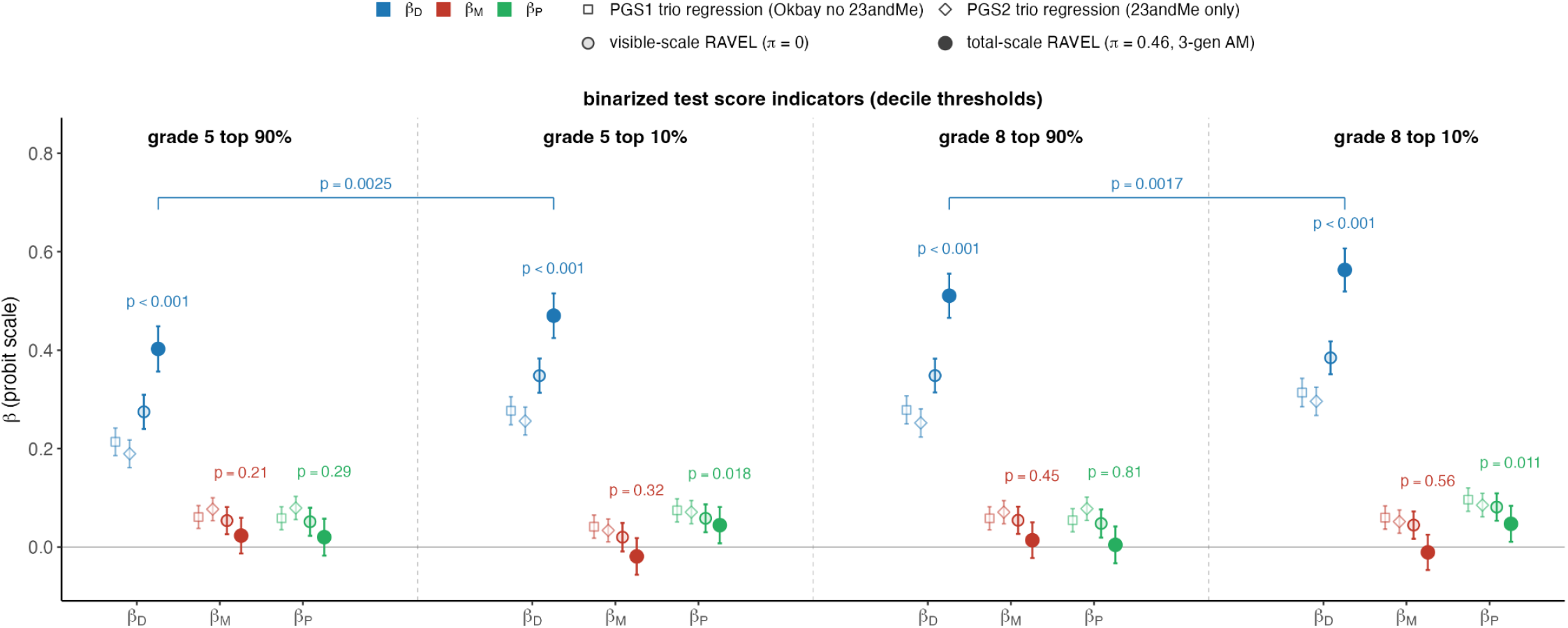
RAVEL applied to the top 10th and 90th percentiles of test scores in MoBa. Each panel reports point estimates with 95% confidence intervals from four estimators applied to complete genotyped MoBa trios (n trios=34,295). Open squares are PGS1 trio OLS estimates (PGS1 constructed from the Okbay et al. 2022 educational-attainment GWAS with 23andMe excluded) and open diamonds are PGS2 trio OLS estimates (PGS2 constructed from the 23andMe-only EA GWAS). Open circles are visible-scale RAVEL estimates fit with no correction (π = 0), and filled circles are total-liability RAVEL estimates corrected with π = 0.46 and three generations of assortment. Coefficients are color-coded: blue for the offspring direct genetic effect β_D_, red for the maternal non-transmitted coefficient β_M_, and green for the paternal non-transmitted coefficient β_P_. Outcomes are reported on the probit scale for binary indicators for the top 90th and 10th percentiles of the residualised grade 5 and grade 8 distributions.

**Supplementary Table 1.** RAVEL performance in the one-generation trio simulator. Summary of RAVEL estimates across 200 replicates of 16,000 trios per parameter cell in the custom one-generation forward simulator (Methods; Supplementary Note Section 10). The simulation grid crosses five genetic architectures: direct-only with β_D_ ∈ {0, 0.5, 0.71} (direct_only_h2_0.00, direct_only_h2_0.25, direct_only_h2_0.50), direct+maternal (β_D_ = 0.5, β_M_ = 0.32, β_P_ = 0), and symmetric both-parents (β_D_ = 0.45, β_M_ = β_P_ = 0.22), with three AM strengths (ρ ∈ {0, 0.2, 0.4}) and two hidden-variant fractions (π ∈ {0.4, 0.8}). Architectures are specified as variance components in the table (h2_d_input, h2_m_input, h2_p_input); the corresponding standardized coefficients are their square roots. Each cell is fit under three analysis conditions, given in the “correction” column: no hidden-variant correction (h0), which targets the visible scale; the true hidden fraction with a one-generation assortment correction (h_true_1gen); and the true hidden fraction with the equilibrium correction (h_true_equil). Because the one-generation simulator generates data under one generation of assortment, the h_true_1gen rows constitute the correctly specified condition and the h_true_equil rows quantify the bias incurred by misspecifying the assortment history. Four estimands are reported per cell (parameter): the offspring direct genetic effect (beta_D), the maternal and paternal non-transmitted coefficients (beta_M, beta_P), and the spousal correlation of the latent genetic liabilities (rho). We estimate the realized true parameters in each simulation run as well as estimate the parameters with RAVEL, so both estimates and truths are summarized across replicates.

**Supplementary Table 2.** RAVEL performance in the fifteen-generation trio simulator. Summary of RAVEL estimates across 200 replicates of 16,000 trios per parameter cell in the GeneEvolve forward-time simulator, run for 15 generations of constant-strength assortment (approximately equilibrium). The grid covers two direct-effect magnitudes (latent_h2_input ∈ {0.25, 0.5}, i.e. β_D_ = 0.5 and 0.71), two hidden-variant fractions (prop_h2_latent ∈ {0.4, 0.8}), two AM settings (phenotypic mate correlation, am11 ∈ {0.2, 0.4}; this parameter is the assortment strength supplied to GeneEvolve), and the presence or absence of symmetric parental indirect effects (phenotypic parental effects, f11 ∈ {0, 0.1}). The realized spousal correlation of the total genetic liability is reported directly in the rho rows. The correction, truth_scale, and parameter columns are defined as in Supplementary Table 1. Because the data-generating process here is 15 generations of assortment, the h_true_equil rows are the correctly specified condition and the h_true_1gen rows quantify the bias from assuming too short an assortment history.

**Supplementary Table 3.** RAVEL and PGS trio regression estimates for all MoBa outcomes. Full association statistics from complete genotyped MoBa trios for seven outcomes: average national test scores at grade 5 and grade 8; binary indicators for the top 90th and 10th percentiles of the residualised score distribution at each grade level; and premature birth (gestational age ≤ 37 completed weeks). Continuous outcomes are reported on the standardized phenotype scale and binary outcomes on the probit liability scale. All outcomes and polygenic scores were residualised on the covariate set given in Methods. Each row reports the point estimate, standard error, and p value for one parameter under one model, with the seven outcomes given side by side. Three model classes are reported in the Model column. The two observed-score regressions are standard OLS trio regressions of the outcome on the child, maternal, and paternal PGS, fit separately for PGS1 (Okbay et al. 2022 EA GWAS excluding 23andMe) and PGS2 (23andMe-only EA GWAS). The two RAVEL models jointly model both scores as indicators of a latent genetic liability per family member. The Cross-person PGS error structure column distinguishes the two RAVEL specifications. In the fixed specification, the cross-person PGS residual correlations are constrained to their Mendelian expectations: 0 between mother and father, who are unrelated, and 0.5 for each parent-offspring pair, reflecting the expected sharing of the score-specific error component through transmission. In the free specification, the same six correlations (three per score) are freely estimated. Estimates of the direct and non-transmitted coefficients are near-identical across the two specifications; the fixed model is used in the main text.

**Supplementary Table 4.** Model fit statistics for the RAVEL structural equation models fit in MoBa. Global fit statistics for the two RAVEL specifications reported in Supplementary Table 3, fit to each of the seven MoBa outcomes. The upper block reports the fixed specification, in which the six cross-person score-residual correlations are constrained to their Mendelian expectations (0 between mother and father; 0.5 for each parent-offspring pair within each score). The lower block reports the free specification, in which those six correlations are freely estimated, yielding 3 degrees of freedom. Continuous outcomes (grade 5 and grade 8 average test scores) were fit with robust maximum likelihood; binary outcomes (top 10th and 90 percentile of the residualised grade distributions, and premature birth) were fit on a probit liability scale, for which AIC and BIC are not defined and are therefore left blank. All models converged.

**Supplementary Table 5.** Sensitivity of RAVEL estimates to the assumed hidden-variant fraction and assortative mating history. Grid search over the two structural parameters that RAVEL cannot identify internally: the fraction of total heritability not captured by the polygenic scores under random mating (π) and the number of generations of constant-strength AM. Estimates are reported for every combination of π ∈ {0.40, 0.42, 0.44, 0.46, 0.48, 0.50} and n ∈ {1, 2, 3, 4, 5, 15} generations, applied to each of the seven MoBa outcomes. Four total-liability-scale quantities are given per cell: the offspring direct genetic effect (bD), the maternal and paternal non-transmitted coefficients (bM, bP), and the spousal correlation of the parental total genetic liabilities (rhoMF). Continuous outcomes are on the standardized phenotype scale and binary outcomes on the probit scale.

**Supplementary Table 6.** Full-sibling IBD regression applied to national test scores. Parameter estimates and standard errors for the a^2^, c^2^, and e^2^ components and number of sibling pairs.

| School grades | $c^2$ | $a^2$ | $e^2$ | N sibling pairs |
| --- | --- | --- | --- | --- |
| Grade 5 | 0.1177 (0.09323) | 0.5356 (0.1855) | 0.3258 (0.09287) | 9,823 |
| Grade 8 | -0.01419<br>(0.1074) | 0.8681 (0.2128) | 0.1447 (0.1059) | 7,035 |

**Supplementary Table 7.** PGS variances and covariances. Variances (diagonal; normalized to the child) and correlations (off-diagonal) of the EA4 (PGS1) and 23andMe (PGS2) PGSs across family members in 34,295 complete trios.

|  | PGS1 child | PGS1 mother | PGS1 father | PGS2 child | PGS2 mother | PGS2 father |
| --- | --- | --- | --- | --- | --- | --- |
| PGS1 child | 1 |  |  |  |  |  |
| PGS1 mother | 0.5642 | 0.9789 |  |  |  |  |
| PGS1 father | 0.5710 | 0.1490 | 0.9917 |  |  |  |
| PGS2 child | 0.6603 | 0.3894 | 0.4018 | 1 |  |  |
| PGS2 mother | 0.3908 | 0.6534 | 0.1413 | 0.5667 | 0.9778 |  |
| PGS2 father | 0.3971 | 0.1382 | 0.6614 | 0.5785 | 0.1442 | 1.0148 |

## Notes

### Competing Interest Statement

The authors have declared no competing interest.

### Summary of Updates

Added a clarifying sentence to the introduction and an edit to a sentence in the Discussion; Acknowledgments updated.

