## Supplementary Note for "Phantom genetic nurture: assortative mating accounts for most of the apparent association between parental genotypes and childhood cognitive performance"

### Supplementary Note: RAVEL theory

Recovering Assortment Adjusted Variance Explained by Genetic Liabilities

### 1 Overview

RAVEL is a trio structural equation framework for estimating direct offspring and parental genetic-liability-correlated variance explained by leveraging two independently constructed polygenic scores per individual as multiple noisy indicators of a common latent genetic liability. The observed quantities are noisy polygenic scores in trios and an offspring phenotype. The quantities identified by the structural equation model are score-indexed latent genetic components, that is, the fraction of additive genetic liability indexed by the measured variants used to construct the observed scores. The phenotype, however, may depend on a broader total liability that includes variation not represented in the observed scores. Inference on the visible scale and inference on the total-liability scale are therefore distinct inferential problems we address here.

For exposition in the manuscript's main text, the special case with no explicit hidden component, an outbred population, and additive transmission coefficients fixed at one half on the raw genetic scale is a natural entry point. The present supplementary note generalizes the visible-scale framework to accommodate Mendelian imputation, consanguinity, and scores that are not exactly linear sums of directly observed variants (e.g., scores for traits that may include interaction or dominance terms). The finite generation hidden component correction and the scalar expectations used in the main text, however, are derived under an additive infinitesimal model with constant genic segregation variance. Non-additive score systems require their own cross-generation covariance model before the same total-liability interpretation can be made.

The trio setting makes this distinction especially important. Measurement error in the observed scores does not simply attenuate the offspring coefficient. Because the parental scores are correlated with the offspring's latent liability through Mendelian transmission and potentially assortative mating, the portion of the offspring's liability not captured by the observed offspring score can be partially represented by the parental regressors. As a result, parental coefficients in ordinary trio regression may reflect a mixture of genuine parental pathways and assortative mating-induced tagging. Mapping the visible scale to the total-liability scale depends on the trio covariance structure.

The purpose of the present supplementary note is to state the statistical model and describe the simulation framework we developed. We begin with the visible-scale trio structural equation model, including the measurement model for the observed scores, the trio covariance structure, and the phenotype regression. We then derive the projection identity linking the fitted visible-scale coefficients to the total-liability coefficients. Next, we derive the raw visible and hidden component variances and their cross-component covariance matrix under three regimes: a one-generation assortative mating setting, an exact $n$-generation recursion, and the equilibrium assortative mating regime. The following section then shows how these visible-scale quantities change when one parent is Mendelian-imputed rather than observed.

### 2 Observed scores, visible-scale genetic liability, hidden components, and total liability

For each family we observe a mother, a father, and an offspring. Subscripts indicate m for the maternal role, p for the paternal role, and c for the child. The observed quantities are two score versions for each role,

$$P_r^{(1)},\ P_r^{(2)}, \qquad r \in \{\mathrm{c}, \mathrm{m}, \mathrm{p}\}. \tag{2.1}$$

Because the derivations concern a generic family, we suppress the family index $i$ from here onward except where simulation statements require it explicitly.

The statistical model distinguishes a raw visible additive component

$$G_r^*, \qquad r \in \{\mathrm{c}, \mathrm{m}, \mathrm{p}\}, \tag{2.2}$$

which is the genetic liability recoverable from the measured variants used to construct the observed scores, from a raw hidden additive component

$$H_r^*. \tag{2.3}$$

Let

$$\omega_G = \sqrt{1-\pi}, \qquad \omega_H = \sqrt{\pi}, \qquad \pi \in [0, 1), \tag{2.4}$$

and define the raw total genetic liability

$$T_r^* = \omega_G G_r^* + \omega_H H_r^*. \tag{2.5}$$

The reference, or founder, generation is scaled so that

$$\mathrm{Var}(G_0^*) = \mathrm{Var}(H_0^*) = 1, \qquad \mathrm{Cov}(G_0^*, H_0^*) = 0. \tag{2.6}$$

Consequently, $\mathrm{Var}(T_0^*) = 1$ and $\pi$ is the hidden variance fraction on this founder or random-mating reference scale. Under assortative mating, the raw variances of $G^*$ and $H^*$ and their within-person covariance are allowed to change across generations. Thus $\pi$ remains the fixed reference-scale weighting parameter; it is not generally equal to the marginal fraction of present-day raw total-liability variance attributable to $H^*$ after assortative mating-induced variance inflation. The observed scores will often index common, tagged variation, whereas the hidden component may contain rare, structural, or de novo variation. The correction below assumes that the modeled portions of both components are additive and that assortative mating induces covariance between the visible and hidden components in proportion to their respective covariances with the total liability. This does not require $T$ to be the only trait relevant to mate choice. Additional traits influencing assortment are compatible with the model provided that their genetic covariance with the visible and hidden components is proportional to that with $T$. If the hidden component has a materially different mating or transmission process, a new extension will be required.

For the fitted structural equation model and for reporting standardized coefficients, define role-specific standardized liabilities

$$G_r = \frac{G_r^*}{\sqrt{V_{G,r}}}, \qquad H_r = \frac{H_r^*}{\sqrt{V_{H,r}}}, \qquad T_r = \frac{T_r^*}{\sqrt{V_{T,r}}}, \tag{2.7}$$

where

$$V_{G,r} = \mathrm{Var}(G_r^*), \qquad V_{H,r} = \mathrm{Var}(H_r^*), \qquad V_{T,r} = \mathrm{Var}(T_r^*). \tag{2.8}$$

Equivalently,

$$T_r = \frac{\omega_G \sqrt{V_{G,r}}\, G_r + \omega_H \sqrt{V_{H,r}}\, H_r}{\sqrt{V_{T,r}}}. \tag{2.9}$$

The raw causal components in Equations (2.2)–(2.5) are propagated between generations without rescaling. Equation (2.7) is applied only when a given parental or offspring generation is represented on the fitted SEM scale.

We define the standardized trio vectors

$$G = \big(G_{\mathrm{c}}, G_{\mathrm{m}}, G_{\mathrm{p}}\big)^{\top}, \qquad H = \big(H_{\mathrm{c}}, H_{\mathrm{m}}, H_{\mathrm{p}}\big)^{\top}, \qquad T = \big(T_{\mathrm{c}}, T_{\mathrm{m}}, T_{\mathrm{p}}\big)^{\top}, \tag{2.10}$$

and their raw counterparts $G^*$, $H^*$, and $T^*$ analogously. We write

$$\Sigma_{GG} = \mathrm{Cov}(G), \qquad \Sigma_{GH} = \mathrm{Cov}(G, H), \qquad \Sigma_{HH} = \mathrm{Cov}(H). \tag{2.11}$$

The visible-scale trio model identifies $\Sigma_{GG}$. The hidden variant correction requires assumptions sufficient to determine $\Sigma_{GH}$ and, when desired, $\Sigma_{HH}$ from the raw component process. Unless stated otherwise, the observed polygenic scores are treated as indexing the same liability whose phenotype is being modeled. The visible–hidden decomposition in Equation (2.5) therefore refers to a single trait throughout.

#### 2.1 External information for the hidden variance fraction

The hidden variance fraction $\pi$ is not identified by the visible-scale score system alone because the observed scores measure only the visible component by construction. In practice, $\pi$ must therefore be supplied from outside the fitted SEM or explored over a plausible range. A natural calibration is obtained by comparing a visible-scale estimate of additive genetic variation represented by the score-derived latent factor to an external estimate of the total additive genetic variation for the same focal trait and outcome scale. If $V_{A,\mathrm{vis}}$ denotes the additive variance represented by the visible score-indexed component and $V_{A,\mathrm{tot}}$ denotes an externally identified estimate of total additive variance, then a natural calibration is

$$\pi = 1 - \frac{V_{A,\mathrm{vis}}}{V_{A,\mathrm{tot}}}. \tag{2.12}$$

Because $\pi$ is defined by Equation (2.6), the numerator and denominator in Equation (2.12) must refer to the same founder, genic, or random-mating reference scale. A ratio of contemporary assortative mating-inflated variances is not automatically the same parameter unless the corresponding variance inflation has first been modeled or removed. The external quantity $V_{A,\mathrm{tot}}$ may be inferred from family-based estimators such as pedigree-based estimates, relatedness disequilibrium regression, or sib regression, provided that the estimate is on the same trait and reference scale as the liability being modeled. Alternatively, the mate genotypic correlation can be estimated in extended pedigrees and used to anchor the total-scale genotypic mate correlation. The correction developed below should therefore be interpreted as conditional on a chosen value of $\pi$ and the assumed number of generations of constant assortment rather than as identifying $\pi$ internally.

### 3 Glossary of recurring terminology

Because the method distinguishes observed scores, visible-scale genetic liability, hidden genetic components, and total liabilities, the same words recur throughout the note in closely related but not identical meanings.

| Term | Symbol | Meaning |
|---|---|---|
| Observed polygenic score | $P^{(1)}, P^{(2)}$ | The measured scores available in the data. |
| Raw visible genetic component | $G^*$ | The additive component indexed by the measured variants, propagated across generations on a fixed founder scale. Its variance may change under assortative mating. |
| Visible-scale genetic liability | $G$ | The role-standardized copy of $G^*$ identified by the observed scores in the fitted SEM. |
| Raw hidden genetic component | $H^*$ | The additive component not represented in the observed scores, propagated on the same fixed founder scale as $G^*$. Its variance may also change under assortative mating. |
| Hidden genetic liability | $H$ | The role-standardized copy of $H^*$. It is not identified from visible-scale data alone. |
| Raw and standardized total liability | $T^*, T$ | The raw liability is $T^* = \sqrt{1-\pi}\, G^* + \sqrt{\pi}\, H^*$; $T$ is its role-standardized version used for standardized coefficient reporting. |
| hidden variance fraction | $\pi$ | The proportion of total additive variance attributable to the hidden component in the founder or random mating reference population. |
| Score loading | $\lambda_{kr}$ | The loading of score version $k$ for role $r$ on the visible-scale liability. |
| Score-residual variance | $\theta_{kr}$ | The residual variance of score version $k$ in role $r$ after conditioning on the common visible-scale liability. |
| Visible-scale spouse correlation | $\rho_G$ | The genotypic spouse correlation on the visible scale. It is not a phenotypic mate correlation. |
| Total-liability spouse correlation | $\rho_T$ | The standardized spouse correlation on the total-liability scale implied by assortment on the broader liability rather than on the visible component alone. |
| Total-liability spouse covariance | $\kappa_T$ | The raw spouse covariance on the total-liability scale. |
| Offspring–parent visible covariance | $\eta_{\mathrm{m}}, \eta_{\mathrm{p}}$ | The maternal and paternal visible-scale offspring–parent covariance parameters. |

| Term | Symbol | Meaning |
| --- | --- | --- |
| Visible transmission coefficients | $t_{\mathrm{m}}, t_{\mathrm{p}}$ | The population projection coefficients obtained by regressing the role-standardized offspring visible liability on the standardized parental visible liabilities. Raw additive inheritance has coefficients one half; after role-specific standardization the fitted coefficients need not equal one half in a transient AM generation. |
| Within-person visible–hidden covariance | $C_{GH,r}, d_r$ | $C_{GH,r} = \mathrm{Cov}(G_r^*, H_r^*)$ is the raw covariance; $d_r = C_{GH,r}/\sqrt{V_{G,r}V_{H,r}}$ is its standardized counterpart. It is zero in the founder generation and accumulates under AM. |
| Cross-spouse visible–hidden covariance | $K_{GH}, q$ | $K_{GH} = \mathrm{Cov}(G_{\mathrm{m}}^*, H_{\mathrm{p}}^*)$ is the raw covariance induced by assortment on $T^*$; $q = K_{GH}/\sqrt{V_{G,\mathrm{par}}V_{H,\mathrm{par}}}$ is its standardized counterpart. |
| Score residual | $\varepsilon$ | The part of an observed score not explained by the common visible-scale liability. If relatives' scores are constructed using the same weights, these residuals are correlated across relatives in proportion to their genetic sharing. |
| Within-score cross-person residual covariance | $\psi_{k,rs}$ | The covariance between score residuals for roles $r$ and $s$ within score version $k$. The spouse term is kept estimable in the general SEM because consanguinity, endogamy, or other departures from the outbred benchmark can make it nonzero. |
| Visible-scale coefficient vector | $b$ | The regression coefficients of the phenotype on the visible-scale liability returned by the fitted SEM. |
| Total-liability coefficient vector | $\beta$ | The coefficient vector for the role-standardized total liabilities $T$. Coefficients on raw total liabilities are denoted $\beta^{\mathrm{raw}}$. |
| Visible trio covariance matrix | $\Sigma_{GG}$ | The covariance matrix of the fitted visible-scale liabilities. In the trio model this contains $\rho_G$ together with the offspring–parent covariances $\eta_{\mathrm{m}}, \eta_{\mathrm{p}}$. |
| Visible–hidden cross-block | $\Sigma_{GH}$ | The covariance matrix linking the role-standardized visible and hidden components. It is obtained by propagating raw component variances and covariances and then applying role-specific standardization. |

### 4 Visible-scale trio structural equation model

#### 4.1 Measurement model for the observed scores

For each role $r \in \{\mathrm{c}, \mathrm{m}, \mathrm{p}\}$, the two observed scores derived from independent GWASs of the same trait are modeled as noisy measurements of the same visible-scale liability:

$$P_r^{(1)} = \lambda_{1r} G_r + \varepsilon_r^{(1)}, \tag{4.1}$$

$$P_r^{(2)} = \lambda_{2r} G_r + \varepsilon_r^{(2)}. \tag{4.2}$$

The liabilities are standardized to unit variance on the fitted visible scale,

$$\mathrm{Var}(G_r) = 1, \tag{4.3}$$

so the factor loadings are interpretable on a standardized visible-genetic scale. The fitted SEM allows the loadings to vary by both score version and role. This is important because the loading is not the score-construction weight itself, but the covariance between the observed score and the visible liability on the fitted scale. Assortment, role-specific residual structure, and Mendelian imputation can therefore produce role-specific loadings even when the same underlying score-construction weights are used across family members. We likewise retain role-specific score-residual variances,

$$\theta_{kr} = \mathrm{Var}(\varepsilon_r^{(k)}), \qquad k \in \{1, 2\}, \tag{4.4}$$

so that role-specific differences in measurement quality are represented directly.

The corresponding model implied reliability is

$$\mathrm{Rel}\left(P_r^{(k)} \mid G_r\right) = \frac{\lambda_{kr}^2}{\lambda_{kr}^2 + \theta_{kr}}. \tag{4.5}$$

If the observed polygenic score has been standardized to variance one, then Equation (4.5) implies

$$\theta_{kr} = 1 - \mathrm{Rel}\left(P_r^{(k)} \mid G_r\right). \tag{4.6}$$

This simple identity is useful when translating estimated cross-person residual covariances into implied residual correlations.

When disequilibrium is discussed below, the raw visible variance $V_{G,r}$ need not remain one across generations.

#### 4.2 Cross-person score-residual covariance

Within a given score version, the same noisy GWAS-derived weight vector is applied to every person in the family. Consequently, even after conditioning on the common visible-scale liability, the score residual remains correlated across relatives. We therefore allow, for each score version separately,

$$\operatorname{Cov}(\varepsilon_{\mathrm{m}}^{(k)}, \varepsilon_{\mathrm{p}}^{(k)}) = \psi_{k,\mathrm{mp}}, \tag{4.7}$$

$$\operatorname{Cov}(\varepsilon_{\mathrm{m}}^{(k)}, \varepsilon_{\mathrm{c}}^{(k)}) = \psi_{k,\mathrm{mc}}, \tag{4.8}$$

$$\operatorname{Cov}(\varepsilon_{\mathrm{p}}^{(k)}, \varepsilon_{\mathrm{c}}^{(k)}) = \psi_{k,\mathrm{pc}}. \tag{4.9}$$

These are covariances on the observed score scale.

**Assumption A 1** (independent score versions)**.** The two observed score versions are constructed from independent weight perturbations, so that residual covariances across score versions vanish after conditioning on the visible-scale liability:

$$\operatorname{Cov}(\varepsilon_r^{(1)}, \varepsilon_s^{(2)}) = 0, \qquad \text{for all roles } r, s. \tag{4.10}$$

Using Equation (4.6), the corresponding residual correlations are

$$\operatorname{Corr}(\varepsilon_{\mathrm{m}}^{(k)}, \varepsilon_{\mathrm{c}}^{(k)}) = \frac{\psi_{k,\mathrm{mc}}}{\sqrt{\theta_{k\mathrm{m}}\theta_{k\mathrm{c}}}}, \qquad \operatorname{Corr}(\varepsilon_{\mathrm{p}}^{(k)}, \varepsilon_{\mathrm{c}}^{(k)}) = \frac{\psi_{k,\mathrm{pc}}}{\sqrt{\theta_{k\mathrm{p}}\theta_{k\mathrm{c}}}}. \tag{4.11}$$

A useful benchmark for these residual correlations follows directly from the shared-weight score-construction mechanism. Let the unstandardized score for role $r$ and score version $k$ be

$$\widetilde{P}_r^{(k)} = \sum_{j=1}^{M} x_{jr} w_j^{(k)}, \tag{4.12}$$

where $x_{jr}$ is the centered additive genotype at visible locus $j$. Write the score weight as

$$w_j^{(k)} = \gamma_j + \xi_j^{(k)}, \tag{4.13}$$

where $\gamma_j$ is the true visible effect size and $\xi_j^{(k)}$ is a mean-zero weight-noise term common to all family members for score version $k$. The corresponding raw-score residual contribution is

$$\tilde{\varepsilon}_r^{(k)} = \sum_{j=1}^{M} x_{jr} \xi_j^{(k)}. \tag{4.14}$$

Assume the weight-noise terms are independent across loci with common variance $\sigma^2_{w_k}$. Then

$$\begin{aligned}\operatorname{Cov}\left(\tilde{\varepsilon}_{\mathrm{c}}^{(k)}, \tilde{\varepsilon}_{\mathrm{m}}^{(k)}\right) &= \sum_{j=1}^{M} \operatorname{Cov}\left(x_{j\mathrm{c}}\xi_j^{(k)}, x_{j\mathrm{m}}\xi_j^{(k)}\right) \\ &= \sigma^2_{w_k} \sum_{j=1}^{M} \operatorname{Cov}(x_{j\mathrm{c}}, x_{j\mathrm{m}}). \end{aligned} \tag{4.15}$$

Under additive Mendelian inheritance,

$$\mathrm{E}(x_{j\mathrm{c}} \mid x_{j\mathrm{m}}, x_{j\mathrm{p}}) = \frac{x_{j\mathrm{m}} + x_{j\mathrm{p}}}{2}. \tag{4.16}$$

Consequently, allowing for spouse covariance at the locus,

$$\begin{aligned}\mathrm{Cov}(x_{j\mathrm{c}}, x_{j\mathrm{m}}) &= \frac{1}{2}\left\{\mathrm{Var}(x_{j\mathrm{m}}) + \mathrm{Cov}(x_{j\mathrm{p}}, x_{j\mathrm{m}})\right\}, \\ \mathrm{Cov}(x_{j\mathrm{c}}, x_{j\mathrm{p}}) &= \frac{1}{2}\left\{\mathrm{Var}(x_{j\mathrm{p}}) + \mathrm{Cov}(x_{j\mathrm{m}}, x_{j\mathrm{p}})\right\}. \end{aligned} \tag{4.17}$$

The same shared-weight calculation therefore gives

$$\begin{aligned}\mathrm{Cov}\left(\tilde{\varepsilon}_{\mathrm{c}}^{(k)}, \tilde{\varepsilon}_{\mathrm{m}}^{(k)}\right) &= \frac{1}{2}\left\{\mathrm{Var}\left(\tilde{\varepsilon}_{\mathrm{m}}^{(k)}\right) + \mathrm{Cov}\left(\tilde{\varepsilon}_{\mathrm{p}}^{(k)}, \tilde{\varepsilon}_{\mathrm{m}}^{(k)}\right)\right\}, \\ \mathrm{Cov}\left(\tilde{\varepsilon}_{\mathrm{c}}^{(k)}, \tilde{\varepsilon}_{\mathrm{p}}^{(k)}\right) &= \frac{1}{2}\left\{\mathrm{Var}\left(\tilde{\varepsilon}_{\mathrm{p}}^{(k)}\right) + \mathrm{Cov}\left(\tilde{\varepsilon}_{\mathrm{m}}^{(k)}, \tilde{\varepsilon}_{\mathrm{p}}^{(k)}\right)\right\}. \end{aligned} \tag{4.18}$$

Thus the familiar benchmark

$$\mathrm{Corr}\left(\tilde{\varepsilon}_{\mathrm{c}}^{(k)}, \tilde{\varepsilon}_{\mathrm{m}}^{(k)}\right) = \frac{1}{2}, \qquad \mathrm{Corr}\left(\tilde{\varepsilon}_{\mathrm{c}}^{(k)}, \tilde{\varepsilon}_{\mathrm{p}}^{(k)}\right) = \frac{1}{2} \tag{4.19}$$

is exact when the residual genetic score has equal variance across the three roles and zero spouse residual covariance. Write $X_r = (x_{1r}, \ldots, x_{Mr})^\top$. For a realized weight-noise vector $\xi^{(k)}$, the within-score spouse residual covariance is

$$\mathrm{Cov}\big(\tilde{\varepsilon}_{\mathrm{m}}^{(k)}, \tilde{\varepsilon}_{\mathrm{p}}^{(k)} \mid \xi^{(k)}\big) = (\xi^{(k)})^\top \,\mathrm{Cov}(X_{\mathrm{m}}, X_{\mathrm{p}})\xi^{(k)}. \tag{4.20}$$

If independent mean-zero weight perturbations with common variance $\sigma_{w_k}^2$ are averaged over, the expectation of Equation (4.20) is $\sigma_{w_k}^2 \sum_j \mathrm{Cov}(x_{j\mathrm{m}}, x_{j\mathrm{p}})$. It is zero under the benchmark in which the fitted residual score direction is orthogonal to the liability on which mating occurs and mates have no additional biological relatedness, endogamy, or other residual genetic similarity. It can be nonzero when those conditions fail. For that reason the general RAVEL SEM retains $\psi_{k,\mathrm{mp}}$ as an estimable within-score cross-spouse residual covariance even though the main-text benchmark fixes it to zero. Equation (4.18) also shows that the parent–offspring residual correlation need not equal one half when spouse residual covariance is nonzero or residual variances differ across roles. The section on Mendelian imputation shows that imputation changes the paternal-side benchmark even under the zero-spouse-residual condition.

#### 4.3 Visible trio covariance structure

The visible-scale trio liabilities are summarized by

$$\mathrm{Corr}(G_{\mathrm{m}}, G_{\mathrm{p}}) = \rho_G, \qquad \mathrm{Cov}(G_{\mathrm{c}}, G_{\mathrm{m}}) = \eta_{\mathrm{m}}, \qquad \mathrm{Cov}(G_{\mathrm{c}}, G_{\mathrm{p}}) = \eta_{\mathrm{p}}. \tag{4.21}$$

In the fitted SEM, where the visible liabilities are standardized to unit variance, the corresponding spouse covariance is numerically identical to $\rho_G$; we nevertheless refer to $\rho_G$ as a correlation throughout. The fitted visible-scale covariance block is agnostic about whether this genotypic spouse correlation arises from primary phenotypic assortment, social homogamy, ancestry-related matching, or another mechanism; the key requirement for the score-residual derivations above is only that the weight noise terms $\xi^{(k)}$ are independent of the visible genetic signal itself. The post-estimation total-liability correction however requires that the cross-mate covariance of the visible and hidden components is proportional to their covariance with the total liability. The quantities $\eta_{\mathrm{m}}$ and $\eta_{\mathrm{p}}$ are offspring–parent covariance parameters. Under the SEM standardization they are numerically also correlations, and they summarize Mendelian transmission together with any inflation of offspring–parent covariance induced by assortative mating.

The visible trio covariance matrix therefore takes the form

$$\Sigma_{GG} = \begin{pmatrix} 1 & \eta_{\mathrm{m}} & \eta_{\mathrm{p}} \\ \eta_{\mathrm{m}} & 1 & \rho_G \\ \eta_{\mathrm{p}} & \rho_G & 1 \end{pmatrix}. \tag{4.22}$$

#### 4.4 Visible-scale phenotype model

For a continuous phenotype,

$$Y = b_D G_{\mathrm{c}} + b_M G_{\mathrm{m}} + b_P G_{\mathrm{p}} + \gamma^\top W + \varepsilon_Y, \tag{4.23}$$

where $W$ denotes fixed effect covariates. For binary outcomes, the same linear predictor is used on an unobserved probit liability scale. Let

$$b = (b_D, b_M, b_P)^\top. \tag{4.24}$$

Because the trio latent liabilities are correlated, the visible-scale genetic variance captured by the model is

$$\sigma^2_{G,\mathrm{vis}} = b^\top \Sigma_{GG} b. \tag{4.25}$$

Expanding Equation (4.25) gives

$$\begin{aligned} \sigma^2_{G,\mathrm{vis}} = {} & b_D^2 + b_M^2 + b_P^2 \\ & + 2b_D b_M \eta_{\mathrm{m}} + 2b_D b_P \eta_{\mathrm{p}} + 2b_M b_P \rho_G. \end{aligned} \tag{4.26}$$

For a continuous phenotype with residual variance $\sigma_Y^2$, the corresponding visible-scale explained proportion is

$$R^2_{\mathrm{vis}} = \frac{\sigma^2_{G,\mathrm{vis}}}{\sigma^2_{G,\mathrm{vis}} + \sigma_Y^2}. \tag{4.27}$$

### 5 Projection and transmission coefficients

#### 5.1 Projection from visible-scale coefficients to total-liability coefficients

Suppose the phenotype is generated by the broader standardized total-liability model

$$Y = \beta_D T_{\mathrm{c}} + \beta_M T_{\mathrm{m}} + \beta_P T_{\mathrm{p}} + \gamma^\top W + \varepsilon_Y, \tag{5.1}$$

where $T_r = T_r^*/\sqrt{V_{T,r}}$ as in Equation (2.7). Thus $\beta_D$, $\beta_M$, and $\beta_P$ are coefficients on role-standardized total liabilities, matching the coefficient scale used by the analytic expectations in the main text. Define

$$\beta = \left(\beta_D, \beta_M, \beta_P\right)^\top. \tag{5.2}$$

Then Equation (5.1) may be written compactly as

$$Y = \beta^\top T + \gamma^\top W + \varepsilon_Y \tag{5.3}$$

and we assume

$$\mathrm{Cov}(G, \varepsilon_Y \mid W) = 0.$$

.

All covariance operators in this section and in the hidden-correction derivations below are understood as partial covariances after linear residualization on the fixed effect covariates $W$ whenever such covariates are included.

Let

$$\begin{aligned} D_G &= \mathrm{diag}\left(\sqrt{V_{G,\mathrm{c}}}, \sqrt{V_{G,\mathrm{m}}}, \sqrt{V_{G,\mathrm{p}}}\right), \\ D_H &= \mathrm{diag}\left(\sqrt{V_{H,\mathrm{c}}}, \sqrt{V_{H,\mathrm{m}}}, \sqrt{V_{H,\mathrm{p}}}\right), \\ D_T &= \mathrm{diag}\left(\sqrt{V_{T,\mathrm{c}}}, \sqrt{V_{T,\mathrm{m}}}, \sqrt{V_{T,\mathrm{p}}}\right). \end{aligned} \tag{5.4}$$

From Equations (2.5) and (2.7),

$$T^* = \omega_G D_G G + \omega_H D_H H, \qquad T = D_T^{-1} T^*. \tag{5.5}$$

By definition of the population projection of $Y$ onto $G$,

$$b = \Sigma_{GG}^{-1}\,\mathrm{Cov}(G, Y). \tag{5.6}$$

Using Equation (5.3),

$$\begin{aligned} \mathrm{Cov}(G, Y) &= \mathrm{Cov}\left(G, \beta^\top T\right) \\ &= \mathrm{Cov}(G, T)\,\beta. \end{aligned} \tag{5.7}$$

Equation (5.5) gives

$$\mathrm{Cov}(G,T) = (\omega_G \Sigma_{GG} D_G + \omega_H \Sigma_{GH} D_H)\, D_T^{-1}. \tag{5.8}$$

Substituting Equation (5.8) into Equation (5.6) gives

$$b = \left[\omega_G D_G + \omega_H \Sigma_{GG}^{-1} \Sigma_{GH} D_H\right] D_T^{-1} \beta. \tag{5.9}$$

Therefore,

$$\boxed{\beta = D_T \left[\omega_G D_G + \omega_H \Sigma_{GG}^{-1} \Sigma_{GH} D_H\right]^{-1} b.} \tag{5.10}$$

The hidden variant problem is thus reduced to construction of the role-specific raw variances and the standardized cross-block $\Sigma_{GH}$.

For completeness, if coefficients on the raw total liabilities $T^*$ are desired, define $Y = (\beta^{\mathrm{raw}})^\top T^* + \gamma^\top W + \varepsilon_Y$. Then

$$\beta^{\mathrm{raw}} = \left[\omega_G D_G + \omega_H \Sigma_{GG}^{-1} \Sigma_{GH} D_H\right]^{-1} b, \qquad \beta = D_T \beta^{\mathrm{raw}}. \tag{5.11}$$

Thus Equation (5.10), rather than a later ad hoc rescaling, directly returns coefficients on the standardized total-liability scale used in the main text.

If visible and hidden components are orthogonal throughout the trio, so that $\Sigma_{GH} = 0$, Equation (5.10) reduces to

$$\beta = \frac{1}{\sqrt{1-\pi}} D_T D_G^{-1} b. \tag{5.12}$$

Under random mating, $D_T = D_G = I_3$ and this further reduces to the simpler $\beta = b/\sqrt{1-\pi}$. During transient assortment, the role-specific variance ratios remain even when the visible–hidden cross-block is set to zero.

#### 5.2 Recovery of the visible transmission coefficients

The child's standardized visible factor can be represented as the population projection on the parental standardized visible factors,

$$G_{\mathrm{c}} = t_{\mathrm{m}} G_{\mathrm{m}} + t_{\mathrm{p}} G_{\mathrm{p}} + u_G, \tag{5.13}$$

with

$$\mathrm{Cov}(u_G, G_{\mathrm{m}}) = 0, \qquad \mathrm{Cov}(u_G, G_{\mathrm{p}}) = 0. \tag{5.14}$$

Taking covariances of Equation (5.13) with the parental visible factors yields the normal equations. First,

$$\begin{aligned}
\mathrm{Cov}(G_{\mathrm{c}}, G_{\mathrm{m}}) &= \mathrm{Cov}(t_{\mathrm{m}} G_{\mathrm{m}} + t_{\mathrm{p}} G_{\mathrm{p}} + u_G,\, G_{\mathrm{m}}) \\
&= t_{\mathrm{m}} \,\mathrm{Var}(G_{\mathrm{m}}) + t_{\mathrm{p}} \,\mathrm{Cov}(G_{\mathrm{p}}, G_{\mathrm{m}}) + \mathrm{Cov}(u_G, G_{\mathrm{m}}) \\
&= t_{\mathrm{m}} + \rho_G t_{\mathrm{p}}.
\end{aligned} \tag{5.15}$$

Because $\mathrm{Cov}(G_c, G_m) = \eta_m$, Equation (5.15) becomes

$$\eta_m = t_m + \rho_G t_p. \tag{5.16}$$

Similarly,

$$\eta_p = \rho_G t_m + t_p. \tag{5.17}$$

Equivalently,

$$\begin{pmatrix} 1 & \rho_G \\ \rho_G & 1 \end{pmatrix} \begin{pmatrix} t_m \\ t_p \end{pmatrix} = \begin{pmatrix} \eta_m \\ \eta_p \end{pmatrix}, \tag{5.18}$$

so that

$$t_m = \frac{\eta_m - \rho_G \eta_p}{1 - \rho_G^2}, \qquad t_p = \frac{\eta_p - \rho_G \eta_m}{1 - \rho_G^2}. \tag{5.19}$$

These are standardized visible-scale projection coefficients implied by the fitted trio covariance block.

**Assumption A 2** (common raw additive transmission across visible and hidden variants)**.** On the fixed founder scale, the additive components satisfy

$$G_c^* = \tfrac{1}{2} G_m^* + \tfrac{1}{2} G_p^* + U_G, \qquad H_c^* = \tfrac{1}{2} H_m^* + \tfrac{1}{2} H_p^* + U_H. \tag{5.20}$$

Under the infinitesimal, constant genic variance approximation used for the main text expectations,

$$\mathrm{Var}(U_G) = \mathrm{Var}(U_H) = \tfrac{1}{2}, \qquad \mathrm{Cov}(U_G, U_H) = 0, \tag{5.21}$$

and the segregation residuals are orthogonal to all parental visible and hidden components,

$$\mathrm{Cov}(U_G, G_m^*) = \mathrm{Cov}(U_G, G_p^*) = \mathrm{Cov}(U_G, H_m^*) = \mathrm{Cov}(U_G, H_p^*) = 0, \tag{5.22}$$

with the analogous conditions for $U_H$.

After role-specific standardization, the raw one half coefficients in Equation (5.20) become

$$t_{G,m} = \frac{1}{2}\sqrt{\frac{V_{G,m}}{V_{G,c}}}, \qquad t_{G,p} = \frac{1}{2}\sqrt{\frac{V_{G,p}}{V_{G,c}}}, \tag{5.23}$$

and

$$t_{H,m} = \frac{1}{2}\sqrt{\frac{V_{H,m}}{V_{H,c}}}, \qquad t_{H,p} = \frac{1}{2}\sqrt{\frac{V_{H,p}}{V_{H,c}}}. \tag{5.24}$$

For complete additive trios, the $t_m, t_p$ recovered from Equation (5.19) correspond to $t_{G,m}, t_{G,p}$. The visible and hidden standardized coefficients are generally different during transient assortment because $V_G$ and $V_H$ inflate by different amounts. They both equal one half under random mating and at equilibrium, where parent and child component variances are equal.

#### 5.3 Transmission coefficients under non-additive scoring schemes

The transmission coefficients $t_{\mathrm{m}}$ and $t_{\mathrm{p}}$ in Equation (5.19) are population projection coefficients on the fitted visible scale. They should therefore not be interpreted mechanically as biological segregation probabilities. In complete trios with directly observed additive genotypes and a shared additive score construction, fixing the coefficients at one half is appropriate on the fixed raw genetic scale. On the role-standardized SEM scale, the corresponding coefficients are those in Equation (5.23); they equal one half only when the relevant parent and child raw variances are the same.

For an additive genotype $X_{jr} \in \{0, 1, 2\}$,

$$\mathrm{E}(X_{j\mathrm{c}} \mid X_{j\mathrm{m}}, X_{j\mathrm{p}}) = \frac{X_{j\mathrm{m}} + X_{j\mathrm{p}}}{2}. \tag{5.25}$$

Consequently, for an additive score

$$S_r = \sum_j w_j \big(X_{jr} - 2f_j\big), \tag{5.26}$$

one has

$$\mathrm{E}(S_{\mathrm{c}} \mid \mathbf{X}_{\mathrm{m}}, \mathbf{X}_{\mathrm{p}}) = \frac{1}{2} S_{\mathrm{m}} + \frac{1}{2} S_{\mathrm{p}}. \tag{5.27}$$

This is the sense in which the additive real-trio benchmark gives one-half maternal and paternal transmission on the raw score scale.

The same conclusion does not hold automatically for Mendelian-imputed scores or for score systems that are not linear additive functions of the directly observed parental genotypes. More generally, suppose the score is defined by a per-locus genotype-scoring function $\varphi_j$,

$$S_r = \sum_j w_j \varphi_j(X_{jr}). \tag{5.28}$$

Let $A_{j\mathrm{m}}$ and $A_{j\mathrm{p}}$ denote the maternally and paternally transmitted alleles at locus $j$. Conditional on a parental genotype $X$, the transmitted allele satisfies

$$\Pr(A = 1 \mid X) = \frac{X}{2}, \qquad \Pr(A = 0 \mid X) = 1 - \frac{X}{2}.$$

Therefore,

$$\mathrm{E}(S_{\mathrm{c}} \mid \mathbf{X}_{\mathrm{m}}, \mathbf{X}_{\mathrm{p}}) = \sum_j w_j \sum_{a=0}^{1} \sum_{b=0}^{1} \varphi_j(a+b) \Pr(A_{j\mathrm{m}} = a \mid X_{j\mathrm{m}}) \Pr(A_{j\mathrm{p}} = b \mid X_{j\mathrm{p}}). \tag{5.29}$$

Only for linear additive scoring does Equation (5.29) reduce to the midparent expression in Equation (5.27).

For example, consider a recessive score

$$S_r^{\mathrm{rec}} = \sum_j w_j \mathbb{1}\{X_{jr} = 2\}. \tag{5.30}$$

At a single locus,

$$\mathrm{E}(\mathbb{1}\{X_{j\mathrm{c}} = 2\} \mid X_{j\mathrm{m}}, X_{j\mathrm{p}}) = \frac{X_{j\mathrm{m}}}{2}\frac{X_{j\mathrm{p}}}{2} = \frac{X_{j\mathrm{m}}X_{j\mathrm{p}}}{4}. \tag{5.31}$$

This is not a linear function of the parental recessive indicators. If both parents are heterozygous, the parental recessive indicators are both zero and the midparent recessive score is zero, but the expected offspring recessive indicator is 1/4. Conversely, if one parent is homozygous for the effect allele and the other is homozygous reference, the midparent recessive indicator is 1/2, but the expected offspring recessive indicator is zero.

These examples show that fixing $t_\mathrm{m} = t_\mathrm{p} = 1/2$ is a raw-score-specific assumption. For a non-additive or imputed score, the visible-scale projection coefficients may be estimated from the fitted covariance matrix using Equation (5.19), or calculated from the actual score definition and parental genotype data. This is sufficient for interpretation of the visible-scale SEM. It is not, by itself, sufficient to justify the additive hidden component correction below. Recessive, dominance, epistatic, and other non-additive score systems require a score-specific raw variance and cross-component transmission model before Equation (5.10) can be given the same total-liability interpretation. De novo variation similarly requires a separate model.

Substantively, this also affects the interpretation of the direct offspring coefficient. For additive complete trio scores, the offspring raw visible component can be viewed as the transmitted deviation from a midparent expectation plus Mendelian sampling. For non-additive and imputed scores, the expected offspring score is generally not the midparent score. The direct coefficient should therefore be interpreted as the association of the offspring visible liability conditional on the parental visible liabilities in the fitted score system, rather than automatically as an effect of deviation from the parental midpoint.

### 6 Cross-component covariance under assortment on the total liability

Let

$$Z_g = \begin{pmatrix} G_g^* \\ H_g^* \end{pmatrix}, \qquad w = \begin{pmatrix} \omega_G \\ \omega_H \end{pmatrix}, \qquad T_g^* = w^\top Z_g, \tag{6.1}$$

and write $S_g = \mathrm{Cov}(Z_g)$ and $V_g = \mathrm{Var}(T_g^*) = w^\top S_g w$. The founder conditions in Equation (2.6) imply $S_0 = I_2$ and $V_0 = 1$.

**Assumption A 3** (common assortment pattern across visible and hidden components)**.** For adults in generation $g$, let

$$c_g = \mathrm{Cov}(Z_g, T_g^*) = S_g w, \qquad V_g = w^\top S_g w.$$

The aggregate cross-mate covariance within the modeled $(G^*, H^*)$ component space is assumed to have the single direction $c_g$:

$$\mathrm{Cov}(Z_{\mathrm{m},g}, Z_{\mathrm{p},g}) = \rho_{T,g}\frac{c_g c_g^\top}{V_g} = \rho_{T,g}\frac{S_g w w^\top S_g}{w^\top S_g w}, \tag{6.2}$$

where $\rho_{T,g} = \mathrm{Corr}(T_{\mathrm{m},g}^*, T_{\mathrm{p},g}^*)$.

This is a restriction on the projection of the complete mate-choice process into the modeled visible–hidden component space; it does not require $T_g^*$ to be the only trait involved in mate choice. An additional assorting axis $A_g$ is compatible with the assumption whenever

$$\mathrm{Cov}(Z_g, A_g) = \frac{\mathrm{Cov}(T_g^*, A_g)}{V_g}\,\mathrm{Cov}(Z_g, T_g^*),$$

or equivalently,

$$\mathrm{Cov}\left(Z_g,\, A_g - \frac{\mathrm{Cov}(A_g, T_g^*)}{V_g} T_g^*\right) = 0.$$

Thus additional axes may affect pairing, but after linear residualization on $T_g^*$ they have no further covariance with the modeled visible or hidden component. This proportionality is what allows the scalar $\rho_{T,g}$ to determine the component-specific cross-mate covariance block.

**Assumption A 4** (parental exchangeability of raw component moments)**.** Within a generation, maternal and paternal marginal raw covariance matrices are equal, and the cross-mate covariance matrix in Equation (6.2) is symmetric under exchange of maternal and paternal roles.

#### 6.1 One generation of assortment

We first derive the visible–hidden cross-component covariance matrix under one generation of assortment. The adults who mate are in the founder state,

$$S_0 = I_2, \qquad \mathrm{Cov}(G^*_{\mathrm{m},0}, H^*_{\mathrm{m},0}) = \mathrm{Cov}(G^*_{\mathrm{p},0}, H^*_{\mathrm{p},0}) = 0. \tag{6.3}$$

Under Assumption 3, $S_0 w = w$ and the raw cross-mate covariance block is

$$K_0 := \mathrm{Cov}(Z_{\mathrm{m},0}, Z_{\mathrm{p},0}) = \rho_T w w^\top = \rho_T \begin{pmatrix} 1-\pi & \sqrt{\pi(1-\pi)} \\ \sqrt{\pi(1-\pi)} & \pi \end{pmatrix}. \tag{6.4}$$

In particular,

$$K_{GG,0} = (1-\pi)\rho_T, \tag{6.5}$$

$$K_{GH,0} = \sqrt{\pi(1-\pi)}\,\rho_T, \tag{6.6}$$

$$K_{HH,0} = \pi\rho_T. \tag{6.7}$$

Because the founder component variances are one, the standardized visible spouse correlation is

$$\rho_G = (1-\pi)\rho_T, \qquad \rho_T = \frac{\rho_G}{1-\pi}, \tag{6.8}$$

and the standardized cross-spouse visible–hidden covariance is

$$q = K_{GH,0} = \sqrt{\pi(1-\pi)}\,\rho_T = \rho_G\sqrt{\frac{\pi}{1-\pi}}. \tag{6.9}$$

Using the raw additive transmission model in Assumption 2,

$$S_1 = \frac{1}{2}S_0 + \frac{1}{2}K_0 + \frac{1}{2}I_2 = I_2 + \frac{\rho_T}{2}ww^\top. \tag{6.10}$$

Hence

$$V_1 = 1 + \frac{\rho_T}{2}, \tag{6.11}$$

with raw child component moments

$$V_{G,1} = 1 + \frac{(1-\pi)\rho_T}{2}, \tag{6.12}$$

$$V_{H,1} = 1 + \frac{\pi\rho_T}{2}, \tag{6.13}$$

$$C_{GH,1} := \mathrm{Cov}(G_1^*, H_1^*) = \frac{\sqrt{\pi(1-\pi)}\rho_T}{2}. \tag{6.14}$$

Let $L_{GH,0} = \frac{1}{2}K_{GH,0}$ denote either raw offspring–parent visible–hidden covariance. In child–mother–father order, the raw cross-block is

$$\Sigma^{(1)}_{G^*H^*} = \begin{pmatrix} C_{GH,1} & L_{GH,0} & L_{GH,0} \\ L_{GH,0} & 0 & K_{GH,0} \\ L_{GH,0} & K_{GH,0} & 0 \end{pmatrix}. \tag{6.15}$$

The standardized block used in Equation (5.10) is

$$\Sigma^{(1)}_{GH} = D_G^{-1}\Sigma^{(1)}_{G^*H^*}D_H^{-1}, \tag{6.16}$$

where $D_G = \mathrm{diag}(\sqrt{V_{G,1}}, 1, 1)$ and $D_H = \mathrm{diag}(\sqrt{V_{H,1}}, 1, 1)$. Although the raw block in Equation (6.15) is symmetric, the role-standardized block need not be symmetric because the child visible and hidden variances are standardized by different factors.

#### 6.2 $n$-generation recursion

The one-generation model is generally insufficient when assortative mating has acted for multiple generations. In that case the raw visible and hidden variances inflate, and the two components become correlated within individuals as well as across spouses.

Under constant $\rho_T$, suppose inductively that

$$S_g = I_2 + (V_g - 1)ww^\top. \tag{6.17}$$

Then $S_g w = V_g w$, and Assumption 3 gives

$$K_g := \mathrm{Cov}(Z_{\mathrm{m},g}, Z_{\mathrm{p},g}) = \rho_T V_g ww^\top. \tag{6.18}$$

Raw additive transmission gives

$$
\begin{aligned}
S_{g+1} &= \frac{1}{2}S_g + \frac{1}{2}K_g + \frac{1}{2}I_2 \\
&= I_2 + \frac{(1+\rho_T)V_g - 1}{2} ww^\top. \qquad (6.19)
\end{aligned}
$$

Thus the form in Equation (6.17) is preserved and the total raw variance is

$$V_{g+1} = \frac{1+\rho_T}{2}V_g + \frac{1}{2}, \qquad V_0 = 1. \qquad (6.20)$$

Define

$$\alpha = \frac{1+\rho_T}{2}, \qquad V_\infty = \frac{1}{1-\rho_T}. \qquad (6.21)$$

The finite generation solution is

$$V_g = V_\infty + (1 - V_\infty)\alpha^g. \qquad (6.22)$$

Expanding Equation (6.17) gives the raw component moments

$$V_{G,g} = \pi + (1-\pi)V_g, \qquad (6.23)$$

$$V_{H,g} = (1-\pi) + \pi V_g, \qquad (6.24)$$

$$C_{GH,g} := \mathrm{Cov}(G_g^*, H_g^*) = \sqrt{\pi(1-\pi)}\,(V_g - 1). \qquad (6.25)$$

These identities imply exactly

$$(1-\pi)V_{G,g} + \pi V_{H,g} + 2\sqrt{\pi(1-\pi)}\,C_{GH,g} = V_g. \qquad (6.26)$$

Thus AM-induced inflation is not represented only by a changing visible–hidden covariance, the marginal raw variances of both components change as well.

For an observed trio produced after $n \geq 1$ mating rounds, let

$$V_{\mathrm{par}} = V_{n-1}, \qquad V_{\mathrm{c}} = V_n, \qquad (6.27)$$

and define $V_{G,\mathrm{par}}, V_{H,\mathrm{par}}, C_{GH,\mathrm{par}}$ and the child quantities by Equations (6.23)–(6.25). The raw component-specific spouse covariances in the focal parental generation are

$$K_{GG} = \rho_T(1-\pi)V_{\mathrm{par}}, \qquad (6.28)$$

$$K_{GH} = \rho_T\sqrt{\pi(1-\pi)}V_{\mathrm{par}}, \qquad (6.29)$$

$$K_{HH} = \rho_T \pi V_{\mathrm{par}}. \qquad (6.30)$$

The corresponding raw offspring–parent covariances are

$$L_{GG} = \frac{1}{2}\left(V_{G,\text{par}} + K_{GG}\right), \tag{6.31}$$
$$L_{GH} = \frac{1}{2}\left(C_{GH,\text{par}} + K_{GH}\right), \tag{6.32}$$
$$L_{HH} = \frac{1}{2}\left(V_{H,\text{par}} + K_{HH}\right). \tag{6.33}$$

On the fitted standardized visible scale, these imply

$$\rho_G = \frac{K_{GG}}{V_{G,\text{par}}} = \rho_T \frac{(1-\pi)V_{\text{par}}}{V_{G,\text{par}}}, \qquad \eta_G = \frac{L_{GG}}{\sqrt{V_{G,\text{c}}V_{G,\text{par}}}}. \tag{6.34}$$

Thus, for specified $\pi$ and AM duration, a fitted $\rho_G$ determines the corresponding $\rho_T$ through Equation (6.34) together with the variance recursion. In child–mother–father order, the raw covariance blocks are

$$\Sigma_{G^*G^*} = \begin{pmatrix} V_{G,\text{c}} & L_{GG} & L_{GG} \\ L_{GG} & V_{G,\text{par}} & K_{GG} \\ L_{GG} & K_{GG} & V_{G,\text{par}} \end{pmatrix}, \tag{6.35}$$

$$\Sigma_{G^*H^*} = \begin{pmatrix} C_{GH,\text{c}} & L_{GH} & L_{GH} \\ L_{GH} & C_{GH,\text{par}} & K_{GH} \\ L_{GH} & K_{GH} & C_{GH,\text{par}} \end{pmatrix}, \tag{6.36}$$

and

$$\Sigma_{H^*H^*} = \begin{pmatrix} V_{H,\text{c}} & L_{HH} & L_{HH} \\ L_{HH} & V_{H,\text{par}} & K_{HH} \\ L_{HH} & K_{HH} & V_{H,\text{par}} \end{pmatrix}. \tag{6.37}$$

Let

$$\begin{aligned} D_G &= \operatorname{diag}\left(\sqrt{V_{G,\text{c}}}, \sqrt{V_{G,\text{par}}}, \sqrt{V_{G,\text{par}}}\right), \\ D_H &= \operatorname{diag}\left(\sqrt{V_{H,\text{c}}}, \sqrt{V_{H,\text{par}}}, \sqrt{V_{H,\text{par}}}\right), \\ D_T &= \operatorname{diag}\left(\sqrt{V_{\text{c}}}, \sqrt{V_{\text{par}}}, \sqrt{V_{\text{par}}}\right). \end{aligned} \tag{6.38}$$

The covariance blocks on the fitted role-standardized scale are

$$\begin{aligned} \Sigma_{GG} &= D_G^{-1}\Sigma_{G^*G^*}D_G^{-1}, \\ \Sigma_{GH} &= D_G^{-1}\Sigma_{G^*H^*}D_H^{-1}, \\ \Sigma_{HH} &= D_H^{-1}\Sigma_{H^*H^*}D_H^{-1}. \end{aligned} \tag{6.39}$$

These matrices, inserted into Equation (5.10), define the exact finite generation correction under the maintained raw infinitesimal model.

The standardized visible and hidden transmission coefficients induced by the same raw one half inheritance are

$$t_G = \frac{1}{2}\sqrt{\frac{V_{G,\text{par}}}{V_{G,\text{c}}}}, \qquad t_H = \frac{1}{2}\sqrt{\frac{V_{H,\text{par}}}{V_{H,\text{c}}}}. \tag{6.40}$$

They are generally unequal in a transient generation. This is why a common standardized $t_\text{m}, t_\text{p}$ cannot be imposed simultaneously on $G$ and $H$ in the finite generation raw model.

##### 6.2.1 Closed-form expected coefficients used in the main text

The displayed scalar expectations in the main text are the complete additive trio case of Equation (5.9). Let $\beta_D, \beta_M, \beta_P$ denote coefficients on the role-standardized total liabilities, let $V_\text{par}$ and $V_\text{c}$ be as in Equation (6.27), and write

$$V_{G,\text{par}} = \pi + (1-\pi)V_\text{par}, \qquad V_{G,\text{c}} = \pi + (1-\pi)V_\text{c}. \tag{6.41}$$

Substitution of Equations (6.35)–(6.39) into Equation (5.9) yields

$$\beta_{\text{vis},D} = \sqrt{\frac{(1-\pi)V_{G,\text{c}}}{V_\text{c}}}\,\beta_D. \tag{6.42}$$

Define

$$A_D = \frac{\pi\sqrt{(1-\pi)V_{G,\text{par}}}\,(V_\text{c}-1)}{\sqrt{V_\text{c}}\,\{V_{G,\text{par}} + \rho_T(1-\pi)V_\text{par}\}}, \tag{6.43}$$

$$A_+ = \frac{(1+\rho_T)\sqrt{(1-\pi)V_\text{par}V_{G,\text{par}}}}{2\,\{V_{G,\text{par}} + \rho_T(1-\pi)V_\text{par}\}}, \tag{6.44}$$

and

$$A_- = \frac{(1-\rho_T)\sqrt{(1-\pi)V_\text{par}V_{G,\text{par}}}}{2\,\{V_{G,\text{par}} - \rho_T(1-\pi)V_\text{par}\}}. \tag{6.45}$$

The expected maternal and paternal visible-scale coefficients are then

$$\beta_{\text{vis},M} = A_D\beta_D + A_+(\beta_M + \beta_P) + A_-(\beta_M - \beta_P), \tag{6.46}$$

$$\beta_{\text{vis},P} = A_D\beta_D + A_+(\beta_M + \beta_P) - A_-(\beta_M - \beta_P). \tag{6.47}$$

Here $\rho_T$ is specifically the standardized total-liability mate correlation. Equation (6.42) also makes explicit that direct coefficient attenuation generally depends on AM through $V_{G,\text{c}}/V_\text{c}$, in addition to its dependence on $\pi$.

The observed PGS shortcut used in the main text expectation plots is a corollary of this same model. If the same additive score weights are applied to all trio members, score weights error is inherited under the same additive raw process, that error is orthogonal to the true visible and hidden components, the phenotype residual, and mate selection, and $r_0$ is the common founder-scale reliability, then the fraction of the founder total liability captured by the observed score is $r_0(1-\pi)$. Hence

$$\pi_\text{PGS} = 1 - r_0(1-\pi). \tag{6.48}$$

Under these assumptions, evaluating Equations (6.42)–(6.47) at $\pi_\text{PGS}$ is equivalent to carrying the additive score-error component through the full covariance calculation.

##### 6.3 Arbitrary assortative mating histories and effective constant-history representations

The preceding recursion assumes that assortative mating has constant strength across generations. This is a useful parameterization, but the hidden variant correction does not depend on the full historical path directly. Under the raw model it depends on the parental raw total-liability variance accumulated from that path and on the current total-liability spouse correlation. This subsection formalizes that statement and clarifies when an arbitrary history can be represented as an effective number of generations of constant assortative mating at the present-day strength.

Let $\rho_{T,g}$ denote the standardized total-liability spouse correlation in generation $g$, and define

$$\alpha_g = \frac{1+\rho_{T,g}}{2}, \qquad \mathcal{V}_\rho(x) = \frac{1+\rho}{2}x + \frac{1}{2}. \tag{6.49}$$

For an arbitrary finite history

$$\mathcal{H}_n = (\rho_{T,0}, \rho_{T,1}, \ldots, \rho_{T,n}), \tag{6.50}$$

the first $n$ correlations generate the raw variance among the focal parents, and $\rho_{T,n}$ is the spouse correlation among those parents. Starting from $V_0 = 1$,

$$V_n(\mathcal{H}_n) = \mathcal{V}_{\rho_{T,n-1}} \circ \cdots \circ \mathcal{V}_{\rho_{T,0}}(1), \tag{6.51}$$

and the child variance is

$$V_{n+1}(\mathcal{H}_n) = \mathcal{V}_{\rho_{T,n}}(V_n(\mathcal{H}_n)). \tag{6.52}$$

The exact finite-history solution for the parental variance is

$$V_n = \left(\prod_{k=0}^{n-1} \alpha_k\right) V_0 + \frac{1}{2}\sum_{\ell=0}^{n-1} \prod_{k=\ell+1}^{n-1} \alpha_k, \tag{6.53}$$

where an empty product equals one. Equivalently, the AM-induced excess variance has the exact weighted representation

$$V_n - 1 = \frac{1}{2}\sum_{j=0}^{n-1} \rho_{T,j} \prod_{k=j+1}^{n-1} \alpha_k. \tag{6.54}$$

Equation (6.54) makes clear why recent assortment receives more weight than ancient assortment.

Given $V_n$ and the current correlation $\rho_{T,n}$, Equations (6.23)–(6.33) determine all raw component moments for the focal trio. Thus two histories that imply the same pair

$$\big(V_n(\mathcal{H}_n), \rho_{T,n}\big) \tag{6.55}$$

imply the same hidden component correction.

Let $\rho_{T,\mathrm{cur}} := \rho_{T,n}$, $\alpha = (1+\rho_{T,\mathrm{cur}})/2$, and $V_\infty = 1/(1-\rho_{T,\mathrm{cur}})$. A constant-current-strength effective duration $n_{\text{eff}}$ is any value satisfying

$$V_n(\mathcal{H}_n) = V_\infty + (1 - V_\infty)\alpha^{n_{\text{eff}}}. \tag{6.56}$$

For $0 < \rho_{T,\text{cur}} < 1$ and $1 \le V_n \le V_\infty$, the real-valued solution is

$$n_{\text{eff}} = \frac{\log\{(V_\infty - V_n)/(V_\infty - 1)\}}{\log \alpha}. \tag{6.57}$$

If $n_{\text{eff}}$ is restricted to integer generations, exact equality requires $V_n$ to lie on the discrete constant-strength orbit; otherwise Equation (6.57) defines the natural continuous interpolation.

The update $\mathcal{V}_\rho(x)$ is monotone nondecreasing in both $\rho$ and $x$. Therefore, if

$$0 \le \rho_{T,0} \le \rho_{T,1} \le \cdots \le \rho_{T,n} = \rho_{T,\text{cur}}, \tag{6.58}$$

then induction gives

$$V_g(\mathcal{H}_n) \le \mathcal{V}^g_{\rho_{T,\text{cur}}}(1), \qquad g = 0, \ldots, n. \tag{6.59}$$

Consequently $1 \le V_n \le V_\infty$, a constant-current-strength effective representation exists, and $0 \le n_{\text{eff}} \le n$. In words, a gradual increase in assortative mating is equivalent, for the purposes of the correction, to fewer generations of constant assortative mating at the present-day strength.

For arbitrary histories, the constant-current-strength representation need not exist. If assortment was stronger in the past than in the present, the accumulated parental variance can exceed the equilibrium value attainable under the current correlation,

$$V_n(\mathcal{H}_n) > V_\infty(\rho_{T,\text{cur}}). \tag{6.60}$$

In that case the correct summary is the effective state in Equation (6.55), which can be inserted directly into the finite generation covariance blocks above.

#### 6.4 Assortment at equilibrium

At equilibrium, Equation (6.20) gives

$$V_\infty = \frac{1}{1 - \rho_T}. \tag{6.61}$$

The raw component moments are

$$V_{G,\infty} = \pi + (1 - \pi)V_\infty, \tag{6.62}$$
$$V_{H,\infty} = (1 - \pi) + \pi V_\infty, \tag{6.63}$$
$$C_{GH,\infty} = \sqrt{\pi(1 - \pi)}(V_\infty - 1). \tag{6.64}$$

The raw cross-spouse visible–hidden covariance is

$$K_{GH,\infty} = \rho_T \sqrt{\pi(1 - \pi)} V_\infty = C_{GH,\infty}. \tag{6.65}$$

Define the standardized within-person and cross-spouse visible–hidden covariances

$$d = \frac{C_{GH,\infty}}{\sqrt{V_{G,\infty} V_{H,\infty}}}, \qquad q = \frac{K_{GH,\infty}}{\sqrt{V_{G,\infty} V_{H,\infty}}}. \tag{6.66}$$

Equation (6.65) implies

$$d = q. \tag{6.67}$$

The standardized visible spouse correlation at equilibrium is

$$\rho_G = \frac{\rho_T(1-\pi)V_\infty}{V_{G,\infty}} = \frac{\rho_T(1-\pi)}{1-\pi\rho_T}. \tag{6.68}$$

Equivalently,

$$\rho_T = \frac{\rho_G}{1-\pi+\pi\rho_G}. \tag{6.69}$$

The corresponding raw total-liability spouse covariance is

$$\kappa_T := \mathrm{Cov}(T^*_{\mathrm{m}}, T^*_{\mathrm{p}}) = \rho_T V_\infty = \frac{\rho_T}{1-\rho_T}. \tag{6.70}$$

Thus $\rho_T$ denotes a standardized spouse correlation and $\kappa_T$ the corresponding raw covariance.

#### 6.5 Equilibrium covariance matrices

At equilibrium, parent and child raw component variances are equal, so the standardized visible and hidden transmission coefficients both equal one half. The visible spouse correlation is $\rho_G$ from Equation (6.68), and the offspring–parent visible covariance is $(1+\rho_G)/2$. Therefore,

$$\Sigma_{GG}^{(\mathrm{eq})} = \begin{pmatrix} 1 & (1+\rho_G)/2 & (1+\rho_G)/2 \\ (1+\rho_G)/2 & 1 & \rho_G \\ (1+\rho_G)/2 & \rho_G & 1 \end{pmatrix}. \tag{6.71}$$

Because $C_{GH,\infty} = K_{GH,\infty}$ and the offspring–parent raw visible–hidden covariance is their average, every entry of the standardized cross-block is $d$:

$$\Sigma_{GH}^{(\mathrm{eq})} = d \begin{pmatrix} 1 & 1 & 1 \\ 1 & 1 & 1 \\ 1 & 1 & 1 \end{pmatrix}. \tag{6.72}$$

For completeness, define

$$\rho_H = \frac{\rho_T \pi V_\infty}{V_{H,\infty}} = \frac{\rho_T \pi}{1-(1-\pi)\rho_T}. \tag{6.73}$$

Then

$$\Sigma_{HH}^{(\mathrm{eq})} = \begin{pmatrix} 1 & (1+\rho_H)/2 & (1+\rho_H)/2 \\ (1+\rho_H)/2 & 1 & \rho_H \\ (1+\rho_H)/2 & \rho_H & 1 \end{pmatrix}. \tag{6.74}$$

At equilibrium, $D_G = \sqrt{V_{G,\infty}} I_3$, $D_H = \sqrt{V_{H,\infty}} I_3$, and $D_T = \sqrt{V_\infty} I_3$. Substituting these quantities into Equation (5.10) gives

$$\beta = \sqrt{V_\infty} \left[ \omega_G \sqrt{V_{G,\infty}} I_3 + \omega_H \sqrt{V_{H,\infty}} (\Sigma_{GG}^{(\mathrm{eq})})^{-1} \Sigma_{GH}^{(\mathrm{eq})} \right]^{-1} b. \tag{6.75}$$

This vector is on the standardized total-liability scale.

Finally, the raw total-liability trio covariance matrix is

$$\begin{aligned} \Sigma_{T^*T^*} &= (1-\pi) D_G \Sigma_{GG}^{(\mathrm{eq})} D_G + \sqrt{\pi(1-\pi)} \left( D_G \Sigma_{GH}^{(\mathrm{eq})} D_H + D_H (\Sigma_{GH}^{(\mathrm{eq})})^\top D_G \right) \\ &\quad + \pi D_H \Sigma_{HH}^{(\mathrm{eq})} D_H, \end{aligned} \tag{6.76}$$

and the standardized total-liability covariance matrix is

$$\Sigma_{TT} = D_T^{-1} \Sigma_{T^*T^*} D_T^{-1}. \tag{6.77}$$

### 7 Interpretation when the observed scores index a genetically correlated liability

The derivations above assume that the observed scores index the same latent liability that is substantively of interest for the phenotype. In practice, polygenic scores are often constructed for a genetically correlated trait rather than for the focal phenotype itself, for example using an educational attainment score to study child development. In that case, the fitted coefficients must be interpreted as projections through cross-trait genetic covariance rather than as direct coefficients on the focal trait's own liability.

Let the observed scores index a source trait $S$, with raw visible and hidden components $G^{(S),*}$ and $H^{(S),*}$, and raw total liability

$$T^{(S),*} = \sqrt{1-\pi_S}\, G^{(S),*} + \sqrt{\pi_S}\, H^{(S),*}.$$

Role-specific standardized copies are defined as in Equation (2.7). Let the focal phenotype be generated by a potentially different target-trait total liability vector $U^{(Y)}$ according to

$$Y = \left(\beta^{(Y)}\right)^\top U^{(Y)} + \gamma^\top W + \varepsilon_Y. \tag{7.1}$$

If one projects the phenotype on the score-indexed visible factors $G^{(S)}$, the population coefficient vector is

$$b^{(S\to Y)} = \left(\Sigma_{G^{(S)}G^{(S)}}\right)^{-1} \Sigma_{G^{(S)}U^{(Y)}}\, \beta^{(Y)}, \tag{7.2}$$

where $\Sigma_{G^{(S)}U^{(Y)}}$ is the cross-trait cross-role genetic covariance matrix. Equation (7.2) shows that the fitted coefficients are not direct effects of the source trait on the focal phenotype. Rather, they are coefficients on the component of the focal phenotype's liability that is tagged by the score-indexed source liability.

A useful special case arises when cross-trait genetic covariance is approximately proportional across all trio roles. Suppose

$$\Sigma_{G^{(S)}U^{(Y)}} = r_g^{(S,Y)} M_{SY}, \tag{7.3}$$

where $r_g^{(S,Y)}$ is a scalar cross-trait genetic correlation and $M_{SY}$ is a role-structured matrix encoding the cross-trait trio covariance structure whose $(r, r')$ entry gives the expected cross-trait genetic covariance between the source-trait visible liability for role $r$ and the target-trait liability entering the phenotype model for role $r'$, up to the common scalar factor $r_g^{(S,Y)}$. Then Equation (7.2) implies that the fitted coefficients scale linearly with the cross-trait genetic correlation. In particular, if the source and target liabilities coincide, then $r_g^{(S,Y)} = 1$ and Equation (7.2) reduces to the within-trait projection developed above. If $0 < r_g^{(S,Y)} < 1$, the coefficients should be interpreted as attenuated projections of target-liability effects onto the score-indexed liability.

Thus, when using a score for one trait to study another, the coefficients in the visible-scale SEM and the hidden variant correction quantify liability-correlated contributions tagged by the score, not a direct causal effect of the score-defining phenotype itself. The parental coefficients in such analyses retain their trio interpretation, but only for the component of the focal phenotype genetically aligned with the score-indexed trait.

### 8 Mendelian imputation of an unobserved parent

#### 8.1 Motivation

In many cohorts of practical interest, many mother–child duos are observed and the paternal genotype must be reconstructed by Mendelian imputation. Under the shared-weight construction of the visible-scale SEM, the imputed paternal polygenic score is not a noisy version of the unobserved paternal score with the same trio covariance structure. Instead, it is a specific linear combination of the observed child and maternal scores that carries a different relationship with every other latent and observed quantity in the trio. The prior trio covariance matrix $\Sigma_{GG}$ and the visible-scale projection coefficients must be modified accordingly. On the score-residual side, Mendelian imputation changes the paternal parent–offspring benchmark, but under the outbred benchmark it does not induce a nonzero mother–imputed-father residual correlation.

This section develops those modifications on the visible scale. Section 8.2 states the imputation scheme and establishes the central identity that the imputed paternal score equals the paternally inherited half of the child's raw visible component. Section 8.3 introduces the physical decomposition of the child's visible component into maternal and paternal half-scores and derives the role-standardized duo covariance structure. Sections 8.4–8.7 treat random mating, one generation of assortment, the exact $n$-generation raw-variance recursion, and equilibrium assortment, respectively. Section 8.8 derives the score-residual covariance structure implied by imputation. The total-liability correction for a duo must use the cross-covariance between the actual duo regressor vector and the true trio total liabilities; it is not obtained merely by substituting duo transmission coefficients into the complete-trio cross-block.

#### 8.2 Phased imputation and the imputed paternal score identity

Let locus $j$ have minor allele frequency $f_j$. For each family $i$ the mother and child are observed, and the father is not. Phased imputation of the father at locus $j$ proceeds in two steps. First, the child's two haplotypes are phased so that the paternally inherited allele count $h^{(\mathrm{p})}_{i\mathrm{c},j} \in \{0,1\}$ is identified. Second, the unobserved nontransmitted paternal allele is assigned its population expectation $f_j$. The imputed paternal genotype at locus $j$ is therefore

$$\hat{x}_{i\mathrm{p},j} = h^{(\mathrm{p})}_{i\mathrm{c},j} + f_j, \tag{8.1}$$

and the corresponding centered quantity is

$$\hat{x}_{i\mathrm{p},j} - 2f_j = h^{(\mathrm{p})}_{i\mathrm{c},j} - f_j = \tilde{h}^{(\mathrm{p})}_{i\mathrm{c},j}. \tag{8.2}$$

Thus the centered imputed paternal genotype at each locus is exactly the child's paternally inherited haplotype. The population-mean fill-in for the nontransmitted slot contributes zero after centering.

Define the child's raw paternal and maternal visible half-scores by

$$G^{(\mathrm{p}),*}_{i\mathrm{c}} = \sum_{j=1}^{M} \gamma_j \tilde{h}^{(\mathrm{p})}_{i\mathrm{c},j}, \qquad G^{(\mathrm{m}),*}_{i\mathrm{c}} = \sum_{j=1}^{M} \gamma_j \tilde{h}^{(\mathrm{m})}_{i\mathrm{c},j}, \tag{8.3}$$

so that $G^*_{i\mathrm{c}} = G^{(\mathrm{m}),*}_{i\mathrm{c}} + G^{(\mathrm{p}),*}_{i\mathrm{c}}$. Applying the shared GWAS-weight vector $w^{(k)}_j = \gamma_j + \xi^{(k)}_j$ uniformly across family members, the imputed paternal observed score is

$$\hat{\tilde{P}}^{(k)}_{i\mathrm{p}} = \sum_{j=1}^{M} w^{(k)}_j \tilde{h}^{(\mathrm{p})}_{i\mathrm{c},j}. \tag{8.4}$$

The corresponding latent-level identity, obtained by using the true effect-size vector $\gamma$ in place of $w^{(k)}$, is

$$\hat{G}^*_{i\mathrm{p}} = G^{(\mathrm{p}),*}_{i\mathrm{c}}. \tag{8.5}$$

This identity holds on the fixed raw scale by construction and does not require any distributional assumption on mating.

Let

$$\phi^* := \mathrm{Var}\left(G^{(\mathrm{p}),*}_{\mathrm{c}}\right) = \mathrm{Var}\left(G^{(\mathrm{m}),*}_{\mathrm{c}}\right), \tag{8.6}$$

where equality follows from symmetric Mendelian transmission. The standardized imputed paternal visible factor is

$$\hat{G}_{\mathrm{p}} = \frac{G^{(\mathrm{p}),*}_{\mathrm{c}}}{\sqrt{\phi^*}}, \qquad \mathrm{Var}(\hat{G}_{\mathrm{p}}) = 1. \tag{8.7}$$

#### 8.3 Physical decomposition of the offspring visible factor

Under additive Mendelian inheritance, the child's raw half-scores admit the physical decomposition

$$G^{(\mathrm{m}),*}_{\mathrm{c}} = \tfrac{1}{2}G^*_{\mathrm{m}} + U_{\mathrm{m}}, \qquad G^{(\mathrm{p}),*}_{\mathrm{c}} = \tfrac{1}{2}G^*_{\mathrm{p}} + U_{\mathrm{p}}, \tag{8.8}$$

where $U_{\mathrm{m}}$ and $U_{\mathrm{p}}$ are maternal and paternal Mendelian sampling deviations. Under the same infinitesimal constant-genic-variance approximation as Assumption 2,

$$\mathrm{Var}(U_{\mathrm{m}}) = \mathrm{Var}(U_{\mathrm{p}}) = \tfrac{1}{4}, \tag{8.9}$$

and the deviations are mutually uncorrelated and orthogonal to the parental visible components,

$$\mathrm{Cov}(U_{\mathrm{m}}, U_{\mathrm{p}}) = 0, \qquad \mathrm{Cov}(U_{\mathrm{m}}, G^*_{\mathrm{m}}) = \mathrm{Cov}(U_{\mathrm{m}}, G^*_{\mathrm{p}}) = \mathrm{Cov}(U_{\mathrm{p}}, G^*_{\mathrm{m}}) = \mathrm{Cov}(U_{\mathrm{p}}, G^*_{\mathrm{p}}) = 0. \tag{8.10}$$

Let

$$V := V_{G,\mathrm{par}}, \qquad K := K_{GG} = \mathrm{Cov}(G^*_{\mathrm{m}}, G^*_{\mathrm{p}}), \qquad V_{\mathrm{c}} := V_{G,\mathrm{c}}. \tag{8.11}$$

Equations (8.8)–(8.10) give

$$\phi^* = \frac{V+1}{4}, \tag{8.12}$$

$$\mathrm{Cov}\Big(G_{\mathrm{c}}^{(\mathrm{m}),*}, G_{\mathrm{c}}^{(\mathrm{p}),*}\Big) = \frac{K}{4}, \tag{8.13}$$

and therefore

$$V_{\mathrm{c}} = 2\phi^* + \frac{K}{2} = \frac{V+K+1}{2}. \tag{8.14}$$

The raw auxiliary covariances are

$$\mathrm{Cov}\Big(G_{\mathrm{c}}^{(\mathrm{p}),*}, G^*_{\mathrm{p}}\Big) = \frac{V}{2}, \tag{8.15}$$

$$\mathrm{Cov}\Big(G_{\mathrm{c}}^{(\mathrm{p}),*}, G^*_{\mathrm{m}}\Big) = \frac{K}{2}, \tag{8.16}$$

$$\mathrm{Cov}\Big(G_{\mathrm{c}}^{(\mathrm{p}),*}, G^*_{\mathrm{c}}\Big) = \phi^* + \frac{K}{4} = \frac{V_{\mathrm{c}}}{2}. \tag{8.17}$$

After standardizing $G^*_{\mathrm{c}}$, $G^*_{\mathrm{m}}$, and the imputed paternal half-score by their own role-specific raw standard deviations, the duo visible covariance matrix is

$$\Sigma^{\mathrm{duo}}_{GG}(V,K) = \begin{pmatrix} 1 & \dfrac{V+K}{2\sqrt{VV_{\mathrm{c}}}} & \sqrt{\dfrac{V_{\mathrm{c}}}{V+1}} \\ \dfrac{V+K}{2\sqrt{VV_{\mathrm{c}}}} & 1 & \dfrac{K}{\sqrt{V(V+1)}} \\ \sqrt{\dfrac{V_{\mathrm{c}}}{V+1}} & \dfrac{K}{\sqrt{V(V+1)}} & 1 \end{pmatrix}. \tag{8.18}$$

The correlation between the imputed paternal factor and the true standardized paternal factor is

$$\mathrm{Corr}(\hat{G}_{\mathrm{p}}, G_{\mathrm{p}}) = \sqrt{\frac{V}{V+1}}. \tag{8.19}$$

Solving the duo normal equations gives

$$t^{\mathrm{duo}}_{\mathrm{m}} = \frac{1}{2}\sqrt{\frac{V}{V_{\mathrm{c}}}}, \qquad t^{\mathrm{duo}}_{\mathrm{p}} = \frac{1}{2}\sqrt{\frac{V+1}{V_{\mathrm{c}}}}, \tag{8.20}$$

and the residual is the standardized maternal Mendelian sampling term,

$$\mathrm{Var}(u_G^{\mathrm{duo}}) = \frac{1}{4V_{\mathrm{c}}}. \tag{8.21}$$

The corresponding complete-trio residual variance is $1/(2V_{\mathrm{c}})$, so the duo residual variance is exactly one half as large.

#### 8.4 Duo covariance and transmission under no assortment

Under random mating, $V = V_{\mathrm{c}} = 1$ and $K = 0$. Equations (8.18)–(8.21) give

$$\mathrm{Cov}(\hat{G}_{\mathrm{p}}, G_{\mathrm{c}}) = \frac{1}{\sqrt{2}}, \tag{8.22}$$

$$\mathrm{Cov}(\hat{G}_{\mathrm{p}}, G_{\mathrm{m}}) = 0, \tag{8.23}$$

$$\mathrm{Corr}(\hat{G}_{\mathrm{p}}, G_{\mathrm{p}}) = \frac{1}{\sqrt{2}}. \tag{8.24}$$

The duo visible covariance matrix is therefore

$$\Sigma_{GG}^{\mathrm{duo,rm}} = \begin{pmatrix} 1 & 1/2 & 1/\sqrt{2} \\ 1/2 & 1 & 0 \\ 1/\sqrt{2} & 0 & 1 \end{pmatrix}. \tag{8.25}$$

The duo projection coefficients and residual variance are

$$t_{\mathrm{m}}^{\mathrm{duo}} = \tfrac{1}{2}, \qquad t_{\mathrm{p}}^{\mathrm{duo}} = \tfrac{1}{\sqrt{2}}, \tag{8.26}$$

$$\mathrm{Var}(u_G^{\mathrm{duo}}) = \tfrac{1}{4}. \tag{8.27}$$

The factor of two relative to the complete-trio residual reflects the fact that the imputed paternal score is a deterministic function of the child's paternally inherited alleles and therefore absorbs all paternal-side Mendelian sampling variance.

#### 8.5 Duo covariance and transmission under one generation of assortment from Hardy–Weinberg equilibrium

Consistent with the one-generation full-trio model, the parents are in the founder state with $V = 1$. The raw visible spouse covariance is

$$K = K_{GG} = (1 - \pi)\rho_T = \rho_G, \tag{8.28}$$

where the last equality holds because the parental visible variance is one. Hence

$$V_{\mathrm{c}} = 1 + \frac{\rho_G}{2}, \tag{8.29}$$

and $\phi^* = 1/2$. For the fitted SEM, $G^*_{\rm c}$ is divided by $\sqrt{V_{\rm c}}$ and the imputed paternal half-score is divided by $\sqrt{\phi^*}$. No raw causal component is rescaled before it is transmitted to a subsequent generation. If $\phi$ denotes the paternal half-score variance after expressing the half-score on the standardized child scale, then

$$\phi = \frac{\phi^*}{V_{\rm c}} = \frac{1}{2+\rho_G}. \tag{8.30}$$

The one-generation duo covariance matrix is

$$\Sigma^{\rm duo,1gen}_{GG} = \begin{pmatrix} 1 & (1+\rho_G)/\sqrt{2(2+\rho_G)} & \sqrt{2+\rho_G}/2 \\ (1+\rho_G)/\sqrt{2(2+\rho_G)} & 1 & \rho_G/\sqrt{2} \\ \sqrt{2+\rho_G}/2 & \rho_G/\sqrt{2} & 1 \end{pmatrix}. \tag{8.31}$$

In particular,

$$\mathrm{Cov}(\hat{G}_{\rm p}, G_{\rm c}) = \frac{\sqrt{2+\rho_G}}{2}, \tag{8.32}$$

$$\mathrm{Cov}(\hat{G}_{\rm p}, G_{\rm m}) = \frac{\rho_G}{\sqrt{2}}, \tag{8.33}$$

$$\mathrm{Corr}(\hat{G}_{\rm p}, G_{\rm p}) = \frac{1}{\sqrt{2}}. \tag{8.34}$$

The projection coefficients are

$$t^{\rm duo}_{\rm m} = \frac{1}{\sqrt{2(2+\rho_G)}}, \qquad t^{\rm duo}_{\rm p} = \frac{1}{\sqrt{2+\rho_G}}, \tag{8.35}$$

and

$$\mathrm{Var}(u^{\rm duo}_G) = \frac{1}{2(2+\rho_G)}. \tag{8.36}$$

The departure of the standardized coefficients from one half is induced by the role-specific standardization of a transient child variance, not by a change in the raw Mendelian inheritance coefficients.

#### 8.6 Exact $n$-generation recursion

When assortment has acted for several generations, no separate restandardized half-score recursion is required. The raw visible variance follows directly from Section 6.2. For adults in generation $g$,

$$V = V_{G,g} = \pi + (1-\pi)V_g, \qquad K = K_{GG,g} = \rho_{T,g}(1-\pi)V_g, \tag{8.37}$$

and their offspring have

$$V_{G,g+1} = \frac{V_{G,g} + K_{GG,g} + 1}{2} = \pi + (1-\pi)V_{g+1}. \tag{8.38}$$

The raw paternal half-score variance for that offspring is

$$\phi^*_{g+1} = \frac{V_{G,g}+1}{4}. \tag{8.39}$$

Substitution of $V = V_{G,g}$, $K = K_{GG,g}$, and $V_c = V_{G,g+1}$ into Equations (8.18)–(8.21) gives the exact $n$-generation duo covariance and projection coefficients. For constant $\rho_T$, $V_g$ is given by Equation (6.22); for a varying history it is given by Equation (6.53).

#### 8.7 Duo covariance and transmission at equilibrium

At equilibrium, $V_{G,\mathrm{c}} = V_{G,\mathrm{par}} =: V$ and Equation (8.38) implies $K = V - 1$. Therefore the standardized visible spouse correlation is

$$\rho_G = \frac{K}{V} = 1 - \frac{1}{V}, \qquad V = \frac{1}{1-\rho_G}. \tag{8.40}$$

The raw paternal half-score variance is $\phi^* = (V+1)/4$. Expressed relative to the standardized child visible factor, it is

$$\phi = \frac{\phi^*}{V} = \frac{2-\rho_G}{4}. \tag{8.41}$$

The equilibrium offspring–parent covariance in the observed-trio SEM is

$$\eta_{\mathrm{m}} = \eta_{\mathrm{p}} = \frac{1+\rho_G}{2}, \tag{8.42}$$

and the equilibrium duo-specific covariances are

$$\mathrm{Cov}(\hat{G}_{\mathrm{p}}, G_{\mathrm{c}}) = \frac{1}{\sqrt{2-\rho_G}}, \tag{8.43}$$

$$\mathrm{Cov}(\hat{G}_{\mathrm{p}}, G_{\mathrm{m}}) = \frac{\rho_G}{\sqrt{2-\rho_G}}, \tag{8.44}$$

$$\mathrm{Corr}(\hat{G}_{\mathrm{p}}, G_{\mathrm{p}}) = \frac{1}{\sqrt{2-\rho_G}}. \tag{8.45}$$

The equilibrium duo visible covariance matrix is therefore

$$\Sigma_{GG}^{\mathrm{duo,eq}} = \begin{pmatrix} 1 & (1+\rho_G)/2 & 1/\sqrt{2-\rho_G} \\ (1+\rho_G)/2 & 1 & \rho_G/\sqrt{2-\rho_G} \\ 1/\sqrt{2-\rho_G} & \rho_G/\sqrt{2-\rho_G} & 1 \end{pmatrix}. \tag{8.46}$$

The duo projection coefficients and residual variance are

$$t_{\mathrm{m}}^{\mathrm{duo}} = \tfrac{1}{2}, \qquad t_{\mathrm{p}}^{\mathrm{duo}} = \frac{\sqrt{2-\rho_G}}{2}, \tag{8.47}$$

$$\mathrm{Var}(u_G^{\mathrm{duo}}) = \frac{1-\rho_G}{4}, \tag{8.48}$$

which is one half of the corresponding complete-trio residual variance $(1-\rho_G)/2$. The improvement in latent imputation quality relative to random mating follows from the raw within-individual disequilibrium accumulated at equilibrium; it does not arise from restandardizing the causal component between generations.

#### 8.8 Implied score-residual covariance structure

Because Equation (8.5) is a latent-level identity, the same derivation applies to the observed scores after substitution of the noisy weight vector $w^{(k)} = \gamma + \xi^{(k)}$. The imputed paternal observed score is

$$\widetilde{\hat{P}}_{\mathrm{p}}^{(k)} \;=\; \sum_{j=1}^{M} \bigl(\gamma_j + \xi_j^{(k)}\bigr) \tilde{h}_{\mathrm{c},j}^{(\mathrm{p})} \;=\; G_{\mathrm{c}}^{(\mathrm{p})} + \widehat{\tilde{\varepsilon}}_{\mathrm{p}}^{(k)}, \qquad \widehat{\tilde{\varepsilon}}_{\mathrm{p}}^{(k)} \;=\; \sum_{j=1}^{M} \xi_j^{(k)} \tilde{h}_{\mathrm{c},j}^{(\mathrm{p})}. \tag{8.49}$$

The imputed raw-score residual contribution $\widehat{\tilde{\varepsilon}}_{\mathrm{p}}^{(k)}$ is constructed from the same weight-noise vector $\xi^{(k)}$ as the child's and mother's raw-score residual contributions $\tilde{\varepsilon}_{\mathrm{c}}^{(k)}, \tilde{\varepsilon}_{\mathrm{m}}^{(k)}$ of Equation (4.14), but weighted by the child's paternally inherited haplotype rather than by the father's genotype.

For the imputation-specific benchmark below, the score-error component is additionally taken to be the fitted residual direction after projection off the visible liability, orthogonal to the total liability on which mating occurs, and free of cross-mate residual covariance. Under this zero-spouse-residual benchmark, the maternally and paternally transmitted error half-scores are uncorrelated. The following $1/\sqrt{2}$ result is conditional on these assumptions; independence of the raw GWAS weight perturbations from the true effect sizes alone is not sufficient if the realized residual score direction also participates in mating.

Let $\tilde{\varepsilon}_{\mathrm{c}}^{(\mathrm{p},k)} = \widehat{\tilde{\varepsilon}}_{\mathrm{p}}^{(k)}$ denote the paternally inherited error half-score, and define the maternal error half-score $\tilde{\varepsilon}_{\mathrm{c}}^{(\mathrm{m},k)}$ analogously. Then

$$\tilde{\varepsilon}_{\mathrm{c}}^{(k)} = \tilde{\varepsilon}_{\mathrm{c}}^{(\mathrm{m},k)} + \tilde{\varepsilon}_{\mathrm{c}}^{(\mathrm{p},k)}. \tag{8.50}$$

Under symmetric transmission and the zero-spouse-residual benchmark, write

$$\mathrm{Var}\Bigl(\tilde{\varepsilon}_{\mathrm{c}}^{(\mathrm{m},k)}\Bigr) = \mathrm{Var}\Bigl(\tilde{\varepsilon}_{\mathrm{c}}^{(\mathrm{p},k)}\Bigr) = \tau_k^2, \qquad \mathrm{Cov}\Bigl(\tilde{\varepsilon}_{\mathrm{c}}^{(\mathrm{m},k)}, \tilde{\varepsilon}_{\mathrm{c}}^{(\mathrm{p},k)}\Bigr) = 0. \tag{8.51}$$

It follows that

$$\mathrm{Cov}\Bigl(\widehat{\tilde{\varepsilon}}_{\mathrm{p}}^{(k)}, \tilde{\varepsilon}_{\mathrm{c}}^{(k)}\Bigr) = \tau_k^2, \qquad \mathrm{Var}\Bigl(\tilde{\varepsilon}_{\mathrm{c}}^{(k)}\Bigr) = 2\tau_k^2, \tag{8.52}$$

and therefore, after standardization of the two residual scores,

$$\mathrm{Corr}\Bigl(\hat{\varepsilon}_{\mathrm{p}}^{(k)}, \varepsilon_{\mathrm{c}}^{(k)}\Bigr) = \frac{1}{\sqrt{2}}. \tag{8.53}$$

This is the imputation-specific paternal parent–offspring benchmark. It is unchanged across the random-mating, one-generation, and equilibrium regimes only under the additional zero-spouse-residual conditions stated above.

For the mother and the imputed father, the same benchmark gives

$$\mathrm{Cov}\Bigl(\widehat{\tilde{\varepsilon}}_{\mathrm{p}}^{(k)}, \tilde{\varepsilon}_{\mathrm{m}}^{(k)}\Bigr) = 0, \qquad \mathrm{Corr}\Bigl(\hat{\varepsilon}_{\mathrm{p}}^{(k)}, \varepsilon_{\mathrm{m}}^{(k)}\Bigr) = 0. \tag{8.54}$$

The zero follows from the imposed absence of residual-score mate covariance and recent biological relatedness between the mother and the paternal transmitted haplotype; it is not implied by Mendelian imputation alone.

Thus Mendelian imputation changes the paternal parent–offspring residual benchmark from the real-trio value of one half in Equation (4.19) to $1/\sqrt{2}$, because the imputed paternal score is built from a single transmitted haplotype rather than a full diploid genotype. By contrast, Mendelian imputation does not by itself induce a nonzero mother–imputed-father residual correlation under the benchmark assumptions used in this note. If one wishes to allow consanguinity, endogamy, or other departures from the outbred benchmark, the corresponding maternal–imputed-father residual covariance should be estimated rather than fixed.

#### 8.9 Summary of duo-specific quantities

Table 2 collects the duo-specific visible-scale quantities across the three assortment regimes, alongside the corresponding real-trio values for reference. All quantities are written in the standardization convention, $\mathrm{Var}(G_\mathrm{m}) = \mathrm{Var}(G_\mathrm{p}) = \mathrm{Var}(G_\mathrm{c}) = 1$.

| Quantity | Random mating | One-generation | Equilibrium |
|---|---|---|---|
| $\mathrm{Cov}(\hat{G}_\mathrm{p}, G_\mathrm{c})$ | $1/\sqrt{2}$ | $\sqrt{2+\rho_G}/2$ | $1/\sqrt{2-\rho_G}$ |
| $\mathrm{Cov}(\hat{G}_\mathrm{p}, G_\mathrm{m})$ | 0 | $\rho_G/\sqrt{2}$ | $\rho_G/\sqrt{2-\rho_G}$ |
| $\mathrm{Corr}(\hat{G}_\mathrm{p}, G_\mathrm{p})$ | $1/\sqrt{2}$ | $1/\sqrt{2}$ | $1/\sqrt{2-\rho_G}$ |
| $t_\mathrm{m}^\mathrm{duo}$ | $1/2$ | $1/\sqrt{2(2+\rho_G)}$ | $1/2$ |
| $t_\mathrm{p}^\mathrm{duo}$ | $1/\sqrt{2}$ | $1/\sqrt{2+\rho_G}$ | $\sqrt{2-\rho_G}/2$ |
| $\mathrm{Var}(u_G^\mathrm{duo})$ | $1/4$ | $1/(2(2+\rho_G))$ | $(1-\rho_G)/4$ |
| Real-trio $\mathrm{Var}(u_G)$ (reference) | $1/2$ | $1/(2+\rho_G)$ | $(1-\rho_G)/2$ |
| $\mathrm{Corr}(\hat{\varepsilon}_\mathrm{p}^{(k)}, \varepsilon_\mathrm{c}^{(k)})$ | $1/\sqrt{2}$ | $1/\sqrt{2}$ | $1/\sqrt{2}$ |
| $\mathrm{Corr}(\hat{\varepsilon}_\mathrm{p}^{(k)}, \varepsilon_\mathrm{m}^{(k)})$ | 0 | 0 | 0 |

Table 2: Duo-specific visible-scale quantities under phased paternal imputation, for three assortment regimes. The score-residual rows additionally impose the zero-spouse-residual benchmark stated in Section 8.8. The real-trio Mendelian residual variance is included for reference. In every regime the duo Mendelian residual variance is exactly one half of the real-trio value. The offspring–imputed-paternal covariance exceeds the real-trio offspring–parent covariance in every regime because the imputed paternal score is constructed from the child's own paternally-inherited haplotype.

The quantities in Table 2 determine the visible-scale covariance block for an imputed-parent group. The total liability mapping must, however, reflect that the third visible regressor is the child's paternally inherited half-score whereas the third target liability is the true paternal total liability. Let

$$X^\mathrm{duo} = (G_\mathrm{c}, G_\mathrm{m}, \hat{G}_\mathrm{p})^\top.$$

Then the relevant population map is

$$b^\mathrm{duo} = \left(\Sigma_{XX}^\mathrm{duo}\right)^{-1} \Sigma_{XT}^\mathrm{duo}\,\beta, \qquad \Sigma_{XT}^\mathrm{duo} = \mathrm{Cov}(X^\mathrm{duo}, T), \tag{8.55}$$

and, when the square map is nonsingular,

$$\beta = \left[\left(\Sigma_{XX}^\mathrm{duo}\right)^{-1} \Sigma_{XT}^\mathrm{duo}\right]^{-1} b^\mathrm{duo}. \tag{8.56}$$

The matrix $\Sigma_{XT}^{\mathrm{duo}}$ is constructed from the raw component moments in Section 6.2 together with the imputation identity in Equation (8.5).

### 9 Sensitivity to imperfect genetic correlation between score versions

The preceding derivations assume that the two observed score versions for a given role are noisy indicators of the same visible-scale latent genetic component. This assumption is exact only if the genetic components indexed by the two score versions are perfectly correlated, apart from score-construction error. In practice, two independently trained polygenic scores for the same nominal trait may have genetic correlation slightly below one. This section gives a simple sensitivity calculation for the resulting bias when score-version-specific genetic signal also covaries with the phenotype.

For a generic role $r \in \{\mathrm{c}, \mathrm{m}, \mathrm{p}\}$, suppose the standardized observed score version $k \in \{1, 2\}$ can be written in the one-factor measurement form

$$P_r^{(k)} = \sqrt{R_{kr} r_g}\, G_r + \varepsilon_r^{(k)}, \tag{9.1}$$

where $G_r$ is the intended shared visible-scale latent component, $r_g$ is the genetic correlation between the error-free genetic components indexed by the two score versions, and $R_{kr}$ is the fraction of variance in $P_r^{(k)}$ attributable to its error-free genetic component. The score residual $\varepsilon_r^{(k)}$ is the same residual term introduced in Equations (4.1)–(4.2). Here we refine it as

$$\varepsilon_r^{(k)} = \sqrt{R_{kr}(1 - r_g)}\, \Omega_r^{(k)} + \varepsilon_r^{(k),0}, \tag{9.2}$$

where $\Omega_r^{(k)}$ is a score-version-specific genetic component and $\varepsilon_r^{(k),0}$ is the classical score-construction component of the residual. We assume

$$\mathrm{Var}(G_r) = \mathrm{Var}(\Omega_r^{(k)}) = 1, \qquad \mathrm{Var}(\varepsilon_r^{(k),0}) = 1 - R_{kr}, \tag{9.3}$$

and

$$\mathrm{Cov}(G_r, \Omega_r^{(1)}) = 0, \qquad \mathrm{Cov}(G_r, \Omega_r^{(2)}) = 0, \qquad \mathrm{Cov}(\Omega_r^{(1)}, \Omega_r^{(2)}) = 0, \tag{9.4}$$

with

$$\mathrm{Cov}(\varepsilon_r^{(k),0}, G_r) = 0, \qquad \mathrm{Cov}(\varepsilon_r^{(k),0}, \Omega_r^{(1)}) = 0, \qquad \mathrm{Cov}(\varepsilon_r^{(k),0}, \Omega_r^{(2)}) = 0, \qquad \mathrm{Cov}(\varepsilon_r^{(k),0}, Y) = 0. \tag{9.5}$$

Thus the error-free genetic score components can be written as

$$P_r^{(1),g} = \sqrt{r_g}\, G_r + \sqrt{1 - r_g}\, \Omega_r^{(1)}, \qquad P_r^{(2),g} = \sqrt{r_g}\, G_r + \sqrt{1 - r_g}\, \Omega_r^{(2)}. \tag{9.6}$$

The loading of score version $k$ on the intended shared component is therefore

$$\lambda_{kr} = \sqrt{R_{kr} r_g}. \tag{9.7}$$

Let

$$\mathcal{C}_r := \mathrm{Cov}(G_r, Y), \qquad \mathcal{O}_{kr} := \mathrm{Cov}(\Omega_r^{(k)}, Y). \tag{9.8}$$

Then

$$\begin{aligned}\mathrm{Cov}(P_r^{(k)}, Y) &= \sqrt{R_{kr} r_g}\,\mathrm{Cov}(G_r, Y) + \sqrt{R_{kr}(1-r_g)}\,\mathrm{Cov}(\Omega_r^{(k)}, Y) \\ &= \sqrt{R_{kr} r_g}\,\mathcal{C}_r + \sqrt{R_{kr}(1-r_g)}\,\mathcal{O}_{kr}. \end{aligned} \tag{9.9}$$

The one-factor measurement model treats the score residual as unrelated to $Y$. Therefore, when the score-version-specific genetic component has nonzero phenotype covariance, the latent-scale covariance inferred from score version $k$ is

$$\begin{aligned}\widehat{\mathcal{C}}_{kr} &= \frac{\mathrm{Cov}(P_r^{(k)}, Y)}{\lambda_{kr}} \\ &= \mathcal{C}_r + \sqrt{\frac{1-r_g}{r_g}}\,\mathcal{O}_{kr}. \end{aligned} \tag{9.10}$$

A two-indicator fit with symmetric information averages the two score-specific contributions, giving

$$\widehat{\mathcal{C}_r} = \mathcal{C}_r + \sqrt{\frac{1-r_g}{r_g}}\,\overline{\mathcal{O}}_r, \qquad \overline{\mathcal{O}}_r = \frac{\mathcal{O}_{1r} + \mathcal{O}_{2r}}{2}. \tag{9.11}$$

Equation (9.11) is the general role-wise bias expression. The classical component $\varepsilon_r^{(k),0}$ is absorbed by the measurement model through the loading and residual variance; the bias arises only because the score-version-specific genetic component is treated as residual variation while covarying with $Y$.

To express the magnitude in terms of $r_g$, write the unstandardized error-free genetic score components as

$$P_r^{(k),g} = Q_r + \Omega_r^{(k),\mathrm{raw}}, \tag{9.12}$$

with

$$\mathrm{Var}(Q_r) = A, \qquad \mathrm{Var}(\Omega_r^{(k),\mathrm{raw}}) = U, \tag{9.13}$$

and

$$\mathrm{Cov}(Q_r, \Omega_r^{(k),\mathrm{raw}}) = 0, \qquad \mathrm{Cov}(\Omega_r^{(1),\mathrm{raw}}, \Omega_r^{(2),\mathrm{raw}}) = 0. \tag{9.14}$$

Then

$$r_g = \frac{A}{A+U}, \qquad \frac{U}{A} = \frac{1-r_g}{r_g}. \tag{9.15}$$

Suppose the raw phenotype covariance of the score-version-specific component is a fraction $\tau_\Omega$ of the raw phenotype covariance of the shared component:

$$\mathrm{Cov}(Q_r, Y) = \alpha_r A, \qquad \mathrm{Cov}(\Omega_r^{(k),\mathrm{raw}}, Y) = \tau_\Omega\,\alpha_r U. \tag{9.16}$$

Here $\tau_\Omega = 0$ corresponds to score-version-specific genetic signal that is uncorrelated with the phenotype, whereas $\tau_\Omega = 1$ corresponds to equal raw phenotype association for the shared and score-version-specific genetic components. Substituting Equation (9.16) into Equation (9.11) gives

$$\frac{\widehat{\mathcal{C}}_r}{\mathcal{C}_r} = 1 + \tau_\Omega \frac{U}{A} = 1 + \tau_\Omega \frac{1 - r_g}{r_g}. \tag{9.17}$$

Thus the visible-scale covariance $\mathrm{Cov}(G_r, Y)$ is unbiased when $\tau_\Omega = 0$, and under equal raw effects ($\tau_\Omega = 1$) is inflated by

$$\frac{\widehat{\mathcal{C}}_r}{\mathcal{C}_r} = \frac{1}{r_g}. \tag{9.18}$$

The natural sensitivity range from "score-version-specific component uncorrelated with $Y$" to "equal raw effect on $Y$" is therefore

$$0 \ \text{ to } \ \frac{1}{r_g} - 1 \tag{9.19}$$

as a proportional inflation of the visible-scale covariance, assuming the score-version-specific component has the same direction of association as the shared component. If $\mathcal{C}_r = 0$, the ratio in Equation (9.17) is not defined; under the raw-effect model in Equation (9.16), however, the absolute bias is also zero.

The same conclusion applies to the visible-scale coefficient vector when the same $r_g$ and raw-effect ratio $\tau_\Omega$ apply across the offspring, maternal, and paternal score systems. In that case, the covariance vector $\mathrm{Cov}(G, Y)$ is multiplied by a common scalar factor, and therefore the visible-scale coefficient vector satisfies

$$\widehat{b} = F(r_g, \tau_\Omega)\, b, \qquad F(r_g, \tau_\Omega) = 1 + \tau_\Omega \frac{1 - r_g}{r_g}. \tag{9.20}$$

Finally, the visible-to-total transformation in Equation (5.10) is linear in the visible coefficient vector for a fixed raw AM state. Writing

$$M_\pi = \left[\omega_G D_G + \omega_H \Sigma_{GG}^{-1} \Sigma_{GH} D_H\right] D_T^{-1}, \tag{9.21}$$

the standardized total-liability correction is

$$\beta = M_\pi^{-1} b. \tag{9.22}$$

Therefore, if the score-version-specific genetic component multiplies the visible coefficient vector by the scalar factor $F(r_g, \tau_\Omega)$, then

$$\widehat{\beta} = M_\pi^{-1} \widehat{b} = M_\pi^{-1} F(r_g, \tau_\Omega) b = F(r_g, \tau_\Omega) \beta. \tag{9.23}$$

The total-liability transformation therefore does not introduce an additional $\pi$-dependent percentage bias. It carries forward the same multiplicative factor already present on the visible scale. Under equal raw effects, $\tau_\Omega = 1$, both the visible-scale coefficients and the total-liability coefficients are inflated by the factor $1/r_g$; under $\tau_\Omega = 0$, neither is inflated by this mechanism.

### 10 Simulation framework

The simulation framework is designed to mirror the conceptual structure of the fitted model. It generates visible and hidden genetic variation separately on a fixed founder scale, combines them into raw total liabilities, imposes assortative mating on the total liabilities, constructs observed polygenic scores from visible variants only, and generates phenotypes from the total trio liabilities.

#### 10.1 Visible and hidden causal variants

Let there be $M$ visible causal variants with effect-size vector

$$\gamma^{\mathrm{vis}} = \left(\gamma_1^{\mathrm{vis}}, \ldots, \gamma_M^{\mathrm{vis}}\right)^\top, \qquad \gamma_j^{\mathrm{vis}} \sim \mathcal{N}(0,1), \tag{10.1}$$

and, when hidden variation is present, $M_{\mathrm{hid}}$ hidden causal variants with effect-size vector

$$\gamma^{\mathrm{hid}} = \left(\gamma_1^{\mathrm{hid}}, \ldots, \gamma_{M_{\mathrm{hid}}}^{\mathrm{hid}}\right)^\top, \qquad \gamma_j^{\mathrm{hid}} \sim \mathcal{N}(0,1). \tag{10.2}$$

Parental visible and hidden genotypes are sampled independently across loci under specified allele frequencies. Let $\mu_{G,0}, s_{G,0}$ and $\mu_{H,0}, s_{H,0}$ be the means and standard deviations of the founder weighted sums. The raw founder-scaled liabilities are

$$G_{ir}^* = \frac{\sum_{j=1}^{M} X_{ijr}^{\mathrm{vis}} \gamma_j^{\mathrm{vis}} - \mu_{G,0}}{s_{G,0}}, \qquad H_{ir}^* = \frac{\sum_{j=1}^{M_{\mathrm{hid}}} X_{ijr}^{\mathrm{hid}} \gamma_j^{\mathrm{hid}} - \mu_{H,0}}{s_{H,0}}, \qquad r \in \{\mathrm{m}, \mathrm{p}\}. \tag{10.3}$$

The same founder constants are retained in every later generation. The raw total parental liabilities are

$$T_{ir}^* = \omega_G G_{ir}^* + \omega_H H_{ir}^*. \tag{10.4}$$

#### 10.2 Assortative mating imposed on the total liability

Assortative mating is imposed directly on the parental raw total liabilities. Let $T_{i\mathrm{m}}^*$ and $T_{i\mathrm{p}}^*$ denote maternal and paternal values before pairing. Draw latent Gaussian pairs

$$Z_i = (Z_{i1}, Z_{i2})^\top \sim \mathcal{N}\left(0, \begin{pmatrix} 1 & \rho_T \\ \rho_T & 1 \end{pmatrix}\right), \qquad i = 1, \ldots, N, \tag{10.5}$$

where $\rho_T$ is the target spouse correlation on the standardized total-liability scale. Let $\phi_{\mathrm{m}}$ be the ordering of mothers induced by $T_{i\mathrm{m}}^*$ and $\phi_{\mathrm{p}}$ the ordering of fathers induced by $T_{i\mathrm{p}}^*$. Let $r_{\mathrm{m}}(i)$ and $r_{\mathrm{p}}(i)$ denote the ranks of $Z_{i1}$ and $Z_{i2}$ among the $N$ Gaussian draws. Mothers and fathers are then matched according to

$$\mathrm{mother}_i = \phi_{\mathrm{m}}\big(r_{\mathrm{m}}(i)\big), \qquad \mathrm{father}_i = \phi_{\mathrm{p}}\big(r_{\mathrm{p}}(i)\big). \tag{10.6}$$

This Gaussian-copula rank-matching procedure preserves the marginal distributions of the parental total liabilities while inducing a spousal correlation close to the target $\rho_T$. Because mating is imposed on the total liabilities, visible and hidden components become correlated through the mating step.

The analytic derivations additionally use Assumption 3. For jointly Gaussian component liabilities, mating through $T^*$ gives the rank-one component covariance in that assumption exactly; for finite polygenic simulations, the realized component moments are checked against that target.

##### 10.3 Observed polygenic scores with controlled reliability

Two observed polygenic scores are generated from the visible variants only. If score version $k \in \{1, 2\}$ is intended to have founder-scale reliability $R_k$, the corresponding weight-noise standard deviation is set to

$$\sigma_{w_k} = \sqrt{\frac{1 - R_k}{R_k}}. \tag{10.7}$$

The noisy weight vector is

$$w_j^{(k)} = \gamma_j^{\text{vis}} + \xi_j^{(k)}, \qquad \xi_j^{(k)} \sim \mathcal{N}(0, \sigma_{w_k}^2), \tag{10.8}$$

and the observed score used in the fitted SEM is the standardized weighted sum

$$P_{ir}^{(k)} = \text{scale}\left(\sum_{j=1}^{M} X_{ijr}^{\text{vis}} w_j^{(k)}\right), \tag{10.9}$$

using the observed-score reference convention chosen for the analysis (for example, a common child reference scale). The same noisy weight vector $w^{(k)}$ is applied to mother, father, and child. This shared-weight construction is the source of the cross-person score-residual covariance estimated by the SEM. Standardization of the observed indicators for fitting does not alter the raw causal-score propagation in Equation (10.3).

##### 10.4 Mendelian transmission

The offspring genotype is created by Mendelian transmission. At each locus, one allele is transmitted from each parent. Conditional on parental genotype $g \in \{0, 1, 2\}$, the transmitted allele count is sampled as

$$X^{\text{trans}} \sim \text{Binomial}(1, g/2). \tag{10.10}$$

This is carried out separately for visible and hidden variants. The offspring raw visible and hidden liabilities are constructed using the same effect vectors and the same founder means and standard deviations as in Equation (10.3); they are not separately rescaled before a further mating round. The offspring raw total liability is

$$T_{ic}^* = \omega_G G_{ic}^* + \omega_H H_{ic}^*. \tag{10.11}$$

For the focal SEM, role-standardized versions of $G_{ir}$, $H_{ir}$, and $T_{ir}$ are then formed according to Equation (2.7). Because visible and hidden variants are inherited through the same additive mechanism, the raw common-transmission assumption used in the correction is aligned with the simulation design.

##### 10.5 Phenotype generation

We then generate the phenotype from the founder-scaled raw total trio liabilities:

$$Y_i = (\beta_D^{\text{raw}})T_{ic}^* + (\beta_M^{\text{raw}})T_{im}^* + (\beta_P^{\text{raw}})T_{ip}^* + \varepsilon_{Yi}. \tag{10.12}$$

Thus the generating coefficients in the simulator are coefficients on the raw total-liability scale. The corresponding coefficients on the role-standardized total-liability scale are

$$\beta = D_T \beta^{\text{raw}}. \tag{10.13}$$

The implementation calculates the realized variance of the joint genetic component and chooses the phenotype residual variance so that the outcome variance is approximately one, subject to a small positive residual-variance floor.

##### 10.6 Validation targets under simulation

The simulation framework yields several distinct targets against which the fitted model can be assessed. First, because the raw total liability coefficients are known by construction, the corrected raw coefficient vector in Equation (5.11) can be compared directly with the generating vector $\beta^{\text{raw}}$. Equivalently, the standardized corrected vector in Equation (5.10) can be compared with the transformed target $D_T\beta^{\text{raw}}$ from Equation (10.13). Second, because the simulation also generates the raw visible and hidden components separately, one can compare the fitted visible-scale coefficients to the regression coefficients obtained by projecting the simulated phenotype on the known role-standardized visible liabilities. Third, one can compare the attenuation-only benchmark in Equation (5.12) to the full assortment correction to quantify the impact of the structured visible–hidden covariance terms. Finally, the simulated raw component variances and covariances can be compared directly with Equations (6.23)–(6.25). These validation targets ensure that the fitted SEM, the raw variance recursion, the hidden variant correction, and the interpretation of coefficient scale can all be checked separately.
